# Octopamine instructs head direction plasticity

**DOI:** 10.64898/2025.12.11.693783

**Authors:** Mark H. Plitt, Daniel B. Turner-Evans, Jessica C. Co, Aryanna Layden, Mark Eddison, Robert P. Ray, Vivek Jayaraman, Yvette E. Fisher

## Abstract

Many plasticity rules rely on adjusting the strength of synapses between pairs of cells based on their coincident activity. We uncovered a new mechanism for coincidence detection in the *Drosophila* head direction network. To maintain an accurate sense of direction, head direction neurons that signal orientation during navigation must learn to anchor to relevant external sensory cues in novel environments. Yet the synaptic mechanism for this form of unsupervised learning is unknown in any organism. In *Drosophila*, GABAergic visual inputs converge onto head direction neurons, and these inhibitory synapses change strength with experience to learn the relationship between visual landmarks and head direction. However, how coincident pre– and postsynaptic activity is detected across this inhibitory synapse is not understood. We discovered that neurons which release the monoamine octopamine close a feedback loop that conveys postsynaptic head direction activity onto presynaptic terminals of visual inputs. This octopamine pathway is required for anchoring the head direction network to visual cues. Furthermore, pairing structured activation of octopamine neurons with a visual cue is sufficient to drive rapid plasticity, even without postsynaptic head direction cell activity. Previous work has extensively characterized coincidence detection mechanisms at excitatory synapses; our work defines a novel mechanism for coincidence detection at an inhibitory synapse, in which postsynaptic activity is relayed via a neuromodulatory neuron onto presynaptic terminals.

## Main text

The discovery of head direction cells in brains across the animal kingdom underscores their importance for adaptive navigation^1–5^. Each head direction cell fires most strongly when an animal faces in that cell’s preferred direction. As a population, head direction cells tile all angles such that their collective activity (referred to as the head direction representation) resembles a compass, estimating the animal’s orientation in space by combining self-motion inputs with external sensory cues (Fig. 1a-b, Extended Data Fig. 1a).

**Figure 1.**
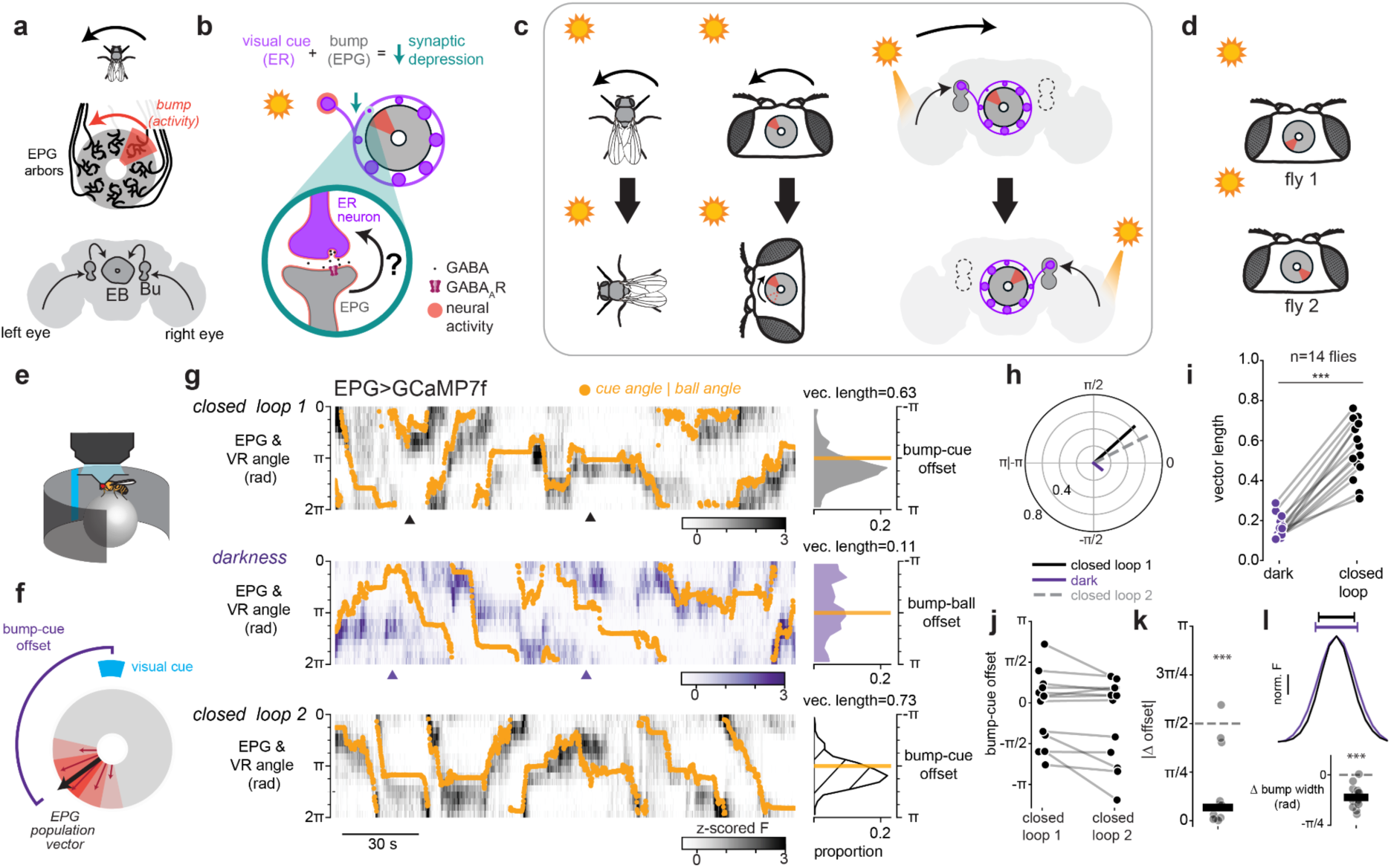
The head direction network stores its relationship to visual cues. a) *Top:* EPG neuron arbors innervate wedges around the EB that encode the fly’s head direction (HD) angle. *Bottom:* Visual information is relayed from the eye through the Bulb (Bu) and into the ellipsoid body (EB). b) Visual information is carried by GABAergic ER neurons (magenta) with synapses onto every EPG wedge (gray). To associate visual cues with changes in head direction, synapses between coactive ER and EPG neurons are thought to undergo depression (teal). Such a learning rule requires coincidence detection at this GABAergic synapse which could occur if active EPGs send an unknown retrograde signal to ER axons. c) Schematic of how ER-EPG plasticity enables the bump of activity in EPGs to accurately track visual cues. As a fly makes a counter-clockwise turn (top to bottom) it will view visual cues (e.g. the sun) from a new angle and the EPG activity bump (red) will swing clockwise around the network by integrating self motion signals with these visual inputs. When the fly faces a different angle, distinct visual ER neurons are active. Plasticity forms a trough of weak synapses (large circles – strong synapses, small circles – weak synapses) that allow ER neurons with distinct visual tuning to move the EPG bump via disinhibition. d) The relationship (offset) between a visual cue angle and the angle of EPG activity bump is arbitrary across flies and determined by ER-EPG plasticity. e) Neural activity is monitored using a 2P microscope while a head-fixed fly walks on an air-cushioned ball in a VR arena. The angle of a blue bar serves as an orienting landmark which tracks the ball yaw angle. f) EPG head direction activity is calculated as the population vector average (PVA) from all ellipsoid body angles weighted by their fluorescence intensity. Bump-cue offset is calculated as the difference between the PVA angle in the brain and the cue angle in VR (the ball angle in dark sessions). g) *Left:* Example EPG imaging data from a single fly that first walked with an orienting visual cue (closed loop 1, top), then in darkness (middle), then with a visual cue again (closed loop 2, bottom). Heatmaps display EPG activity unwrapped by ellipsoid body angle from the first 3 minutes of each 6 minute session. Orange points indicate the cue angle or ball angle. Carets highlight how bump-cue offsets are relatively constant throughout closed loop sessions (gray) and vary dramatically in the dark session (purple). *Right:* Histogram of bump-cue offset across the entire session. The average offset vector length quantifies offset consistency. The orange line denotes an offset of 0 radians. h) Average bump-cue offset vectors from [g] in polar coordinates. i) Offsets are more consistent in closed loop vs dark sessions. Each dot is the length of the average bump-cue offset vector for a single fly in darkness or closed loop sessions. Lines connect data from the same fly. Wilcoxon signed-rank W=0, p=1.22×10^-4^ j) Mean bump-cue offsets across the two closed loop sessions. Lines connect data points from the same fly. k) Absolute difference in bump-cue offset between closed loop 1 and 2 for each fly. Thick line indicates median across flies. Bump-cue offsets are maintained across the dark period (Wilcoxon signed-rank test for difference from *π*/2 W=2, p=7.32×10^-4^) l) EPG activity bumps are narrower in closed loop than dark trials. *Top:* PVA angle centered EPG bump profile for closed loop (black) and dark (purple) trials. Shaded region shows across fly mean +/− sem. Bars show bump width (full-width half maximum) for across fly averages. *Bottom:* Difference in bump width (closed loop – dark) for each fly. Thick line indicates across fly median. Wilcoxon signed-rank W=1, p=2.44×10^-4^.

Anchoring to external sensory cues such as distal visual landmarks is crucial for head direction cells to maintain an accurate sense of direction^6,7^. Such cues can strongly influence head direction representations and correct for errors that accumulate from integrating self-movement signals over time. Indeed, when environments lack external cues, such as in darkness, the head direction representation drifts relative to the animal’s orientation^3,8–13^.

What are the mechanisms by which external sensory cues calibrate head direction cells? Classic models propose associative synaptic plasticity between sensory inputs and head direction neurons^14,15^, and in *Drosophila* there is considerable evidence that an associative process tethers the head direction representation to visual cues^16,17^ (Fig. 1c, Extended Data Fig. 1b). Yet, the core synaptic mechanisms underlying plasticity between sensory neurons and head direction cells is not known in any organism. We took advantage of the compact and highly organized head direction circuit in *Drosophila* to uncover the key components of plasticity between visual inputs and head direction neurons.

To identify the mechanism of visual synaptic plasticity we focused on the canonical head direction cells, EPG neurons, and their visual inputs. EPGs arborize in wedges around a donut-shaped midline structure called the ellipsoid body where adjacent wedges encode adjacent angles of head direction (Fig. 1a). As the fly turns, a single “bump” of increased neural activity swings around the donut like the needle of a compass (Fig. 1c). External sensory cues that serve as landmarks enter this network via a large population of inhibitory neurons called ER neurons (also known as ring neurons, Fig. 1b) with different ER neuron subtypes encoding distinct external cues such as visual features (ER4d, ER2a-d, and likely some ER3 subtypes), linear polarization (ER4m and likely ER5 and ER3w), and wind direction (ER1 and ER3a)^18–23^. Each sensory ER neuron releases γ-Aminobutyric acid (GABA) from a ring-shaped axon that synapses onto every EPG cell creating all-to-all connectivity between sensory inputs and head direction cells^24–31^. In this work, we focus on visual ER neurons, which have local receptive fields that tile the fly’s field of view^21,23^.

Associative plasticity of ER→EPG synapses is proposed to flexibly link the head direction representation to visual cues (Fig. 1b,c). Prior work suggests that synapses between co-active ER neurons and EPG cells depress (Fig. 1b,c, Extended Data Fig. 1a,b)^16,17^. As a result of this plasticity, recurring experience of a visual cue at a particular position on the retina reliably disinhibits the same subset of EPG neurons. This pushes the EPG bump to the same location in the ellipsoid body for each corresponding azimuthal location of the cue, resulting in head direction activity that is robustly anchored to the fly’s current visual surroundings (Fig. 1c).

Explicitly, when a fly encounters a novel environment, visual synaptic weights are predicted to be uninformative and the head direction network’s estimate relies solely on error-prone integration of self-movement signals. Through exploration, visual synapses become sculpted, allowing the network to error-correct using visual cues^32,33^. Indeed, the relationship between visual cues and the EPG head direction representation is not hard-wired, suggesting it forms *de novo* in each environment ^3,16,34^ (Fig. 1d, Extended Data Fig. 1b). Optogenetic pairing or disorienting visual experience can also induce changes in the visual-to-EPG mapping, providing evidence that visual tethering is associative^16,17,35^. Thus, plasticity at the GABAergic synapses between ER and EPG neurons serves as a form of spatial memory that keeps the brain’s internal compass aligned with current surroundings. However, the mechanisms that mediate associative plasticity at this inhibitory synapse are unknown^6,36^. We discovered that a population of octopaminergic neurons provides feedback from head direction cells (EPG neurons) to their visual presynaptic partners (ER neurons). This octopamine pathway is both necessary and sufficient for plasticity. We propose a model where octopamine instructs plasticity by signaling to presynaptic sensory axons when postsynaptic head direction cells are active, implementing a retrograde signal via a circuit mechanism.

### The head direction representation stores its relationship to visual cues

We first established that head direction cells anchor to a visual cue and store this relationship over many minutes in our experimental set-up. We performed two photon (2P) calcium imaging in the ellipsoid body from EPG neurons while a fly walked on an air-cushioned ball in darkness or in virtual reality (VR) with a visual cue (Fig. 1e). In the initial trial, a visual cue tracked the fly’s angular orientation as the fly walked on the ball, simulating a cue at infinity (closed loop 1). We decoded the EPG head direction angle at each timepoint by calculating the population vector average (PVA) using fluorescence values from each ellipsoid body wedge (Fig. 1f, see Methods^3^). With a visual cue to orient the fly, the EPG activity bump (PVA angle), closely tracked the cue; the angular relationship between the EPG bump and VR cue was consistent over time (see Methods, Fig. 1g-i, Extended Data Fig. 1c-g, Supplementary Video 1). To assess the impact of this visual cue on compass tracking, next we had the fly explore in darkness. To quantify tracking consistency in darkness, we compared the offset between the EPG bump and the rotation of the ball (which is the signal used to animate the cue location in closed-loop). In darkness, the offset between the bump angle and the fly’s orientation in darkness (bump-ball offset) was highly variable (Fig. 1i)^3^. Next, we assessed whether the network’s relationship to the visual cue is stored. After the fly explored in darkness for 6 minutes, we returned it to the original VR environment with the same visual cue (closed loop 2). We found that the network’s relationship to the visual cue was stored across this experience (Fig. 1g,h). In the majority of flies, a consistent bump-cue offset was maintained between the first and second exposure to the cue (Fig. 1j,k). A small subset of flies remapped, exhibiting a distinct bump-cue offset on the second closed loop trial. Prior studies observed that the head direction network can recall its relationship with a single visual cue or a complex scene over similar timescales^16,34,37^. Thus, our data, together with previous work, suggest that the learning mechanism that anchors head direction cells to visual cues can reliably store a visual association for minutes. As further evidence that the EPG population activity is shaped by sculpted synaptic weights, we found that the EPG bump is narrower during VR sessions with a visual cue than during dark sessions (Fig. 1l) consistent with a previous report^34^. Moreover, the robust recall of the EPG bump-cue offsets following exploration in darkness favors models that require visual experience for synapses to either strengthen or weaken^16^. Corroborating these results, our data was better fit by computational models in which both synaptic potentiation and synaptic depression require presynaptic activity (Extended Data Fig. 2). Such “presynaptically-gated” models were previously shown to increase the capacity of head direction networks to map different visual scenes^16^.

### Presynaptic neuromodulators track locomotion and head direction

To uncover how synaptic plasticity is implemented, we examined candidate pathways that could mediate the detection of coincident activity between presynaptic ER neurons and postsynaptic EPG neurons. Previous results support a plasticity rule where coincident ER and EPG activity triggers synaptic depression of the GABAergic synapse that connects them^16,17,38^. In mammalian circuits, associative depression at GABAergic synapses often occurs in the presynaptic terminal and requires a retrograde signal from the postsynaptic cell^39^. This retrograde signal is then bound by receptors on the presynapse and coincidence between this signal and presynaptic activity induces depression^40–42^ but see^43^.

The logic of a retrograde signaling mechanism is appealing given the circuit’s anatomy. If coincidence detection occurs on the presynaptic side (in the ER axons), plasticity cues do not need to be perfectly synapse-specific as cross-talk between adjacent synapses would not be detrimental^44–48^. If second messengers from a retrograde signal diffused through a local segment of ER axon (Fig. 2a), they would preferentially affect ER-EPG presynapses in the same wedge. All EPG neurons within the same wedge code for similar head directions, so all ER-EPG synapses within a wedge should ideally change in the same direction to support learning. By analyzing connectomics data we found that the minimum intracellular cable distance between ER4d presynapses onto different head direction-tuned EPGs, is often many micrometers (Fig. 2a,b). A signal that spreads locally in the ER axon would therefore rarely reach presynapses onto EPGs with highly distinct head direction tuning.

**Figure 2:**
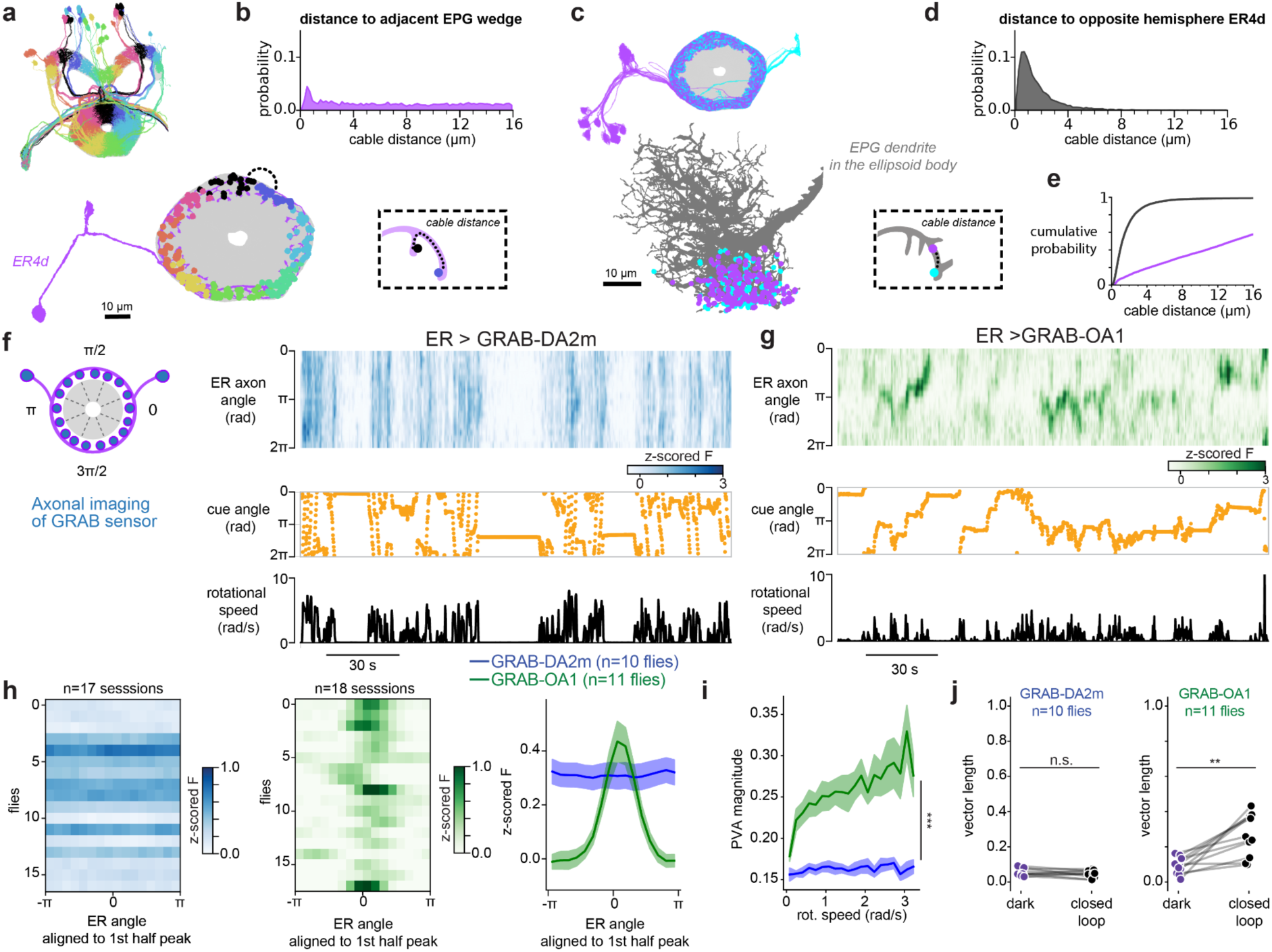
Evidence for presynaptic coincidence detection and neuromodulation. a) ER4d-EPG presynapses are organized by EPG HD tuning (rainbow). *Top:* EPG neurons, colored by their ellipsoid body wedge. *Bottom:* Single ER4d neuron with presynapses highlighted as in *(top)*. Inset highlights the “cable distance” (dotted line) between presynapses in neighboring wedges. b) Histogram of cable distances between ER4d-EPG presynapses in adjacent wedges. For each presynapse, we calculate the distance through the ER4d axon to the closest presynapse in a different wedge. c) EPG arbors have intermixed postsynapses from ER4d neurons with different visual tuning. *Top:* All ER4d neurons from the left (magenta) and right (cyan) hemisphere. *Bottom:* ER4d synapses onto a single EPG dendrite. Inset highlights cable distance (dotted line) between synapses from different ER4d sources. d) Histogram of shortest cable distance between pairs of ER4d-EPG postsynapses with ER4d sources from different hemispheres. e) Cumulative probability of presynaptic (b, magenta) and postsynaptic (d, black) cable distances. f) *Left:* Schematic of imaging from ER axons expressing a GRAB indicator *Right:* Example GRAB-DA2m imaging data from ER axon terminals. Cue angle and fly rotational speed are shown below. g) Same as [f] for GRAB-OA1 imaging from ER terminals h) GRAB-OA1 but not GRAB-DA2m shows a bump of activity that encodes head direction. Each row in the heatmaps shows the temporal average of data from one session, where each frame was shifted to align with a cross-validated fluorescence peak (*left:* GRAB-DA2m, *middle:* GRAB-OA1, *right:* across fly average (sessions averaged within fly first then across), blue-GRAB-DA2m, green-GRAB-OA1). Shaded regions show +/− sem. i) PVA magnitude quantifies the bump amplitude of GRAB-OA1 (green) and GRAB-DA2m (blue) plotted across rotational speeds (closed loop data only). Mixed effects ANOVA on log-transformed values, GRAB sensor main effect: F=32.7, p=1.12×10^-5^. j) Lengths of average bump-cue offset vectors are larger in closed loop sessions for GRAB-OA but not GRAB-DA2m. Same as (Fig 1i) for GRAB-DA2m (left) and GRAB-OA1 (right). Mixed effects ANOVA on logit-transformed values, GRAB sensor main effect: F=37.1 p=7.44×10^-6^, closed loop vs dark main effect: F=8.98 p=0.007. Interaction: F=18.7 p=3.61×10^-4^. Posthoc tests for difference between dark and closed loop (holm-corrected p values): GRAB-DA2m t=1.40 p=0.195, GRAB-OA1 t=4.39 p=0.003. Posthoc test for difference between GRAB-DA2m and GRAB-OA1 closed loop: t=7.03, p=2.21×10^-6^.

By contrast, if plasticity induction occurs postsynaptically, in the EPG dendrite, interspersed ER-EPG postsynapses from different ER visual sources need distinct weight changes. According to the model that best reproduces our data (Extended Data Fig. 2), only synapses from active ER neurons should depress ^16^. Connectome analysis shows that ER4d-EPG postsynapse pairs from opposite hemispheres, and thus distinct visual tuning, are typically separated by less than 2 micrometers (Fig 2c-e). An intracellular signaling molecule that indicates coincident activity between an ER and EPG neuron may easily diffuse to nearby silent ER postsynaptic sites on the same EPG, causing incorrect ER synapses to undergo plasticity (Fig. 2c).

While anatomy favors presynaptic coincidence detection, no previously described retrograde signaling mechanism is a strong candidate for this circuit. Flies do not have direct homologs for common retrograde pathways in mammals such as endocannabinoid signaling or BDNF/Trk-B^49–51^ and, enzymes for gaseous or peptidergic retrograde transmitters^52–57^ are not highly expressed in EPG neurons^27^ (Extended Data Fig. 3a). Inspired by studies of plasticity in other circuits^58–60^, we wondered whether a monoamine released by a third neuron onto presynaptic terminals could function like a retrograde signal. A monoaminergic signal that conveys postsynaptic activity to the presynaptic terminal would exhibit a highly local release pattern that closely tracks the EPG head direction activity. We focused on two monoamines: dopamine and octopamine. We chose dopamine because previous work suggests that dopamine regulates head direction plasticity^38^. We also explored octopamine, a common invertebrate neuromodulator that is similar to norepinephrine, because of recent reports of a putative octopaminergic cell type with arbors that tile the ellipsoid body in a similar pattern to EPG cells^26,31,61^. Furthermore, RNA sequencing and receptor knock-in studies suggest that many classes of ER neurons express dopamine and octopamine receptors^27,62–65^. We confirmed using expansion-assisted fluorescent *in situ* hybridization (EASI-FISH)^66^ that putatively visual ER neurons (ER4d and ER3p) and mechanosensory ER neurons (ER1) contain transcripts for multiple octopamine receptors (Extended Data Fig. 3b-e).

To assess whether dopamine or octopamine signals head direction information onto ER presynaptic terminals, we measured the release patterns of these monoamines using the fluorescent dopamine sensor, GRAB-DA2m^67^, and, separately, the fluorescent octopamine sensor, GRAB-OA1^68^. We expressed either sensor in ER neurons using a split-Gal4 driver line we developed that strongly and specifically labels a large population of these cells (Extended Data Fig. 4). We then performed 2P imaging of ER neuron axons in the ellipsoid body while the fly walked in VR.

We observed that dopamine (GRAB-DA2m) signaling was distributed globally across the ER axons. Bursts of dopamine release were relatively uniform across the entire ellipsoid body and peaked when the fly made rapid turns (Fig. 2f). Each of the four dopamine neurons in this region, called ExR2 cells, have arbors that span the full ellipsoid body. These imaging data therefore suggest that dopamine release from their axons is not compartmentalized. These data are also consistent with calcium imaging showing that ExR2 activity is locked to the fly’s rotational speed^38,69^.

In contrast, octopamine signaling was highly localized and tracked the fly’s head direction in VR (Fig. 2g). To visualize head direction tracking, we first aligned all timepoints by the heading of the fly. We then averaged the first half of the data over time and found the phase of the largest peak in fluorescence (see Methods, Extended Data Fig. 5a,b). That phase was then used to shift held-out imaging data to align the peaks across flies (Fig. 2h, Extended Data Fig. 5). For octopamine, this cross-validated analysis revealed a peak that was consistent across flies (Fig. 2h), indicating that octopamine release tracks the flies’ head direction. When we performed the same analysis on dopamine, we found no evidence that dopamine tracks heading as the heading-aligned data showed a uniform activity profile (Fig. 2h). Quantitatively, the GRAB-OA1 bump magnitude (PVA magnitude, see Methods) was significantly larger than the GRAB-DA2m bump magnitude (Fig. 2i, Extended Data Fig. 5c). The GRAB-OA1 bump magnitude also increased further when the fly walked with high rotational speed, similar to observations of EPG population activity (Fig. 2i, Extended Data Fig. 1e & 5c)^3,12,13,38,70^. In addition, the bump-cue offsets were more consistent in closed loop trials than darkness trials for GRAB-OA1 but not for GRAB-DA2m (Fig. 2j). In summary, we find that dopamine signaling in the ellipsoid body is global, suggesting it does not signal head direction information to ER presynaptic terminals. Global motor-locked dopamine is well positioned, however, to modulate plasticity at times when the fly makes rapid turns (Extended Data Fig. 5d)^35,38,69^. On the other hand, octopamine signaling tracks head direction and is a strong candidate to convey EPG activity onto ER terminals for coincidence detection.

### EL neurons are octopaminergic and carry head direction activity

To identify the octopaminergic neurons that relay head direction information onto local sections of visual ER neuron axons we analyzed RNA expression levels and connectomic data (Fig. 3a). Recent work identified EL neurons, a columnar cell type that tiles wedges of the ellipsoid body and expresses the enzymes required for octopamine biogenesis^26,31^. We used two split-Gal4 driver lines that label EL neurons to confirm the neurotransmitter identity of these cells. Using RNA sequencing and FISH we confirmed that EL neurons are enriched for RNAs that encode enzymes necessary for octopamine production and vesicle loading (tyrosine decarboxylase-2: Tdc2, tyramine beta-hydroxylase: T*β*H, & vesicular monoamine transporter: VMAT) and found no transcriptomic evidence that EL co-produces any other fast neurotransmitter (Fig. 3b, Extended Data Fig. 6a, b).

**Figure 3.**
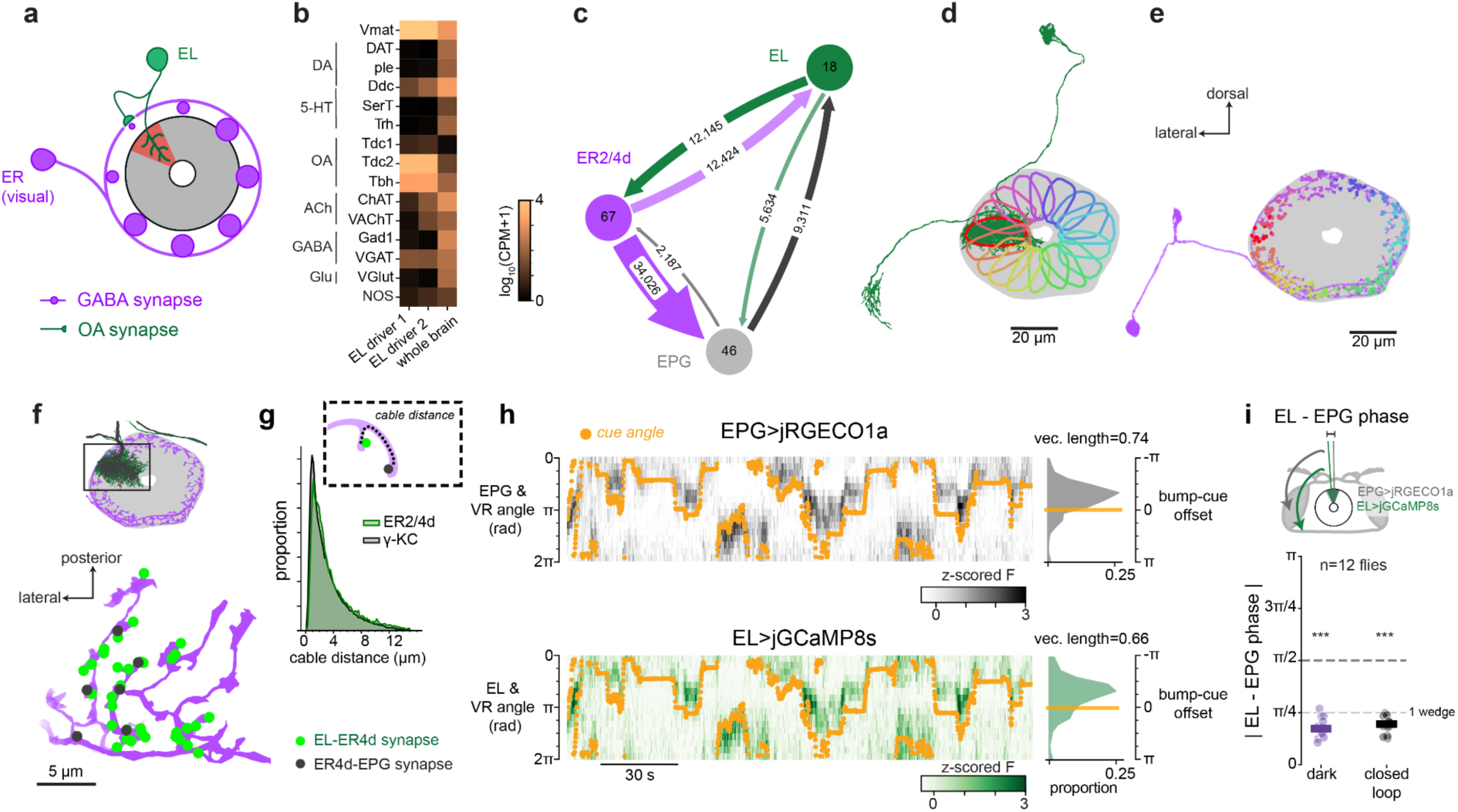
EL neurons are octopaminergic and carry head direction activity. a) Each EL neuron (green) innervates a slice of the ellipsoid body, forming a feedback loop that relays activity from the EPG bump (red) to segments of visual ER axons (purple). b) Bulk RNA sequencing shows EL neurons express mRNA for enzymes necessary for octopamine (OA) synthesis and vesicular packaging, but no other neurotransmitters. Heatmap shows average log counts per million (CPM) across replicates for two different EL split-Gal4 lines and a whole brain reference. c) Synapse counts between visual ER neurons (ER2 and ER4d), EPG neurons, and EL neurons from Hemibrain connectome^30,132^. Numbers in circles indicate the number of neurons in each class. Arrow width indicates synapse counts (also shown numerically). d) EL neurons tile the ellipsoid body (grey). EM reconstruction of a single EL neuron (green). The most dense region of presynapses from the highlighted EL neuron are outlined in red (peeled convex hull, see Methods) while other colors indicate presynapse dense regions for the 17 other EL neurons. e) ER4d neurons receive inputs from every EL neuron around the ellipsoid body. EM reconstruction of a single ER4d neuron (purple) with input synapses from EL neurons are colored according to EL neuron identity as in [d]. f) EL-ER synapses are intermingled with ER-EPG synapses. *Top*: EM reconstructions of example interconnected ER4d (purple), EPG (black) and EL (green) neurons. *Bottom:* Close up of the ER4d axon segmented with EL synaptic inputs shown in green and EPG synaptic outputs in black. g) Histogram of the cable distances through the ER neuron (ER2 & ER4d) arbors from each ER-EPG synapse to the closest modulatory (EL-ER) contact (green). For analysis of all ER subtypes see Extended Data Fig. 8c. For comparison, cable distances through *γ*-kenyon cell (KC) arbors between each *γ*-KC-mushroom body output neuron (MBON) synapse and the closest modulatory (dopamine neuron [DAN]-*γ*KC) synapse are shown in black. h) Example closed loop VR dual imaging session EPG (top) and EL (bottom). Plotted as in Fig. 1g. i) EL and EPG activity bumps have similar phases. Average absolute differences in bump angle (PVA angle) are plotted for each fly for dark and closed loop sessions. Differences are significantly less than chance (*π*/2) in both conditions by Wilcoxon signed-rank test. holm-corrected p-values: dark W=0 p=9.77×10^-4^, closed loop W=0 p=9.77×10^-4^. One wedge is defined as the average angular span of a single EPG dendritic arbor.

We also made several notable observations from RNA sequencing that we do not investigate further in this work. Consistent with a previous report, EL neurons appear to express the peptide SIFamide (Extended Data Fig. 6b)^26,71^. EL neurons also express receptors for dopamine and serotonin, suggesting that octopamine signaling may be regulated by other monoamines. ExR2 neurons, on the other hand, do not express high levels of octopamine or serotonin receptors (Extended Data Fig. 6b). Lastly, EL neurons express transcripts encoding voltage-gated sodium channel subunits at high levels, and whole-cell electrophysiological recordings from EL neurons showed that they fire action potentials (Extended Data Fig. 6b, c), similar to other cell types in the central complex ^13,17,72,73^.

We next used connectomic data^30^ to show that EL neurons are uniquely positioned to instruct plasticity at ER-EPG synapses. EL neurons receive strong synaptic input from EPGs within the ellipsoid body and, in turn, send the majority of their synaptic outputs to nearby sensory ER neuron axons (Fig. 3c, Extended Data Fig. 7)^31^. ELs also receive input from PENa–a class of columnar neurons (i.e., a cell type that tile the ellipsoid body wedges) that tracks head direction conjunctively with angular velocity and shifts the EPG bump around the ellipsoid body in response to self-motion^12,13^ via recurrent connections with EPGs. Our analysis shows that EL neurons are the best candidate to implement a local feedback loop from head direction neurons onto sensory axons (Fig. 3d). EL neurons make substantially more synapses onto sensory ER neurons than any other columnar cell type (Extended Data Fig. 7c-k). For ER terminals to flexibly learn to anchor to the EPG activity bump, ER axons need to receive local feedback about EPG activity in each wedge. Accordingly, visual ER neurons (ER4d, ER2, ER3p subtypes) receive synaptic input from EL neurons at all angles around the ellipsoid body (example ER4d-Fig. 3e, Extended Data Fig. 7c-k). Furthermore, we found that octopaminergic inputs onto visual ER neurons were positioned in close enough proximity to modulate the strength of ER-EPG synapses (Fig. 3f,g, Extended Data Fig. 8a-c). Examining a stretch of ER4d axon (purple), we see that the EL-mediated feedback loop is extremely tight. The outputs from an ER4d neuron onto a single EPG (black) are highly interspersed with inputs from an EL neuron that is downstream of the same EPG (green, Fig. 3f). To approximate the distance from bound octopamine receptors to the synapses they would modulate, we calculated the intracellular path through the ER axon (i.e. cable distance) from each GABAergic ER->EPG output (presynapse) to the closest octopamine input (EL->ER postsynapse). From this, we estimate that octopamine receptor second messengers would need to diffuse intercellularly less than 2 *μ*m to modulate the majority of ER4d-EPG synapses (Fig 3g). Extrasynaptic binding of octopamine through bulk transmission or spillover would shorten this distance even further. We think this distance is plausible for modulating synaptic strength, because it closely matches the cable distances for well established dopamine-driven plasticity onto gamma kenyon cells in the mushroom body (Fig. 3g)^74–76^. We also found similar cable distances when performing these calculations for other ER subtypes, indicating EL feedback is a plausible mechanism for plasticity across subtypes (Extended Data Fig. 8c).

Multiple factors argue that EL affects plasticity by acting presynaptically. First, presynaptic ER neurons strongly express multiple octopamine receptors spanning G*α_s_*-, G*α_q_*– and G*α_i_*_/*o*_-coupled receptors meaning octopamine can stimulate multiple signaling cascades implicated in plasticity (Extended Data Fig. 3b-e)^62,65,77^. EPG neurons have only modest transcript levels for two G*α_s_*-coupled receptors (Extended Data Fig. 3b). Second, as mentioned above, a diffusible plasticity cue derived from EL octopamine signaling could feasibly instruct wedge-specific plasticity on the presynaptic ER side, but not on the postsynaptic EPG side (Fig. 2a-e). Third, EL neurons make more synapses onto ER than EPG neurons (Fig. 3c, Extended Data Fig. 7c). Fourth, using an *ex vivo* preparation we found that ER axons show much larger increases in cAMP (GFlamp2) in response to EL stimulation than EPG neurons (Extended Data Fig. 8d,e). Together, our GRAB data, *ex vivo* cAMP imaging, sequencing, and connectomic analyses identify EL neurons as a strong candidate for relaying local EPG activity onto ER neuron terminals. This octopaminergic signal is well positioned to enable coincidence detection that could instruct presynaptic plasticity of ER-EPG synapses.

To show that EL population activity tracks EPG activity, we performed simultaneous imaging from EL neurons and EPG neurons by using orthogonal genetic driver lines for the two populations (EPG-Gal4 & EL-LexA) to express two spectrally separable calcium indicators (jGCaMP8s and jRGECO1a) (Fig. 3h, see Extended Data Fig. 9 for EL-LexA characterization)^78–80^. We found that both cell types exhibited a clear bump of neural activity that tracked head direction when the fly walked in VR or in darkness (Fig. 3h,i). Importantly, when the activity of both cell types was measured simultaneously, they circled the network together. When the fly walked in VR, the two neural populations had extremely similar bump-cue offsets (Fig. 3h) and the phases of the EL bump and EPG bump were nearly identical when the fly walked in VR or darkness (mean absolute difference <*π*/4 Fig. 3i). Significant deviations in EL and EPG bump phases only occur when the amplitude of the bumps (PVA magnitude) is small, indicating that these deviations are likely driven by measurement noise (Extended Data Fig. 10a). Examining the dynamics of these populations more closely we found the EL bump slightly leads the EPG bump (Extended Data Fig. 10b,c). This effect is unlikely to be due to differences in indicator dynamics because we observed a similar phase difference in a separate cohort in which we controlled for this confound explicitly (Extended Data Fig. 10d-h). It is somewhat surprising that the EL population leads the EPG population since EL is largely postsynaptic to EPG, but we note that EL also receives input from PENa neurons (Extended Data Fig. 7a,e) that conjunctively encode head direction and rotation velocity and whose population activity leads the EPG bump in the ellipsoid body^12,13^. To confirm that EL neurons inherit head direction activity from EPG neurons, we acutely silenced EPG neurons using temperature sensitive shibire (shi^TS^)^81^ while imaging EL population activity. When EPG neurons were silenced, the bump of activity in EL neurons was abolished (Extended Data 10i-k). These data and a previous report, suggest that EPG output is central to generating an activity bump^27^. Collectively, these data argue that EL neurons inherit an activity bump from other columnar neurons (predominantly EPG and PENa) and then release octopamine locally near where head direction activity is highest.

### Octopamine synthesis and receptors are required for head direction activity to track visual cues

We hypothesized that this octopaminergic feedback loop is necessary for visual plasticity. To test this, we disrupted the ability of EL neurons to produce octopamine and measured changes in how the head direction network anchored to a visual cue. First, we used RNA interference (RNAi) to knock-down the octopamine synthesis enzyme T*β*H in EL neurons and confirmed T*β*H knockdown largely eliminated octopamine immunolabeling in EL somata (Extended Data Fig. 11a-c), a result that provides additional evidence these cells are octopaminergic. Then, we measured EPG neural activity as these flies walked in VR (Fig. 4a-c, Extended Data Fig. 11d,e). To quantify EPG visual tracking, we compared trials where the fly walked in darkness to trials with a visual cue. As also shown above (Fig. 1g-i), when control genotypes walked in darkness, their EPG bump drifted relative to the fly’s heading in darkness (low vector length), but when the cue was visible the EPG bump accurately tracked the cue angle (high vector length)(Fig. 4b,c). However, when octopamine synthesis was disrupted, EPG tracking of head direction was poor regardless of whether the fly walked in darkness or was given a visual cue (Fig. 4c). Moreover, in flies with reduced EL octopamine, the EPG activity was significantly less tethered to the cue than in either control genotype (Fig. 4c). We also obtained similar results in a parallel set of experiments where we instead used EL population activity to measure bump position. This allowed us to use a more specific EL split-Gal4 driver line for T*β*H knockdown (Extended Data Fig. 11f-i). Importantly, network deficits caused by disrupting T*β*H were specific to the EPG bump’s ability to track a visual cue, as the movement of the EPG (or EL) bump still tracked the fly’s angular rotation, as in control genotypes (Extended Data Fig. 11d,e,h,i). We note that in this data set we were unable to detect changes in EPG bump width between darkness and VR trials in either control or experimental groups (data not shown). It is likely this is because we used a slower calcium indicator for these experiments (jGCaMP7b)^70^. In summary, when EL neurons lack octopamine, the dynamics of the head direction representation are similar to conditions without visual cues. Our interpretation is that this perturbation disrupts synaptic plasticity between visual inputs and head direction cells.

**Figure 4.**
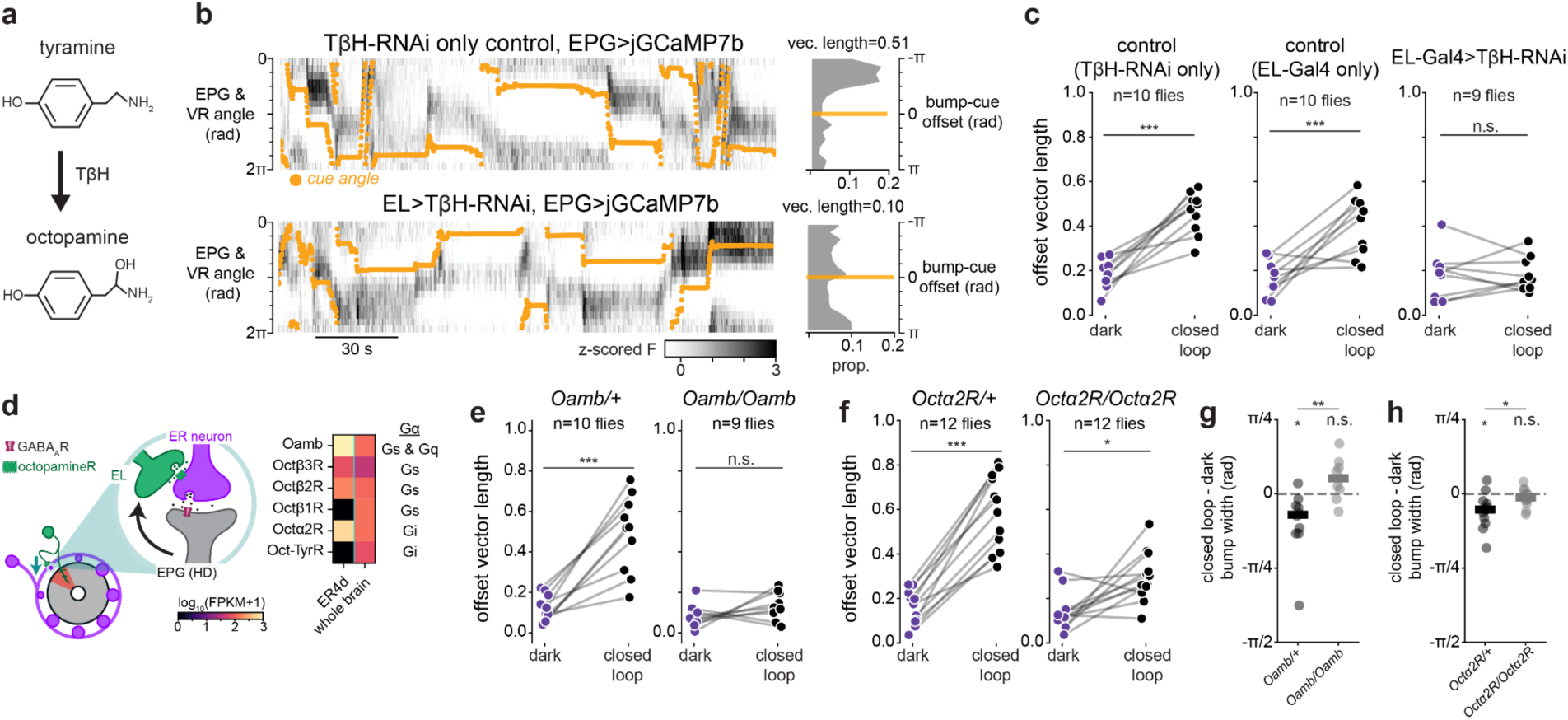
Octopamine synthesis and receptors are required for visual tethering. a) Schematic showing final step of octopamine synthesis pathway. b) Example EPG imaging data in closed loop from a control fly (*top:* T*β*H-RNAi only) and experimental fly with T*β*H-RNAi expressed in EL neurons (*bottom:* EL-Gal4>Tβ H-RNAi). Data plotted as in Fig. 1g. c) EL T*β*H-RNAi prevents the EPG bump from learning a visual scene. Lengths of average bump-cue offset vectors are plotted as in Fig. 1i. Vector lengths are not different between darkness and closed loop when flies lack octopamine in EL neurons. Mixed effects ANOVA on logit-transformed vector lengths: main effect of genotype F=8.36 p=1.57×10^-3^; main effect of dark vs closed loop F=42.2 p=6.97×10^-7^; interaction F=7.31 p=0.003. Posthoc comparisons (p-values holm corrected): darkness vs closed loop – T*β*H-RNAi only: t=6.47 p=3.46×10^-4^ EL-Gal4 only: t=4.20 p=0.002 EL-Gal4>T*β*H-RNAi: t=0.515 p=0.621. Closed loop comparisons (independent t-tests, p-values holm-corrected): T*β*H-RNAi only vs EL-Gal4 only: t=1.09 p=1.00, T*β*H-RNAi only vs EL-Gal4>T*β*H-RNAi: t=6.69 p=4.04×10^-5^, EL-Gal4 only vs. EL-Gal4>T*β*H-RNAi: t=4.70 p=0.001. d) ER4d neurons express multiple octopamine receptors that could mediate EL-driven plasticity. *Right:* ER4d averaged low cell RNA sequencing^27^. Log FPKM values with a whole brain reference. e) Oamb is required for EPG visual tethering. Lengths of trial average EPG bump-cue offset vectors for heterozygous (*Oamb*/+) and homozygous (*Oamb*/*Oamb*) receptor knockout animals in darkness and closed loop. Mixed effects ANOVA on logit-transformed vector lengths: main effect of genotype F=25.9 p=9.14×10^-5^; main effect of dark vs closed loop F=24.3 p=1.26×10^-4^; interaction F=4.31 p=0.028. Posthoc comparisons (p-values holm corrected): darkness vs closed loop – *Oamb*/+: t=-7.40 p=8.20×10^-5^, *Oamb*/*Oamb*: t=-1.33 p=0.222. Closed loop comparison (independent t-test): t=-5.34 p=1.07×10^-4^. f) Oct*α*2R is required for EPG visual tethering. Lengths of trial average EPG bump-cue offset vectors for heterozygous (*Octα2R*/+) and homozygous (*Octα2R*/*Octα2R*) receptor knockout flies. Mixed effects ANOVA on logit-transformed vector lengths: main effect of genotype F=13.1 p=1.52×10^-3^; main effect of dark vs closed loop F=88.8 p=3.50×10^-9^; interaction F=3.74 p=0.004. Posthoc comparisons (p-values holm corrected): darkness vs closed loop – *Octα2R*/+: t=-11.2 p=4.64×10^-7^, *Octα2R*/*Octα2R*: t=-3.76 p=0.003. Closed loop comparison (independent t-test): t=4.91 p=6.48×10^-5^. g) Oamb is required for visual sculpting of the EPG bump. Change in EPG bump width (closed loop – dark). Wilcoxon signed-rank tests for difference in bump width (holm-corrected p-values): *Oamb*/+: W=2 p=0.012, *Oamb*/*Oamb*: W=7 p=0.074. Mann-Whitney U test for difference in genotypes: U=8 p=0.003 h) Oct*α*2R is required for visual sculpting of the EPG bump. Change in EPG bump width (closed loop – dark). Wilcoxon signed-rank tests for difference in bump width (holm-corrected p-values): *Octα2R*/+: W=6 p=0.014, *Octα2R*/*Octα2R*: W=20 p=0.151. Mann-Whitney U test for difference in genotypes: U=33 p=0.026

We next tested the effect of octopamine receptor mutants on the ability of EPG activity to track a visual cue. ER4d and other ER neuron types express transcripts for multiple octopamine G-protein coupled receptors (GPCRs) (Fig. 4d, Extended Data Fig. 3b-e)^62,65,77^. We selected Oamb and Oct*α*2R mutants for analysis because they are the highest expressed in ER4d, their coupled G-proteins are not redundant with other octopamine receptors (Fig. 4d), they have negligible transcript levels in EPG neurons (Extended Data Fig. 3b), and they are relatively sparsely expressed in the rest of the fly nervous system ^62,65,71,77,82–84^. Flies homozygous mutant for Oamb or Oct*α*2R showed deficits in the bump’s ability to track a visual cue, relative to heterozygous controls (EPG>jGCaMP8s, Fig. 4e,f, Extended Data Fig. 12)^79^. Interestingly, Oamb mutants also displayed an attenuated EPG bump, shown by a decrease in PVA magnitude across all conditions and rotational speeds (Extended Data Fig. 12b). EPG activity bumps from heterozygous control animals (Oamb/+ & Oct*α*2R/+) were narrower when a visual cue was available, indicating sculpted synaptic weights, while there was no effect of the visual cue in either group of octopamine receptor knockout flies (Fig 4g,h compare Fig. 1l). Taken together with the T*β*H knockdown result, these experiments support a model where the release of octopamine from EL neurons onto visual ER presynapses instructs plasticity via multiple octopamine receptors.

### Octopaminergic EL neurons activity is necessary for visual plasticity

The data so far supports a model in which EL neurons release octopamine onto visual ER presynaptic terminals allowing visual information and the head direction representation to be associated. That is, if an ER neuron terminal is active and also receives octopamine, then that local axon segment has detected coincident ER-EPG activity, and nearby synapses could undergo depression (Fig. 1b). However, our constitutive T*β*H knockdown experiments (RNAi) and whole-animal receptor mutants cannot distinguish between the acute role of octopamine for plasticity induction and its requirement to establish or maintain synaptic function. To test whether EL activity is required during plasticity induction we combined acute EL silencing (HcKCR1)^85^ with a previously established optogenetic plasticity protocol^16^. In this experiment, EPG neurons express CsChrimson and focal 2P optogenetics is used to generate bump activity at experimentally-defined locations around the network (Fig. 5a-c, Supplemental Video 2, Methods). Using this approach, we could entrain a new arbitrary bump-cue offset in the EPG network (Fig. 5d). For each wedge of the ellipsoid body, we paired localized activation of EPG arbors with the presentation of a visual cue at a consistent bump-cue offset (Fig. 5a-c)^16^. The optogenetic stimulation is moved through eight stimulation locations around the ellipsoid body and the visual cue is rotated correspondingly to maintain a constant bump-cue offset (Fig. 5c). Before and after this pairing protocol, we monitored EPG activity to assess each fly’s baseline bump-cue offset, and to measure changes to the network’s visual tethering following pairing. Each fly underwent two rounds of plasticity induction (0 radians and *π* radians in pseudorandom order) so that changes in bump-cue offset due to plasticity could be easily distinguished from random fluctuations in bump-cue offset or preexisting differences at baseline. This plasticity protocol robustly induced changes to the bump-cue offset that were aligned to the phase offset reinforced during pairing (Fig. 5d,f, Extended Data Fig. 13).

**Figure 5.**
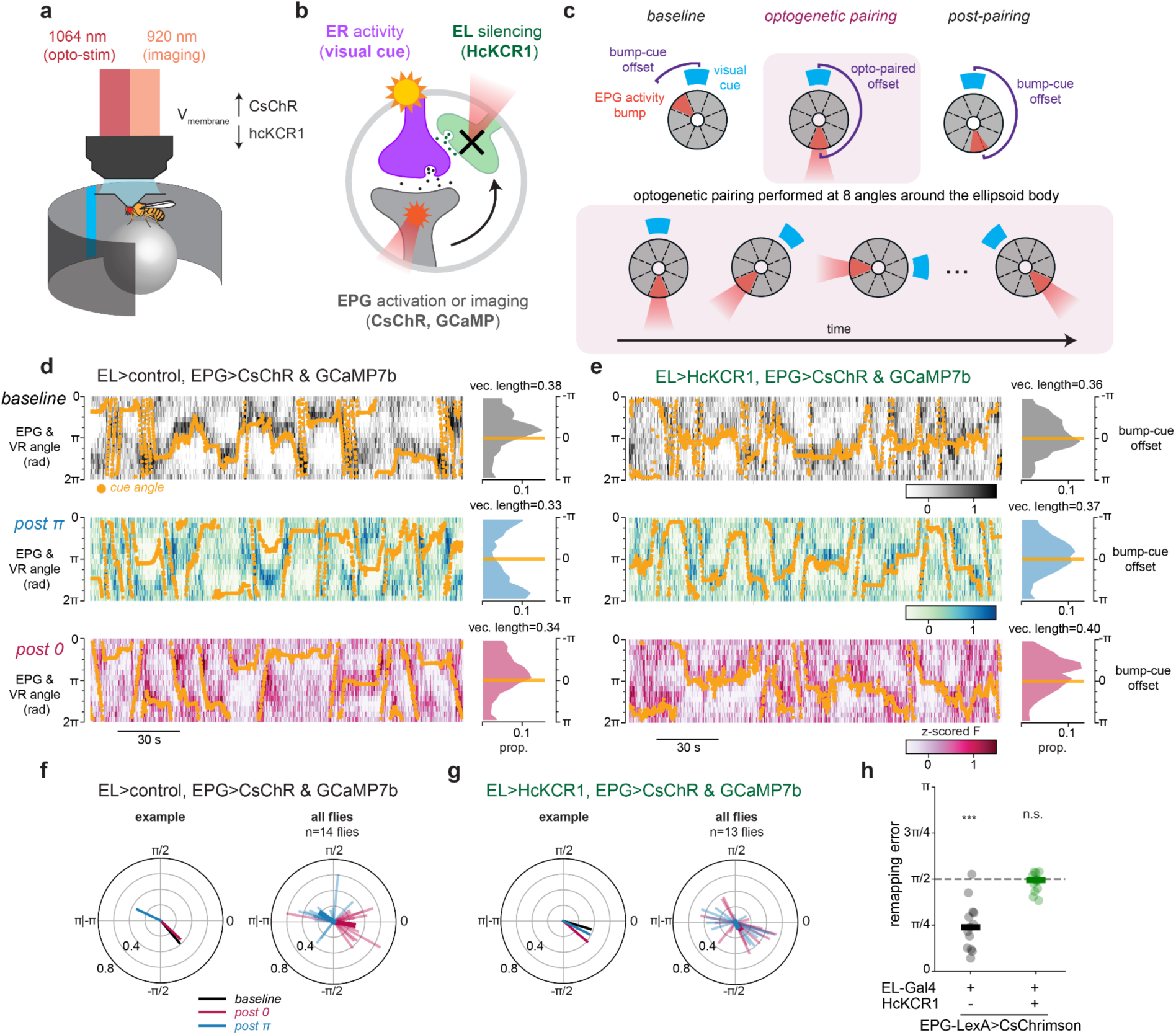
Octopaminergic EL neuron activity is necessary for plasticity induction. a) Schematic of simultaneous 2P imaging and 2P optogenetics. Flies walk in VR while two independent light paths are used for imaging (920 nm, resonant-galvo) and optogenetics (1064 nm, galvo-galvo). b) In the experiment a subset of ER neurons are activated by the visual cue. EPG activity is measured using GCaMP, and they are depolarized using CsChrimson. HcKCR1 is used to hyperpolarize EL neurons. c) In the optogenetic plasticity protocol, first flies walk in closed loop VR to measure the baseline EPG bump-cue offset (*top left*). Then a new bump-cue offset is entrained by pairing visual cue presentations with focal 2P optogenetics (*opto-pairing*, *π* offset shown in example). The bump-cue offset is measured again after 2P entrainment (post-pairing, *top right*). d) Example data showing successful EPG bump-cue plasticity induction in a fly with EPG stimulation but no EL inhibition. Imaging data as shown previously (*top-*baseline, *middle*-post *π* pairing, *bottom*-post 0 pairing). e) Example fly with EPG stimulation and EL inhibition showing no opto-pairing induced plasticity and instead a stable EPG bump-cue offset throughout (*top-*baseline, *middle*-post *π* pairing, *bottom*-post 0 pairing). f) Average bump-cue offset vectors for example shown in [d](*left*) and all control flies (*right)*. g) Same as [f] for flies with EL inhibition h) EL inhibition with HcKCR1 prevents optogenetically induced plasticity. Average remapping error for each fly. Lines show mean across flies. Wilcoxon signed-rank test for difference from *π*/2 (holm-corrected p-values): EL-Gal4 only: W=1 p=9.77×10^-4^, EL-Gal4>HcKCR1: W=29 p=0.273. Mann-Whitney U EL-Gal4 only vs EL-Gal4>HcKCR1: U=12 p=2.22×10^-4^.

Our hypothesis is that pairing EPG stimulation with a visual cue drives plasticity because focally activating EPG cells also recruits EL neurons innervating the same angle in the ellipsoid body. To test this directly we silenced EL neurons during plasticity induction using the potassium-selective inhibitory opsin (HcKCR1)^85^. First, we generated transgenic flies to express HcKCR1 with an epitope tag (V5) and used immunostaining to confirm that HcKCR1-V5 is well expressed by EL neurons (Extended Data Fig. 13a). In a separate validation experiment, we also confirmed using electrophysiological recordings that photostimulation of HcKCR1 robustly hyperpolarizes EL neurons (Extended Data Fig. 13b,c). HcKCR1 has a similar 2P absorption to CsChrimson, which allowed us to use the same laser (1064 nm) to both locally activate EPG neurons using CsChrimson and to locally silence EL neurons using HcKCR1 (Fig. 5a,b, Methods). Thus, in this experiment, optogenetic photostimulation simultaneously activates EPG neurons and hyperpolarizes EL neurons in the same location. We found that acutely silencing EL neurons in this manner completely disrupted optogenetically-induced plasticity (Fig. 5e,g,h). Notably, flies with HcKCR1 expressed in EL neurons exhibited an EPG bump that was well tethered to the cue at baseline, showing that visual plasticity is still intact in this genotype (Fig. 5e, Extended Data Fig. 13f). However, the optogenetic pairing protocol no longer induced new bump-cue offsets, and most flies retained the same bump-cue offset as the baseline session across the full plasticity protocol (Fig. 5e,g, Extended Data Fig. 13g). We quantified the effectiveness of optogenetic pairing by measuring the average angular error between the entrained bump-cue offset during optogenetic pairing and the offset exhibited afterwards in the post-pairing trial (‘remapping error’). Flies that expressed CsChrimson in EPG neurons without EL silencing had low remapping error, indicating robust plasticity induction, while flies that expressed HcKCR1 in EL neurons showed no evidence of plasticity and thus a high remapping error (Fig. 5h). EL silencing led to an average remapping error of around *π*/2 which is what we would expect if there was no plasticity and the bump-cue offset stayed unchanged across the two optogenetic pairing bouts (Extended Data Fig. 13g).

One explanation for this result is that EL silencing abolished our ability to optogenetically generate a bump in the EPG population. However, we could robustly move the EPG bump in this genotype (Extended Data Fig. 13d-e), which favors the interpretation that plasticity requires local EL activity at the position of the EPG bump. When EL neurons at that location are silenced during induction, plasticity does not occur and the network retains its previously learned relationship to the visual cue. In summary, EL octopamine release, and more specifically, the activity of EL neurons at the location of peak EPG activity is necessary for the head direction network to learn a new association with a visual cue.

### Localized EL activity and visual experience alone are sufficient for plasticity

We next sought to find the minimal circuit conditions that can elicit plasticity. We hypothesized that octopamine release onto ER terminals acts as the proximal signal for plasticity, which predicts that we should be able to bypass EPG activity by directly stimulating EL neurons to drive plasticity. To do this, we modified our optogenetic plasticity protocol by replacing EPG>CsChrimson activation with EL>CsChrimson activation. This new plasticity protocol was also successful in instructing new EPG bump-cue offsets by pairing EL stimulation with a visual cue (Fig. 6a-d, Extended Data Fig. 14, Supplemental Video 2). Importantly, optogenetically stimulating EL neurons without a visual cue did not systematically shift EPG bump-cue offsets (Fig. 6d, Extended Data Fig. 14c). Additionally, when we repeated the same pairing protocol in a control genotype with no CsChrimson expression, EPG bump-cue offsets did not change (Fig. 6d, Extended Data Fig. 14c) showing that laser induced heating of the tissue does not cause this effect. Therefore this EL-induced plasticity is associative exactly like EPG-driven plasticity^16^; changes in bump-cue offset required the simultaneous presentation of a visual cue and EL optogenetic stimulation. In separate experiments in which we imaged EL, instead of EPG, and also optogenetically stimulated EL, we confirmed that the EL activity bump moves to the location of optogenetic stimulation and that the EL bump-cue offset is also plastic, consistent with EL carrying a copy of columnar head direction neuron activity (Extended Data Fig. 15).

**Figure 6.**
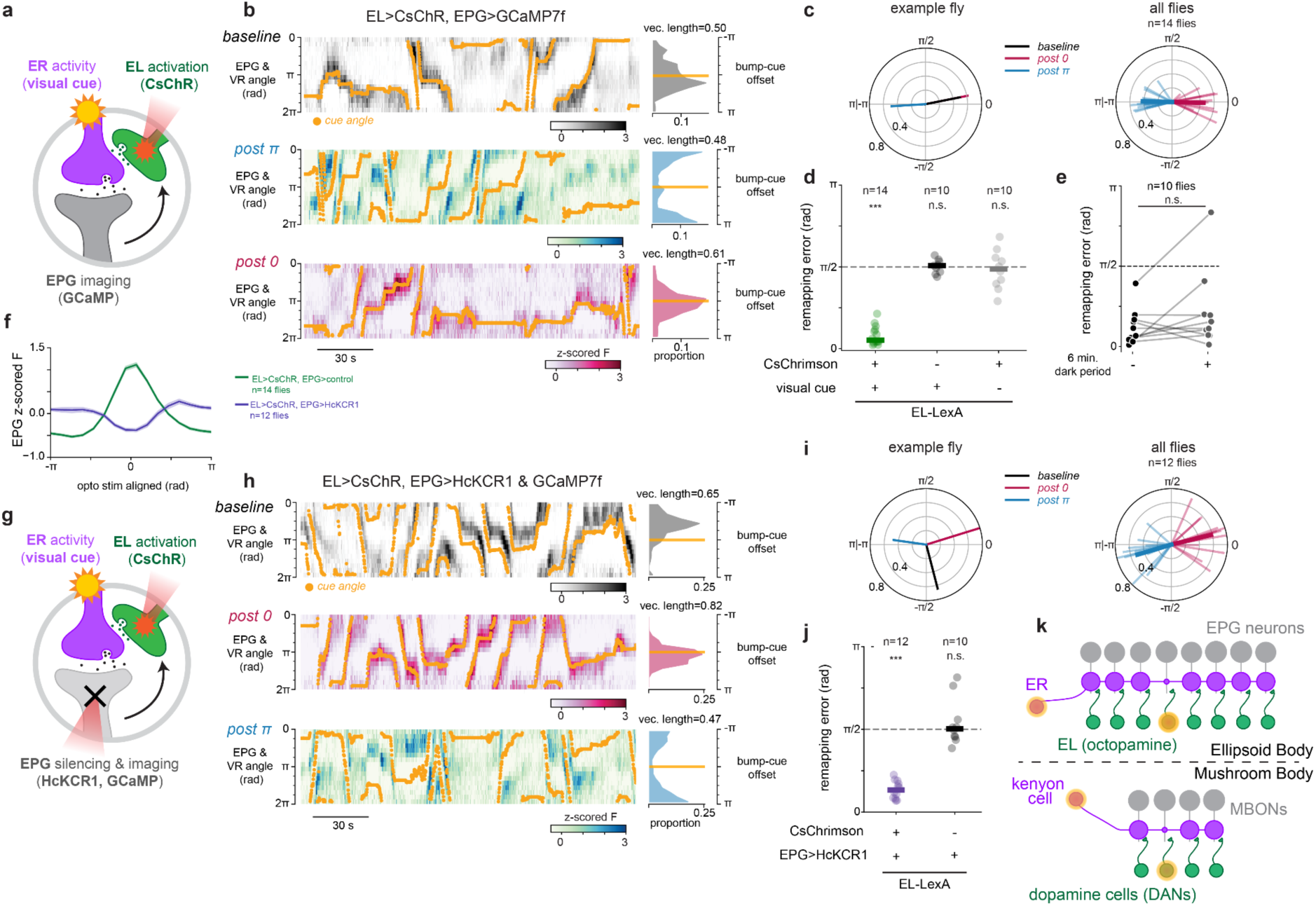
Localized EL activity and visual experience alone are sufficient for plasticity. a) Illustration of optogenetic plasticity strategy as in Fig. 5b, but now EL neurons are stimulated using CsChrimson (EL-LexA>CsChrimson) while EPG neurons are imaged (EPG-Gal4>jGCaMP7f). b) Example imaging data showing successful EL-induced EPG bump-cue plasticity. Data as shown in Fig. 1 & 5. c) Average bump-cue offset vectors for example data in [b] (*left*) and all flies from this genotype (*right*). d) EL-induced plasticity is associative. Data as shown in Fig. 5h. “n” shows the number of flies. Wilcoxon signed-rank test for difference from *π*/2 (holm-corrected p-values): EL-LexA>CsChrimson: W=0 p=3.66×10^-4^, EL-LexA only: W=20 p=1.00, EL-LexA>CsChrimson no visual cue: W=24 p=1.00. Kruskal-Wallis test for differences in experimental conditions: H=23.5 p=7.83×10^-6^. Posthoc Mann-Whitney U tests (holm-corrected p-values): EL-LexA>CsChrimson vs EL-LexA only: U=0 p=1.65×10^-4^, EL-LexA>CsChrimson vs EL-LexA>CsChrimson no visual cue: U=0 p=1.41×10^-4^, EL-LexA only vs EL-LexA>CsChrimson no visual cue: U=50 p=0.713. e) EL-induced plasticity is stable over prolonged dark periods. Each fly experiences standard optogenetic pairing (black) or optogenetic pairing followed by a 6 minute dark period (grey) before the bar reappears in closed loop. Data as shown in [d]. Wilcoxon signed-rank test for difference from *π*/2 (holm-corrected p-values): standard remapping protocol: W=0 p=0.006, pairing followed by 6 minute dark period: W=4 p=.027. Wilcoxon signed-rank test for difference between conditions: W=16 p=0.275. f) EL stimulation excites EPGs, and EPG inhibition is effective regardless of EL stimulation. EPG fluorescence in response to optogenetic stimulation, centered on the stimulated wedge (green: EL-LexA>CsChrimson EPG-Gal4>jGCaMP7f, purple: EL-LexA>CsChrimson EPG-Gal4>HcKCR1 & jGCaMP7f). g) Illustration of plasticity strategy as in [a]. EL neurons are depolarized using CsChrimson while EPG neurons are hyperpolarized using HcKCR1 (EL-LexA>CsChrimson, EPG-Gal4>HcKCR1 & jGCaMP7f) h) Example data showing successful EL-induced EPG bump-cue plasticity despite EPG inhibition. i) Average bump-cue offset vectors for example data in [h] (*left*) and all flies from this genotype (*right*). j) EPG depolarization is not required for EL-induced bump-cue offset plasticity and EPG hyperpolarization alone does not induce bump-cue offset plasticity. Remapping errors as shown previously. Wilcoxon signed-rank test for difference from *π*/2 (holm-corrected p-values): EL-LexA>CsChrimson EPG-Gal4>HcKCR1: W=0 p=9.77×10^-4^, EL-LexA only EPG-Gal4>HcKCR1: W=25 p=0.846. Mann-Whitney U test EL-LexA>CsChrimson EPG-Gal4>HcKCR1 vs EL-LexA only EPG-Gal4>HcKCR1: U=0 p=8.73×10^-5^ k) Schematic of similar plasticity rules in the ellipsoid body (*top*) and the mushroom body (*bottom*). The ellipsoid body is linearized. Coincident EL and ER activity drives synaptic depression. EPG activity is not necessary for plasticity. Similarly, in the *γ*-lobe of the mushroom body, plasticity between KCs and MBONs is driven by the coincident activity of KCs and DANs.

During exploration in VR the spontaneously learned EPG bump-cue offset is stored for many minutes (Fig 1d)^16,34,37^. To test whether visual plasticity that is optogenetically induced can also be stored for minutes, we performed a variant of this experiment where, immediately following the second EL optogenetic pairing, the fly walked in darkness for 6 minutes before returning to closed-loop VR. This protocol was as effective in updating the bump-cue offset as our standard protocol which had only a 5 second dark period following optogenetic pairing (Fig. 6e).

Intriguingly, we noticed that EL optogenetic activation reliably moved the EPG bump to the site of EL stimulation regardless of whether a visual stimulus was present (Fig. 6f, Extended Data Fig. 14a,b). This is likely due to EL-driven G*α*s signaling in EPGs, which we observed *ex vivo* (Extended Data Fig. 8d,e). Regardless of the mechanism, this means that the above experiment cannot determine whether EL activation and visually-driven ER activity alone are sufficient for plasticity, as EPGs were also activated during optogenetic pairing.

To distinguish between a learning rule for synaptic depression that requires simultaneous activity of ER, EPG and EL neurons (three-factor) and a rule that requires *only* the coactivity of ER and EL neurons (two-factor), we modified our experiment to inhibit EPG neurons, using HcKCR1, while simultaneously activating EL neurons with CsChrimson and activating ER neurons using a visual cue (Fig. 6g). First, we confirmed that photostimulating HcKCR1 suppresses EPG activity at the site where EL neurons are activated by CsChrimson (Fig. 6f, Extended Data Fig. 16a-c). This allowed us to repeat the optogenetic experiment, now testing whether EPG activity is dispensable for optogenetically induced plasticity. Pairing visual stimuli with simultaneous EL stimulation during EPG inhibition robustly drove plasticity (Fig. 6h,i). The enforced changes in EPG bump-cue offsets were consistently observed (Fig. 6i,j, Extended Data Fig. 16d,e) even though EPG neurons were locally silenced during plasticity induction. These results indicate that simultaneous EL and ER neuron activity are together sufficient conditions for inducing plasticity at ER-EPG synapses. In a control experiment, we also confirmed that silencing EPGs without activating EL neurons had no significant effect on visual tethering (Fig. 6j, Extended Data Fig. 16b-e). Thus plasticity induction can, in principle, be decoupled from the activity of the core head direction neurons. These data support a two-factor learning rule where visual activation of ER neurons simultaneous with EL neuron activity is sufficient to drive plasticity, while EPG depolarization during induction is dispensable.

## Discussion

Here we describe a novel motif for synaptic plasticity where coincidence detection at an inhibitory synapse is enabled by activity of a third octopaminergic cell type. This mechanism provides insight into the biological implementation for synaptic learning rules that were proposed for scene learning in head direction circuits over 30 years ago^14^. Specifically, we propose that ER-EPG synaptic depression occurs when visually-driven ER neuron activity is coincident with octopamine released by EL neurons onto active presynaptic terminals (Fig. 6k). EL neuron activity mirrors local EPG activity which means that ER neurons receive a copy of head direction information via this feedback loop. ER neurons express RNA for numerous octopamine receptors (Extended Data Fig. 3)^27,62–65^ which are necessary for accurate head direction tracking, and EL synaptic inputs are adjacent to ER->EPG visual presynapses, allowing signaling for plasticity induction to be spatially confined within the ellipsoid body. Our data support a model in which ER presynapses undergo depression when their own activity is correlated with octopamine receptor signaling pathways (e.g. cAMP)^74,86,87^.

Octopamine receptors are expressed in both ER and EPG neurons and EL stimulation drives measurable cAMP increases in both populations (Extended Data Fig. 8d,e). We speculate that octopamine signaling serves distinct functions on either side of the synapse. ER octopamine signaling is well suited to modulate ER-EPG synaptic strength. On the other hand, EL-driven excitation of EPGs completes a GPCR-dependent excitatory recurrent loop that could contribute to EPG bump activity maintenance and ring attractor dynamics as well as preventing EL and EPG activity bumps from becoming untethered. Further work is needed to explore these roles of octopamine signaling.

Why are there so many different octopamine receptors in ER neurons? Distinct GPCR signaling cascades may contribute to different aspects of plasticity. In the fly mushroom body, coincident increases in cAMP and presynaptic calcium drive depression of kenyon cell presynapses. Three different octopamine receptors expressed in ER4d are G*α_s_*-coupled and thus capable of increasing cAMP (Oamb, Oct*β*2R & Oct*β*3R)^82,83,88^. Oamb has the highest transcript levels making it the likely dominant source of octopamine driven cAMP. Knocking out Oamb produced deficits in the ability of the EPG bump to tether to visual cues. This manipulation is predicted to dramatically reduce cAMP production in response to EL octopamine release leading to less synaptic depression. However, Oamb can also signal through G*α_q_* signaling cascades. This pathway has been linked to synaptic potentiation in kenyon cells, so it is possible that Oamb is a critical receptor involved in both synaptic potentiation and depression. Knocking out G*α_i_*_/*o*_-coupled Oct*α*2R also caused a visual tethering deficit. One possible interpretation is that Oct*α*2R inhibition of adenylyl cyclase shortens the duration of octopamine induced cAMP transients in ER neurons to ensure that synaptic plasticity dynamics can match the timescale of navigation. In addition, *βγ* subunits from G*α_i_*_/*o*_-coupled receptors in other systems are known to directly inhibit Cav2 calcium channels. Thus, Oct*α*2R could drive immediate presynaptic depression ^89–92^. Dissecting the differential roles of these receptor signaling pathways in the biochemical coincidence detection mechanism is an important future direction. In the mushroom body, coincidence detection between presynaptic activity and G*α_s_* signaling is often attributed to Ca²⁺/calmodulin-sensitive adenylyl cyclase, Rutabaga^93–96^, but recent work shows that cAMP alone cannot drive depression of kenyon cell presynapses^86^. At hippocampal CA1 GABAergic synapses, associative depression utilizes a distinct mechanism that requires both calcineurin activity and suppression of cAMP signaling^42,97^. Therefore, ER axons could detect coincident presynaptic activity and octopamine using a variety of signaling cascades.

While octopamine has been extensively studied in the *Drosophila* nervous system it has not been previously implicated in spatial navigation. For example, octopamine has been shown to modulate odor learning by shaping synaptic plasticity in the mushroom body, often by modulating dopamine-dependent plasticity of kenyon cell cholinergic transmission^59,68,98^ or by changing the sign of spike timing dependent plasticity^99^. Beyond olfactory learning, octopamine has been found to regulate various physiological processes, including locomotion^100^, sleep^26,101^, courtship^102^, ovulation^64^, feeding^103^, and aggression^104^. Interestingly, at the *Drosophila* neuromuscular junction octopamine also alters presynaptic function either by acutely modulating glutamate release or by regulating synapse growth during development^105,106^. These results extend our understanding of the diverse synaptic actions of octopamine to include depression of GABAergic synapses in the central complex.

In this work, we focus on visual plasticity, which is thought to be mediated by ER4d and ER2a-d visual ER neurons but may also involve other ER subtypes. For example, ER3p and ER3d are predicted by connectomics to have localized visual responses^23^ and are reported to carry motor information^20,107^. However, EL neurons provide feedback onto all ER neuron subtypes (Extended Data Fig. 7) with notably higher innervation of certain subtypes (ER4d, ER3p_a, ER3p_b, ER2_c, ER3d_a & ER3d_a). Future work is needed to describe the specific visual, or non-visual, features carried by these different ER subtypes to explore the functional relevance of the differential extent of octopamine modulation for spatial processing. EL neurons also synapse onto ER1 neurons that encode wind direction and contain RNA transcripts encoding octopamine receptors (Extended Data Fig. 3e)^22^. We speculate that wind and visual input pathways engage similar plasticity mechanisms^22,34^. Interestingly, EL neurons only rarely synapse onto ER4m neurons that are tuned to the e-vector angle of polarized light^19,23^ suggesting the association between sky polarization signals and head direction may not be plastic^31^, or may engage distinct plasticity mechanisms.

It is puzzling that EL neurons carry a copy of EPG activity, when a retrograde molecule could have conveyed the same information. For example, in mammals, endocannabinoids are often released from postsynaptic neurons and drive associative synaptic depression at inhibitory presynapses^39^. However, decoupling plasticity from postsynaptic network activity may confer computational advantages to this network. First, it could allow plasticity to be regulated without affecting the ongoing network dynamics. For example, EL neurons receive strong inputs from neurons involved in sleep and circadian rhythms (ER5, ExR1) and express dopamine and serotonin receptors suggesting that EL activity, and thus synaptic plasticity, is modulated by fly behavior^38,69^ and other internal states^31,108–110^. Another potential advantage of utilizing a third neuron for coincidence detection is to counteract circuit delays. For example, the EPG bump often lags behavior (Extended Data Fig. 1c)^12,13^. Having EL neurons’ activity precede the EPG activity bump could compensate for differences in timing between movement of the head direction bump and arrival of feedforward sensory inputs^17,23,111^. Another potential benefit of a third neuron over a retrograde molecule is to reduce noise. Each EL neuron pools inputs from 6-10 EPG neurons (Extended Data Fig. 7g) meaning that an instructive signal from EL neurons is potentially less noisy than if a retrograde signal was released by each individual EPG neuron. Lastly, signaling via octopamine receptors provides more cell-type and synaptic specificity than a gaseous retrograde transmitter that readily permeates membranes. NO, for example, activates soluble guanylyl cyclases which are expressed in many cell types in addition to ER neurons^52,112^.

To enable network balance, visual inputs should undergo synaptic depression or synaptic potentiation. Such bidirectional plasticity would allow highly structured visual inputs to emerge robustly with experience^16,17,113,114^. Our data strongly constrains the learning rule that associates visual cues with head direction activity via depression of ER-EPG synapses. We predict coincident ER activity and octopamine signaling is sensed in the ER axon terminal; however, we cannot fully exclude some contribution from voltage-independent GPCR signaling in EPG neurons. In parallel to synaptic depression, potentiation of ER-EPG synapses would serve as the “forgetting” rule for visual-head direction associations. Our results are consistent with two categories of potentiation conditions: 1) Potentiation occurs when ER neurons are visually active without receiving octopamine (pre-synaptic gating)^14,16^ or 2) potentiation occurs when visual activity and octopaminergic input occur with non-coincident or incorrect timing– similar to time-window based plasticity observed in the fly mushroom body^74^, hippocampus^115,116^, and many other systems^99,117^. Notably, we think potentiation is gated by visual activity because the network’s bump-cue offset is well recalled after prolonged periods in darkness (Fig. 1g,h,j,k, Fig 6e) or following interleaved experiences of distinct visual scenes^16^. In addition, activation of EL neurons in darkness does not reset the network’s relationship to a visual cue (Extended Data Fig. 14c,d). More direct synaptic measurements in future work will be important to determine the exact synaptic algorithm and to uncover how learning is modulated by motor-locked dopamine^16,35,38^.

To our knowledge, the use of an octopamine neuron as a feedback pathway to implement coincidence detection represents a novel plasticity motif. Our findings highlight how neuromodulation of presynaptic terminals can enable coincidence detection at GABAergic synapses, where postsynaptic induction mechanisms are far less common (but see^43^). Our experiments suggest that plasticity between ER neurons and EPG neurons is induced presynaptically. This novel motif bears strong resemblance to plasticity mechanisms in other invertebrate circuits where coincident neuromodulatory drive and presynaptic activity changes synaptic strength^58,76,118^. The similarities between the synaptic mechanisms we have uncovered in the head direction network and plasticity in the *γ*-lobe of the mushroom body are particularly striking (Fig. 6k). In the mushroom body, synaptic plasticity for odor learning is instructed by dopamine neurons (DANs) that innervate specific compartments of odor-driven kenyon cell axon terminals^75,119,120^. Coincident kenyon cell activity and DAN input are necessary and sufficient for depression of the excitatory synapses between kenyon cells and mushroom body output neurons (MBONs). An important topic for future work will be to uncover whether mechanisms of plasticity expression are also shared between these circuits.

Neuromodulation of presynaptic terminals by monoamines such as dopamine^121^, serotonin^122^, and norepinephrine^123–125^ is also widespread in vertebrates. What distinguishes the synaptic plasticity we have discovered in the ellipsoid body from previous reports is that here the neuromodulatory cell carries a near copy of the postsynaptic neuron’s activity. By contrast, in other systems monoaminergic neurons that instruct plasticity often carry information about valence, novelty, or movement^59,126–128^. It appears evolution has co-opted a motif that is common in reinforcement-based learning to solve the problem of forming an accurate internal representation of space. Given the widespread expression of GPCRs for biogenic amines on presynaptic terminals, we speculate that similar presynaptic neuromodulation motifs could implement diverse learning algorithms in many other circuits.

Our experiments provide insight into the synaptic mechanisms by which head direction circuits anchor their representations to the sensory world. Importantly, recent theoretical work argues that the principles governing unsupervised spatial learning, such as the tethering of head direction cells to landmarks, generalizes to all relational memory^129–131^. Therefore, the synaptic principles engaged by head direction cells can inform not only spatial learning performed by grid and place cells but also non-spatial relational learning. This work lays a foundation for future investigations into the synaptic and biochemical substrates of unsupervised learning algorithms and should motivate exploration into the role of neuromodulators for associative plasticity beyond reinforcement learning contexts.

## Figures and legends

**Extended Data Figure 1.**
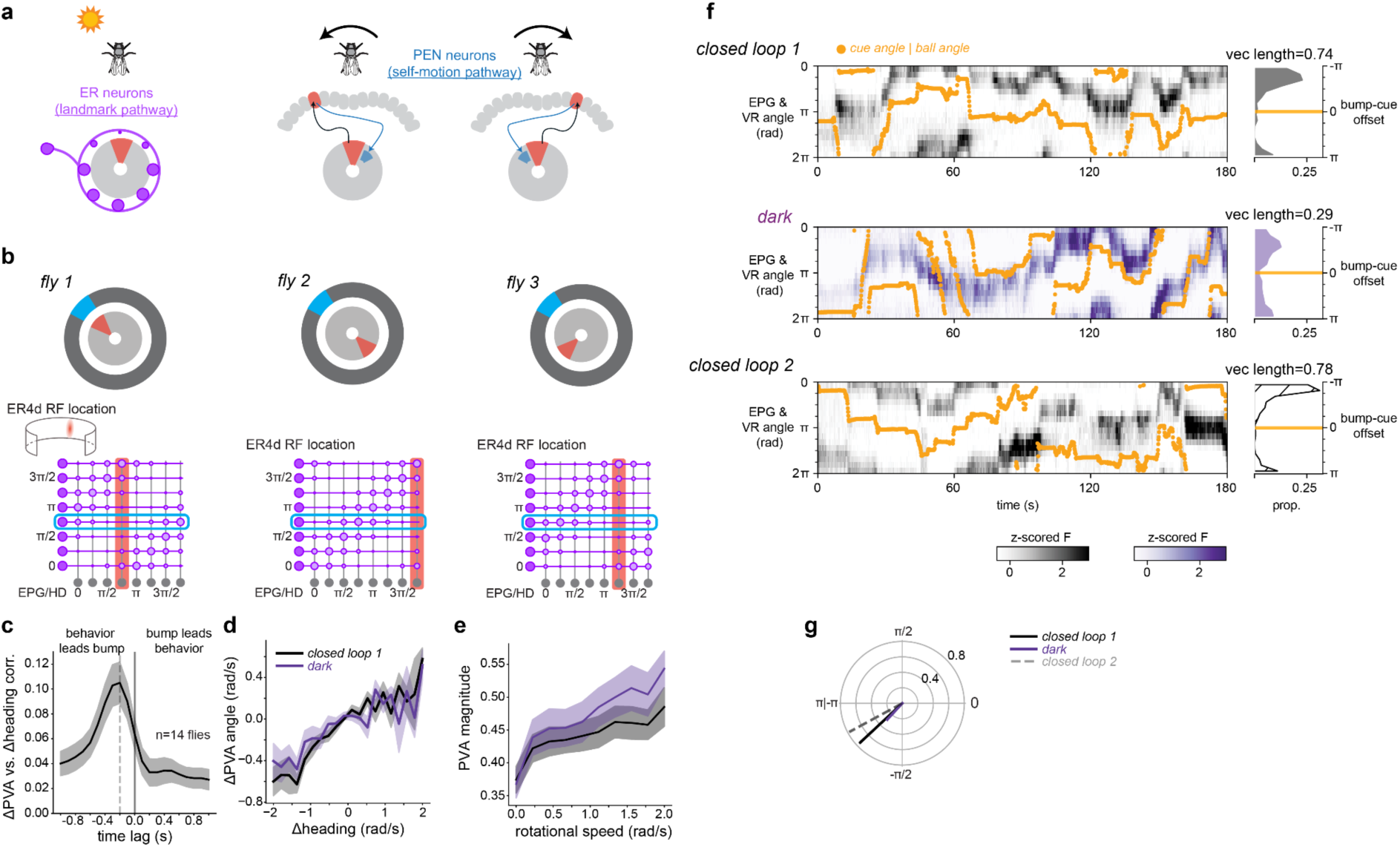
EPG bump dynamics during behavior and across dark periods. a) Schematic illustrating the landmark and self-motion pathways that collectively update the EPG bump position. Landmark signals (visual and other modalities) enter the circuit via ER neurons, while self-motion signals are conveyed via PEN neurons that serve to spin the bump position as the animal turns to the left or right. When external sensory cues are present both pathways are engaged. Without external sensory cues, such as in darkness, head direction estimation relies solely on integrating inputs to the self-motion pathway. b) Schematic illustrating hypothesized ER-EPG synaptic strengths. *Top:* 3 example bump-cue offsets. *Bottom:* corresponding synaptic strength matrices. Connections between EPG neurons (columns, gray), sorted by angle in the ellipsoid body, and ER4d neurons (rows, purple), sorted by the azimuthal angle of their visual receptive field, are shown. Strong synapses are indicated by large circles, and weak synapses are indicated by small circles. The active EPG (red) and ER4d (cyan) neurons are highlighted. The diagonal location of the trough of weak synapses determines the bump-cue offset. c) EPG bump movement lags changes in behavior. Correlation coefficient between the movement of the EPG bump (change in PVA angle) and the change in heading in VR as a function of the shift applied to the heading timeseries. Correlation peaks at a lag of 200 milliseconds (dotted line). d) Change in PVA angle (bump velocity) plotted as a function of change in heading (fly rotational velocity) for closed loop and dark sessions. EPG timeseries is shifted by 200 milliseconds to maximize correlation. e) PVA magnitude as a function of rotational speed for closed loop and dark sessions. For panels c-e, data is plotted as mean ± sem across flies. f-g) Additional example fly imaging data as in Fig. 1g-h

**Extended Data Figure 2.**
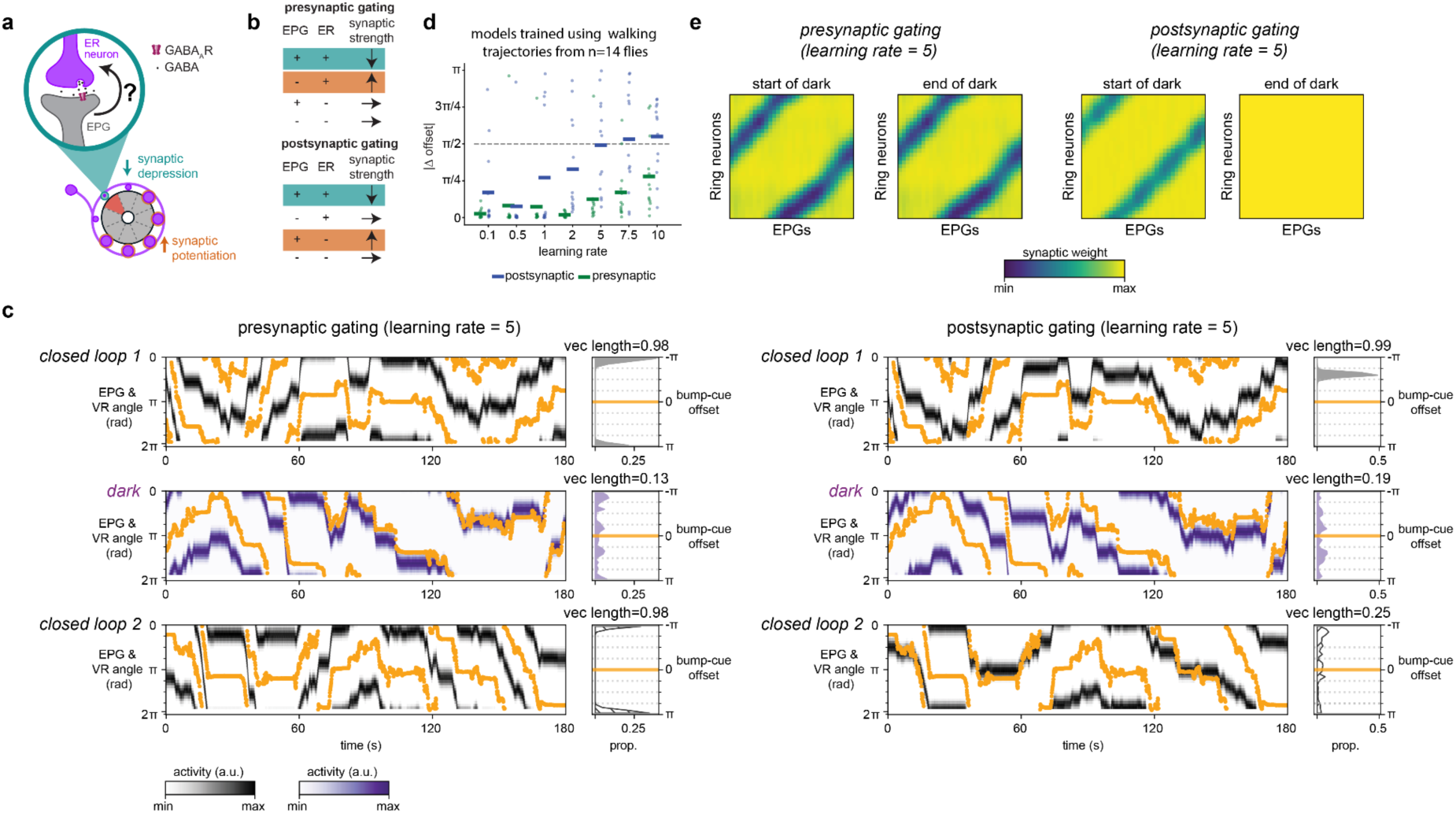
Simulated network dynamics across dark periods. a) Schematic illustrating plasticity model. Active ER-EPG synapses that coincide with the EPG bump depress; other synapses potentiate. b) Summary of synaptic learning rules under the “presynaptic gating” and “postsynaptic gating” models^16^. In presynaptically gated models, an ER→EPG synapse changes only when the ER neuron is active; in postsynaptically gated models, the synapse changes whenever the EPG neuron is active. Coincident or non-coincident activity determines the direction of weight change. Note that these terms describe which activity is required for plasticity, not where plasticity is induced. Depression conditions highlighted in teal. Potentiation conditions highlighted in orange. c) Simulated network activity using walking trajectories from behavioral data in Fig. 1g (left-presynaptically gated learning rule, right-postsynaptically gated learning rule) d) A presynaptically gated learning rule results in smaller changes to bump-cue offsets across dark periods for a range of learning rates. Change in simulated bump-cue offset across the two closed loop sessions using the same walking trajectories as in Fig. 1i-k. Each dot is a single model run using the walking data from a single fly. e) Dark periods do not affect synaptic structure in presynaptically gated models but abolish synaptic structure in postsynaptically gated models. ER-EPG synaptic strength matrices at the beginning and end of the dark session for the simulations shown in [c].

**Extended Data Figure 3.**
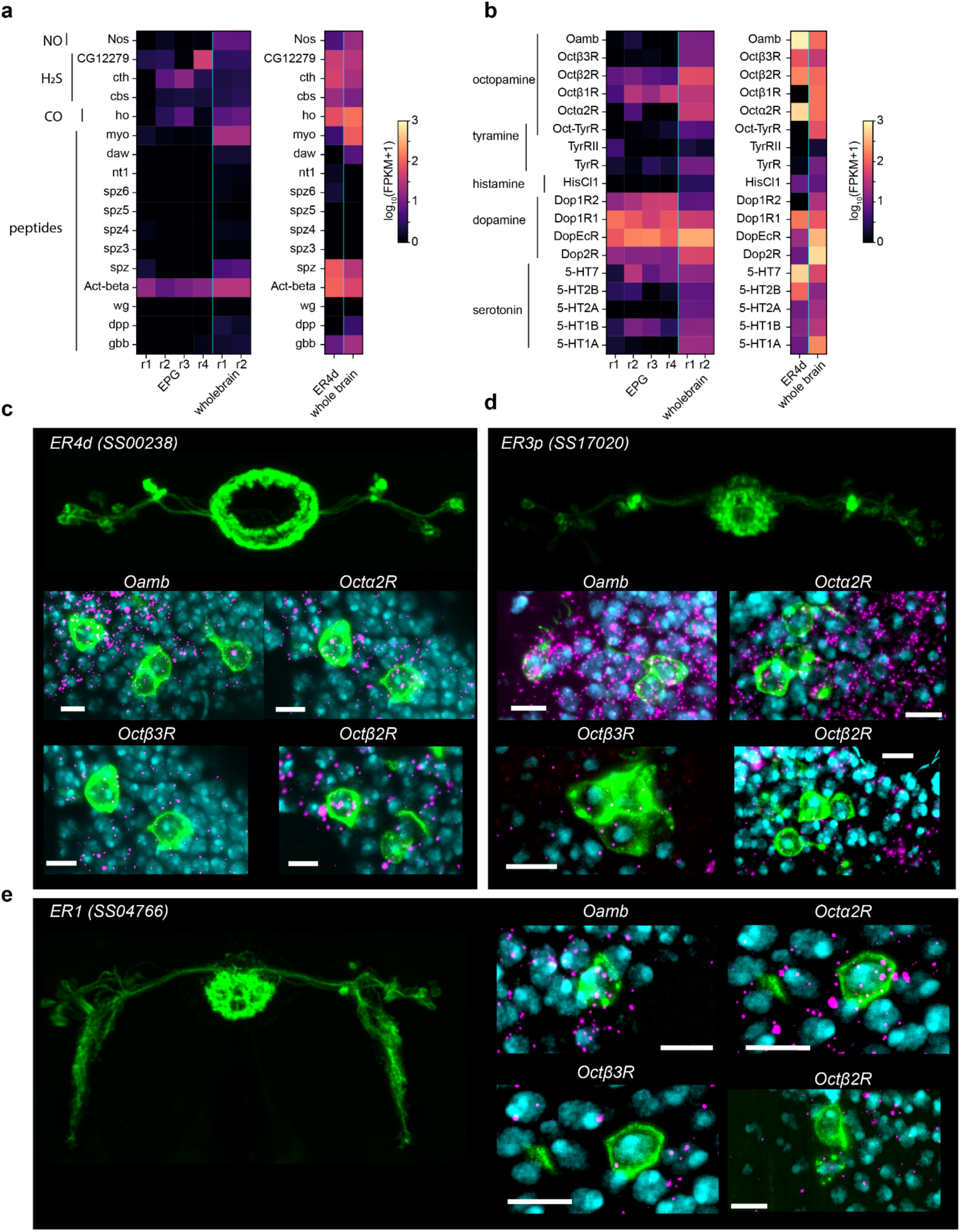
mRNA transcript measurements for retrograde signals and monoamine receptors in EPG and ER neuron subtypes. a) Analysis of RNA sequencing data from a previously published dataset of EPG and ER4d neurons^27^ showing genes related to potential retrograde signaling molecules. Enzymes for gaseous transmitter production: nitrous oxide (NO), hydrogen sulfide (H_2_S) and carbon monoxide (CO). Peptides with potential retrograde signaling functions are also listed. Log FPKM values from each cell type with a whole brain reference. EPG data shows multiple replicates from bulk sequencing. ER4d data is averaged low cell sequencing values. b) Monoaminergic receptors RNA transcripts from EPG, ER4d, and whole brain as in [a]. c) Example images from EASI-FISH showing octopamine receptor mRNA transcripts in ER4d somata (anti-GFP-green, RNA-magenta, DAPI-cyan, see Methods). Images are partial z-stack maximum intensity projections through example cell bodies. Octα2R and Octβ3R data are from different channels in the same field of view. Scale bars are 10 μm. Max projection image of full neuron morphology shown for reference. Raw ER Split Gal4 images courtesy of the FlyLight Project Team, Janelia Research Campus, HHMI (CC BY 4.0). d) Same as [c] for ER3p neurons. e) Same as [c] for ER1 neurons.

**Extended Data Figure 4.**
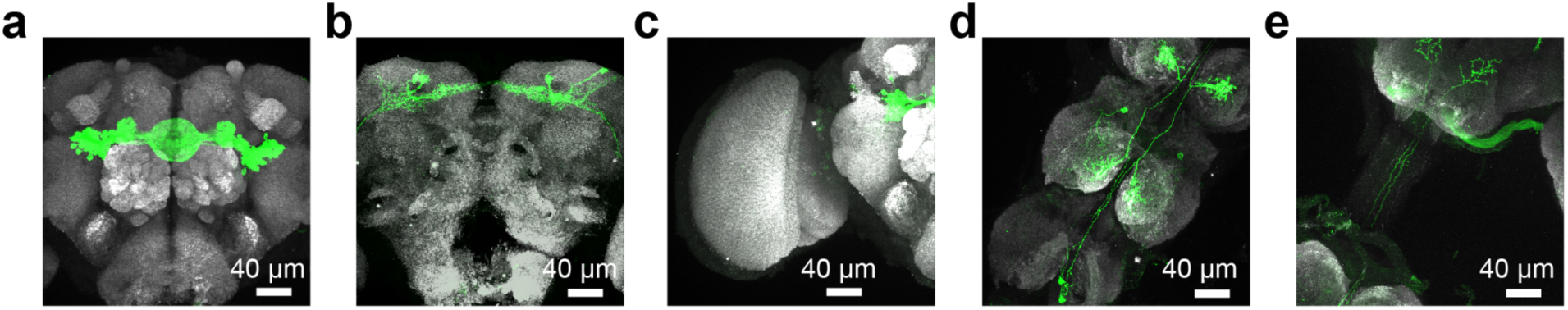
Characterization of ER split-Gal4 driver line. a-e) Adult brains with mCD8::GFP expression (anti-GFP, green) driven by the ER neuron split Gal4 (R56H10-p65.AD {VK00027}, R20A02-DBD {attp2}) with Bruchpilot staining (nc82, grey). Confocal images are partial Z-stack maximum intensity projections. Expression in the central brain is largely restricted to ER neurons. A small number of neurons are labeled in the superior protocerebrum, optic lobe, and ventral nerve cord. a) Anterior brain b) Posterior brain c) Optic lobe d-e) Ventral nerve cord

**Extended Data Figure 5.**
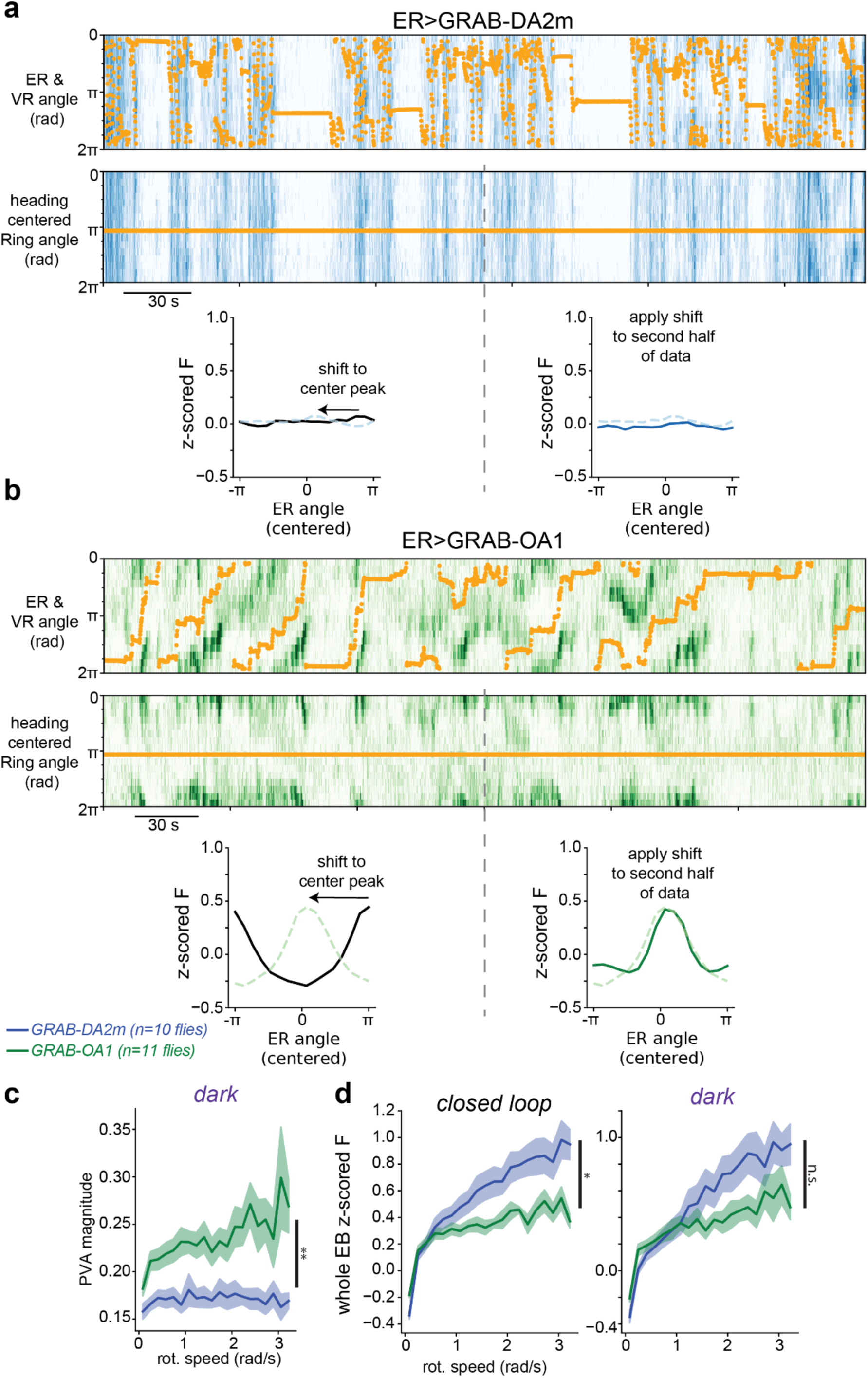
Schematics and further analyses of GRAB sensor imaging. a-b) Illustration of analysis in Fig. 2h. a) *Top:* Example 6 minute ER>GRAB-DA2m closed loop imaging session for an additional example fly. *Middle:* Each column in the heatmap is circularly shifted such that heading makes a straight line. *Bottom left:* For the first half of the imaging session, a temporal average is taken from the heading-aligned data. The spatial shift required to center the peak of that average trace is then calculated. *Bottom right:* The shift is applied to the temporal average of the second half of the data, resulting in a cross-validated centering of the activity bump. b) Same as [a] for the example ER>GRAB-OA1 session in Fig. 2g. c) Same as Fig. 2i but for dark sessions. Mixed effects ANOVA on log-transformed values: GRAB sensor main effect: F=12.8, p=0.002. d) Whole ellipsoid body GRAB fluorescence as a function of flies’ rotational speed. GRAB-DA2m fluorescence (blue) shows a larger increase as a function of rotational speed than GRAB-OA1 fluorescence (green) in both closed loop (left) and dark sessions (right). Mixed effects ANOVA on log-transformed values: closed-loop – GRAB sensor main effect F=6.45 p=0.023; dark – GRAB sensor main effect F=4.24 p=0.078.

**Extended Data Figure 6.**
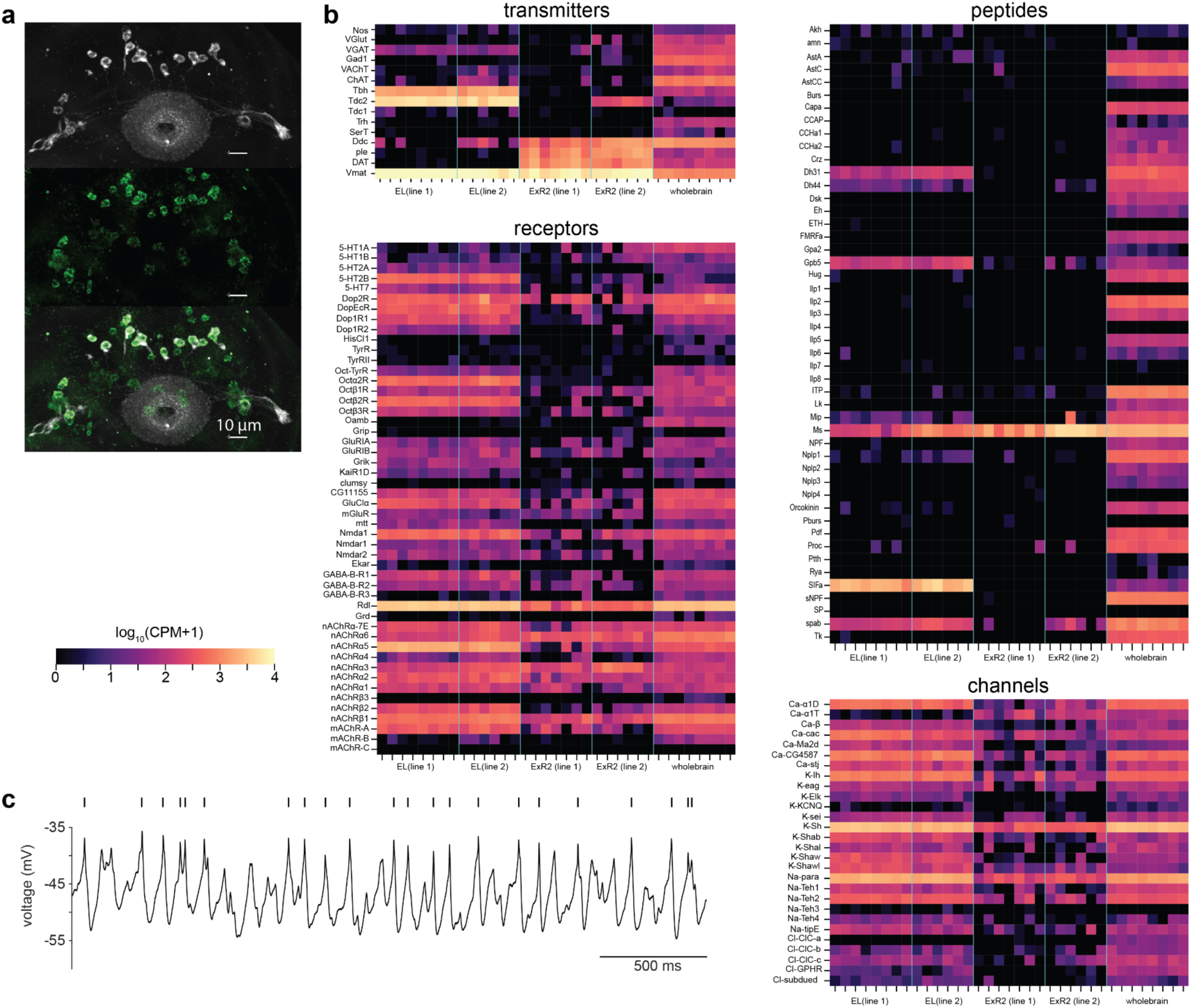
Characterization of EL and ExR2 transmitter, receptor, peptides, and channel expression. a) FISH showing TβH RNA in EL neurons. *Top:* EL Line 1 (flylight number SS00027, see Methods for full genotype) HaloTag expression (greyscale). *Middle:* TβH RNA (green). *Bottom:* Merged. All EL somata are double labeled. b) Select results from bulk RNA sequencing of EL Line 1 (SS00027), EL Line 2 (SS50573), ExR2 Line 1 (SS70424) and ExR2 Line 2 (MB233B). Log CPM values are shown for all replicates and a whole brain reference. c) Whole-cell current-clamp recording from an EL neuron demonstrating that EL neurons are intrinsically capable of generating action potentials. Putative action potentials are annotated with tick marks.

**Extended Data Figure 7.**
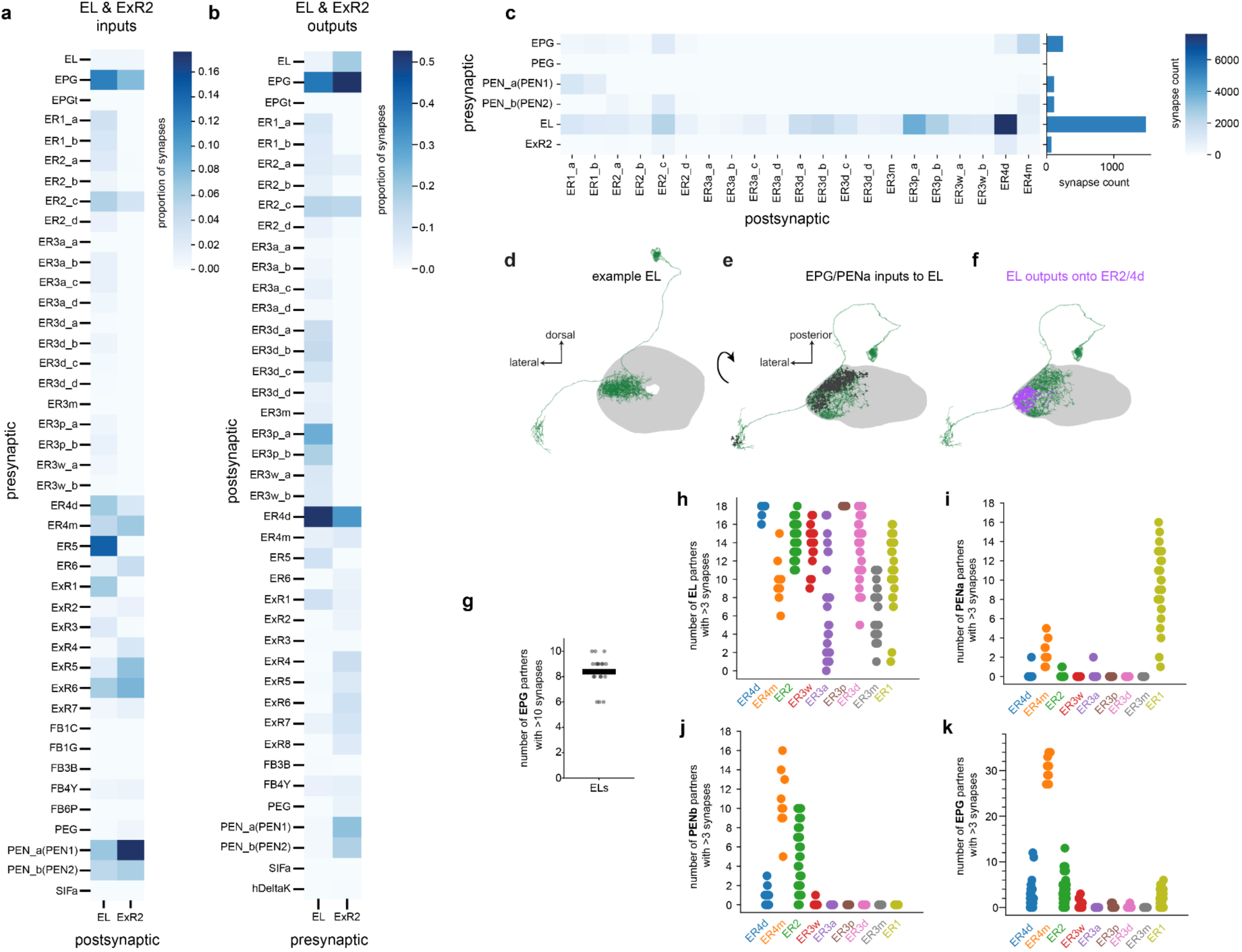
Analysis of the Hemibrain connectome exploring input and output connectivity of EL and ExR2. a) All synaptic inputs to EL and ExR2 neurons with at least 3 synapses. Synapse counts are scaled to show proportion of EL and ExR2 postsynapses to account for differences in total synapse number between the two populations. b) All synaptic outputs from EL and ExR2 neurons with at least 3 synapses. Counts are normalized to proportions of presynapses. c) EL provides the largest source of columnar neuron feedback onto ER neurons. EL provides the strongest feedback onto ER4d and ER3p neurons. *Left:* Synapse counts between presynaptic columnar neurons (ExR2 also included) and postsynaptic sensory ER neuron subtypes. *Right:* Summed synapse counts across all ER neurons. EL overwhelmingly makes the most synapses onto ER neurons. Note that polarized light encoding ER4m neurons receive stronger EPG input than EL input which likely represents a special case. Mechanosensory coding ER1 neurons also receive relatively strong PENa input. d-f) EL neurons have regions of their arbors in the ellipsoid body with distinct synaptic partners. See Fig. 3f & g for analysis of proximity of pre and postsynaptic specializations within ER4d and ER2 neurons where we hypothesize coincidence detection occurs. d) Reconstruction of a single EL neuron projecting along the approximate anterior-posterior axis to visualize the donut shape of the ellipsoid body. e) Same EL neuron as in [d] projected along the dorsal-ventral axis to view the ellipsoid body cross section. Overlaid postsynapses from EPG and PENa inputs (black). f) Same as [e] with overlaid presynapses onto ER4d and ER2 neurons (magenta). g) EL neurons pool inputs from multiple EPG neurons. The number of EPG presynaptic partners with greater than 10 synapses for each EL neuron. h) Strip chart showing the number of EL neuron synaptic partners with greater than 3 synapses that inputs to each ER neuron subtype. Each dot is one ER neuron instance. i-k) Same as in [h] but for PENa [i], PENb [j] and EPG [k]

**Extended Data Figure 8.**
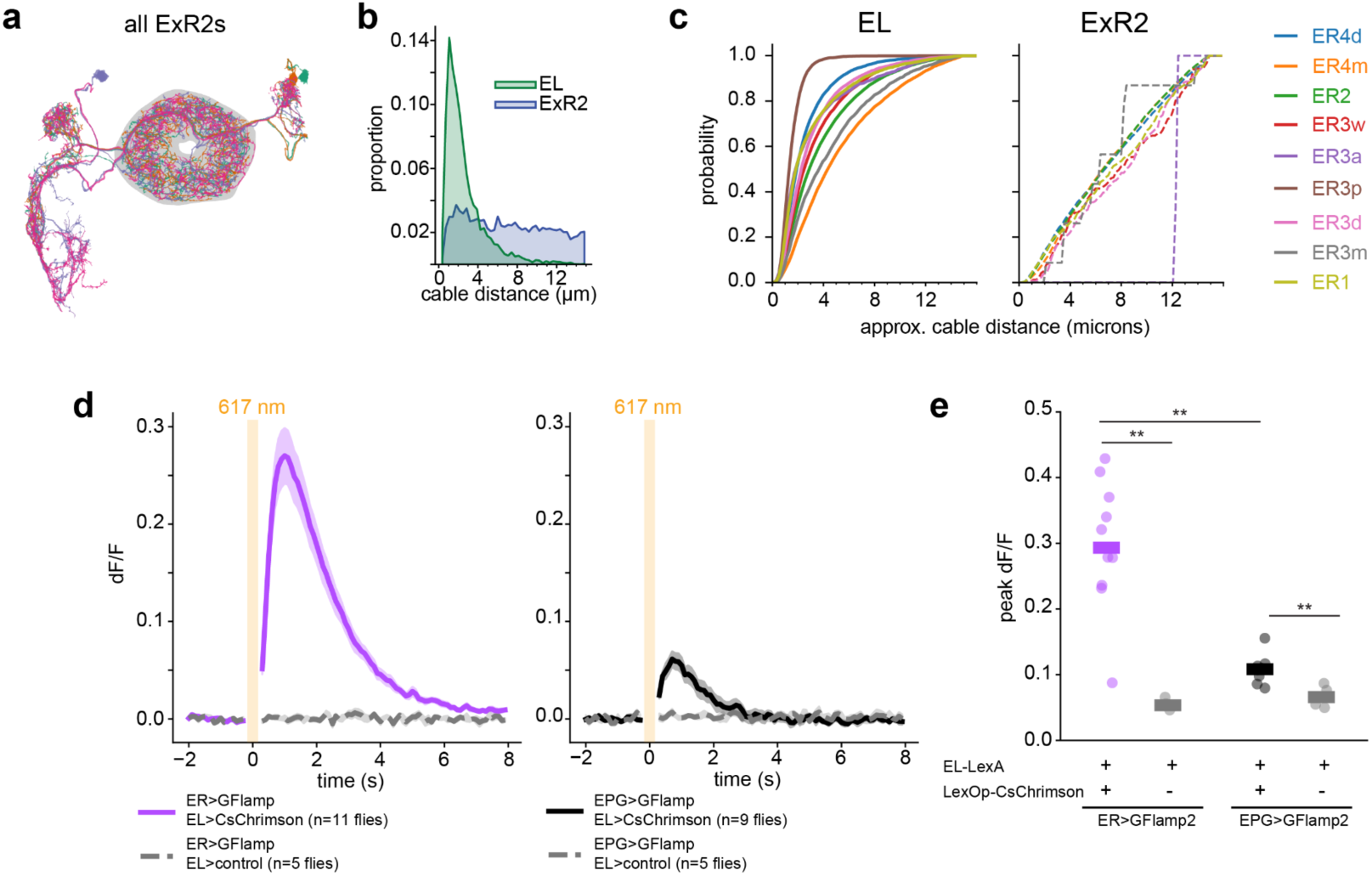
Analysis of ER-columnar neuron connectivity from Hemibrain connectome and EL-driven cAMP signaling in ER and EPG neurons. a) EM reconstruction of all ExR2 neuron arbors within the hemibrain volume^30,132^. b) Cable distance from ER2/4d-EPG presynapses to EL-ER2/4d postsynapses (green, same as Fig. 3g) and for comparison the same analysis for ExR2 is shown in blue with cable distance from ER-EPG presynapses to ExR2-ER postsynapses. c) *Left-EL* cable distance analysis from Fig. 3g was repeated for different ER neuron subtypes and shown here as a cumulative histogram. EL contacts are particularly close to ER3p-EPG synapses and ER4d-EPG synapses. *Right-* Cable distance analysis for ExR2, cumulative histogram shows distance from ER-EPG synapses to closest ExR2-ER synapse for different ER neuron subtypes. d) ER (magenta) and EPG (black) neurons display increases in cAMP (GFlamp2^133^) in response to optogenetic EL stimulation (CsChrimson, 50 msec stimulation with 617 nm LED) *ex vivo*. e) ER and EPG neurons show significant cAMP increases in response to EL stimulation. The cAMP increase in ER neurons is significantly larger than EPGs. Average peak GFlamp fluorescence for each fly. Thick lines show across fly median. Mann-Whitney U tests (holm-corrected p-values): ER GFlamp with vs without EL>CsChrimson U=55 p=0.001, EPG GFlamp with vs without EL>CsChrimson U=43 p=0.004, ER vs EPG GFlamp U=92 p=0.003

**Extended Data Figure 9.**
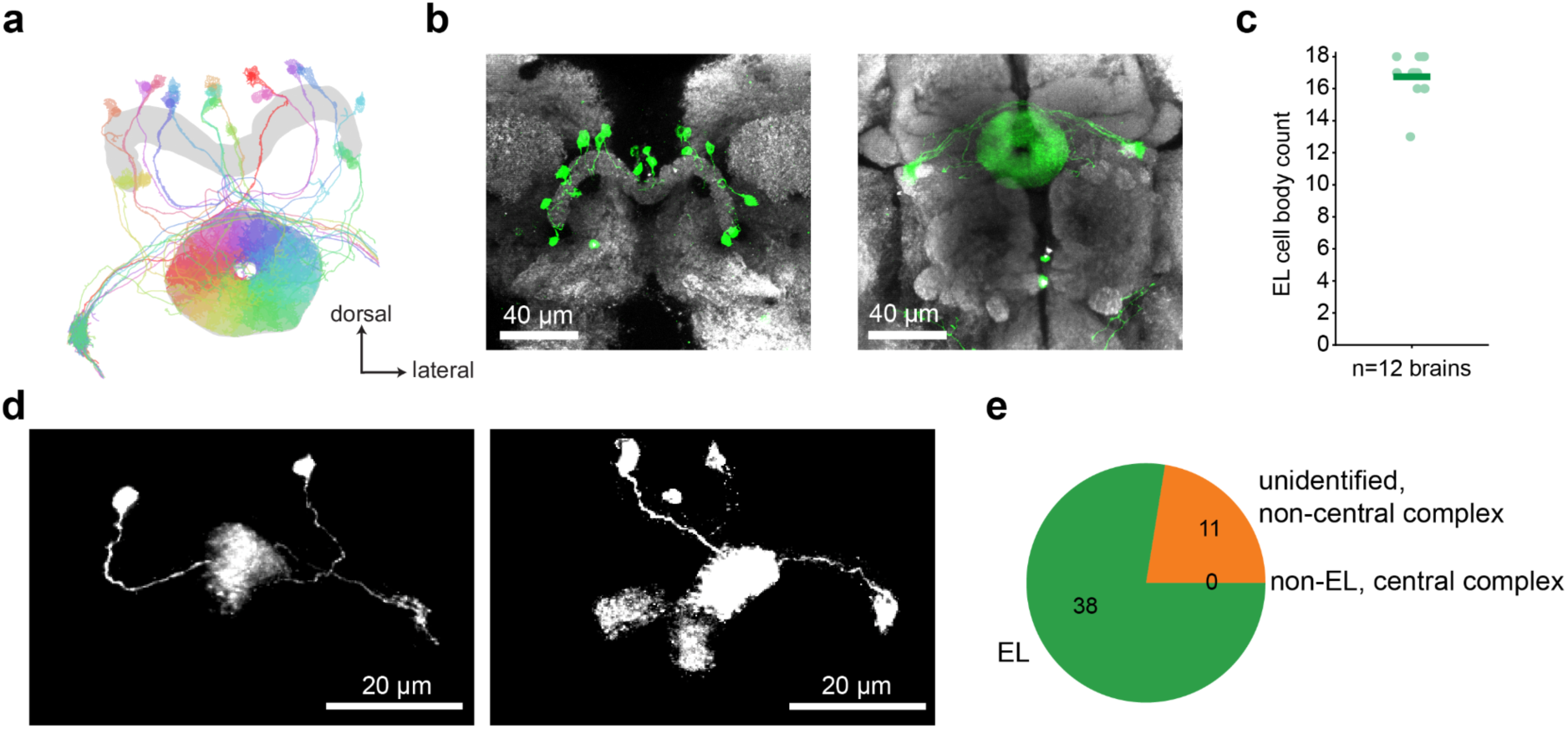
Characterization of EL LexA driver line (R18B05-LexA) a) EM reconstruction of all EL neurons showing key elements of their anatomy^30,132^. Cell bodies are near the protocerebral bridge. EL neurons, however, do not arborize in the protocerebral bridge. Their processes tile the wedges of the ellipsoid body and they additionally arborize in the gall surround. b) Confocal images of an adult brain with mCD8::GFP expression driven by the R18B05-LexA driver line (left: protocerebral bridge, right: ellipsoid body; anti-GFP-green; anti-Bruchpilot/nc82-grey). Images are partial z-stack maximum intensity projections. Scale bars are 40μm. This driver line is specific for EL neurons in the central complex. A small number of unidentified putative ascending neurons are labeled as well. c) Number of EL neurons labeled by R18B05-LexA per brain. Each dot represents the number of GFP-labeled EL cell bodies from one brain. The green bar shows the mean cells per brain (16.75). Connectomics data suggests there are 17-18 EL neurons per fly brain^30,132,134–136^. d) MultiColor FlpOut (MCFO)^137^ anatomical characterization of the R18B05-LexA driver line. Stochastically labeled example neurons expressing HA-tagged smGFP reporter from two different flies. *Left*: 2 EL neurons. Right: 3 EL neurons. Both images are maximum intensity projections from the bulb to the protocerebral bridge. e): Pie chart summarizing R18B05-LexA MCFO data. All neurons with arbors in the ellipsoid body were identified as EL neurons. 77.6% of all cells labeled were identified as EL neurons, 22.4% of cells were non-EL, non-columnar neurons, and 0% were non-EL, columnar neurons. Importantly we never identified cells with EPG, PEG, PENa or PENb morphology. N flies = 7, N cells = 49.

**Extended Data Figure 10.**
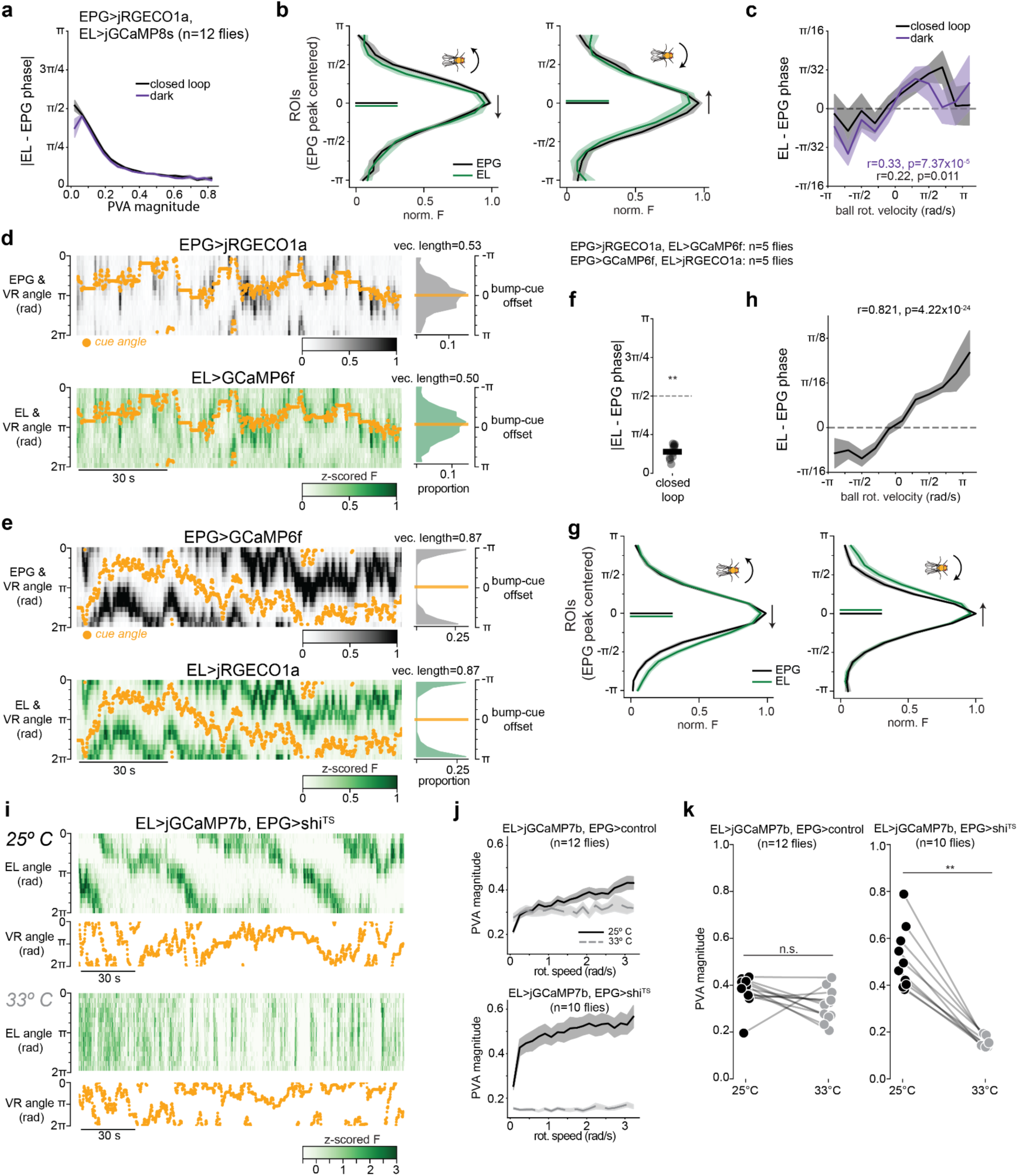
Co-imaging of EPG and EL activity with additional indicator combinations and EL bump activity requires EPG input. a) Analysis of EL and EPG bump phase during closed loop VR dual imaging session (same data set as in Fig. 3h & i). Large deviations in EL and EPG bump phase only occur when bump magnitudes are small. The absolute value of the difference in EL and EPG bump phase is plotted against the average EL and EPG PVA magnitude in closed loop (black) or darkness (purple). Data is plotted as mean ± sem across flies. b) The EL bump phase slightly leads the EPG bump phase. Data from both EL (green) and EPG (black) is centered by the EPG PVA angle during rapid counterclockwise (left) or clockwise (right) turns. The direction of bump motion during the turns is indicated by the arrows. Data is plotted as mean ± sem across flies. Inset lines indicate the PVA angle of the plotted data. c) The difference in EL and EPG bump phase is plotted against the fly’s rotational velocity in closed loop (black) or darkness (purple). Inset statistics show Spearman rank correlation. Data is plotted as mean ± sem across flies. d-e) Example data of simultaneous 2P imaging of EL and EPG during closed loop VR as in Fig. 2 for two additional calcium indicator combinations. For all flies indicators are driven by EL-Gal4 and EPG-LexA. For panels [f-h], data from both color combinations are combined. f) EL and EPG activity bumps have similar phases with these additional indicator combinations. Average absolute differences in bump angle (PVA angle) are plotted for each fly from closed loop sessions. Differences are significantly less than chance (*π*/2) by Wilcoxon signed-rank test: W=0 p=0.002 g) Data as in [b] for flies with indicator combinations shown in [d-e] h) Data as in [c] for flies with indicator combinations shown in [d-e]. Spearman rank correlation shown in panel. i-k) EL “bump” activity requires EPG input. i) Example EL imaging data from a single fly walking in darkness expressing shi^TS^ in EPG neurons. At permissive temperatures (top) EL neurons display an activity bump that roughly tracks the fly’s movements. At restrictive temperatures (bottom) the EL activity bump is abolished. j) PVA magnitude as a function of rotational speed for permissive (black) and restrictive (gray) temperatures. (top-EPG>control, bottom-EPG>shi^TS^). Shaded regions show across fly mean +/− sem. k) Average PVA magnitude at permissive and restrictive temperature for each fly matched for rotational speed. Lines connect repeated measurements from the same fly. EPG silencing reduces PVA magnitude. Increased temperature alone does not affect PVA magnitude. Wilcoxon signed rank tests comparing permissive and restrictive temperature PVA magnitudes (holm-corrected p-values): EPG>control: W=16 p=0.077 EPG>shi^TS^: W=0 p=0.004. Mann-Whitney U test comparing restrictive temperature PVA magnitude across genotypes: U=0 p=8.74×10^-5^.

**Extended Data Figure 11.**
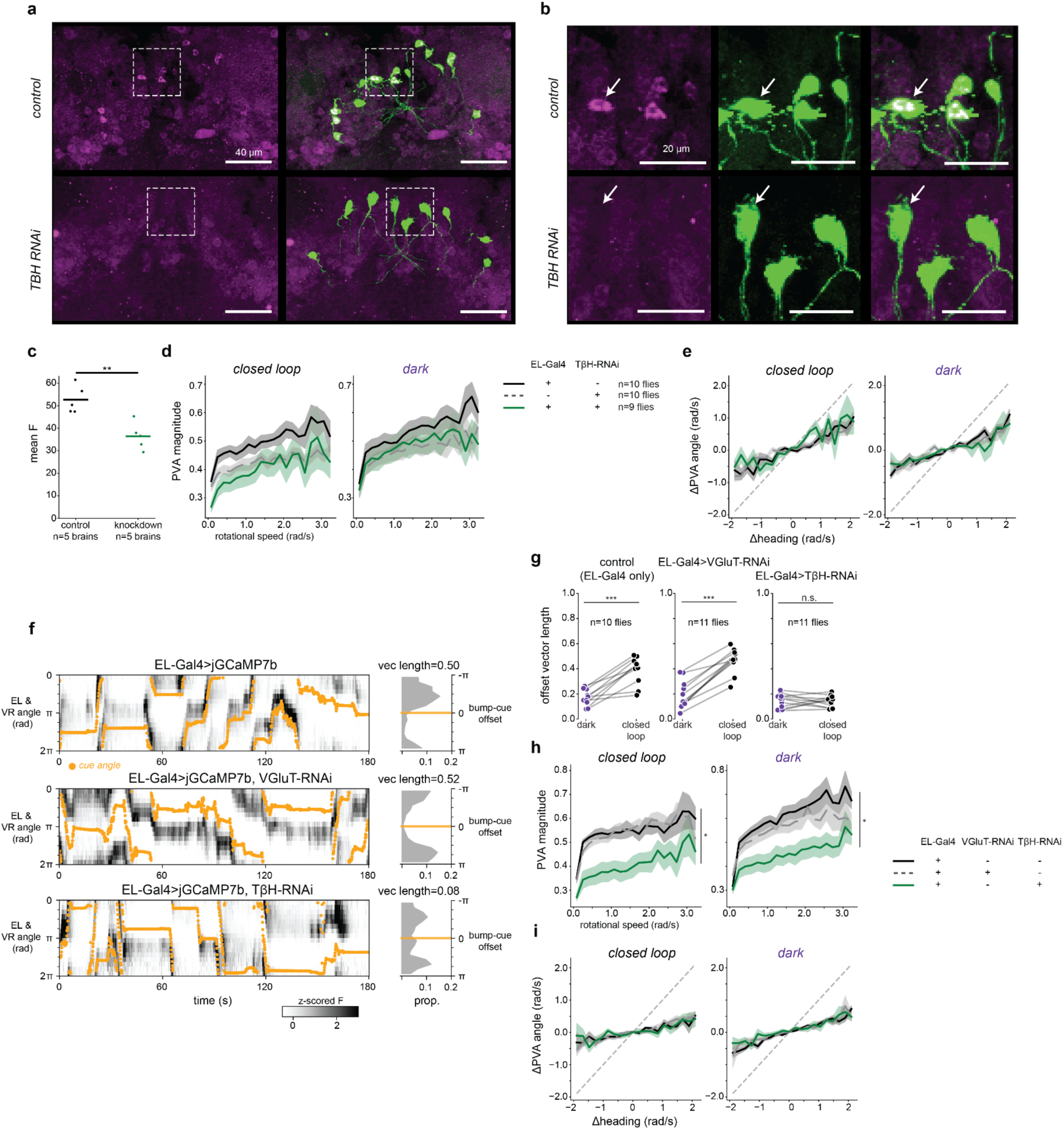
Additional quantification of the effects of knocking down TβH in EL neurons. a) Confocal images of adult brains showing staining for octopamine alone (anti-octopamine, magenta, left) or merged with EL cell bodies labeled by mCD8::GFP using an EL Split-GAL4 driver line (anti-GFP, green, right). *Top*: GFP only control. *Bottom*: TβH RNAi. Images are maximum intensity projections. Scale bars are 40 μm. Dotted boxes show insets in [b] b) Confocal images showing the cell bodies from [a]. Staining for octopamine (anti-octopamine, left), mCD8::GFP (anti-GFP, green, middle), and merged (right). *Top*: GFP only control. *Bottom:* TβH RNAi. Images are maximum intensity projections. Scale bars are 20μm. c) Expressing TβH RNAi in EL neurons decreases octopamine antibody staining intensity. Mean anti-octopamine fluorescence within GFP-labeled EL cell bodies in the control brains (black) versus the TβH knockdown brains (green). Each dot is a single brain (N control = 5 flies, N TβH knockdown = 5 flies). Mann Whitney U=0 p = 0.008 d) PVA magnitude as a function of rotational speed for data in Fig. 4c. Data is plotted as mean ± sem across flies. Kruskal-Wallis test for difference across genotypes: closed loop – H=5.98 p=0.050; dark – H=2.25 p=0.325. e) Change in PVA angle as a function of rotational velocity for data in Fig. 4c. Data is plotted as mean ± sem across flies. Linear model including genotype, dark vs closed loop, and genotype x rotational speed as predictors. Wald test for significant genotype coefficients: F(4,1166)=1.16 p=0.325. Wald test for significant genotype x rotational speed interaction: F(4,1166)=1.70 p=0.147 f-i) TβH RNAi experiments were repeated using a more specific EL-split Gal4 line while also imaging calcium from EL neurons. In addition to an EL-Gal4 only control we performed RNAi against the vesicular glutamate transporter (VGluT), a required protein for vesicular packaging of glutamate. This condition serves as an additional control for nonspecific effects of RNAi. f) Example data from closed loop imaging sessions from each genotype as in Fig. 4b. g) Average offset vector lengths for each genotype. Mixed effect ANOVA on logit transformed vector lengths: main effect of genotype: F=15.6 p=2.50×10^-5^, main effect of dark vs closed loop: F=68.9 p=3.76×10^-9^, interaction: F=17.9 p=8.68×10^-6^. Posthoc paired t-tests for dark vs closed loop (p-values holm corrected): Gal4 only control: t=5.85 p=4.87×10^-4^, EL-Gal4>VGluT-RNAi: t=9.00 p=1.2×10^-5^, EL-Gal4>TβH-RNAi: t=0.122 p=0.905. Posthoc unpaired t-test for difference in closed loop vector length between genotypes (p-values holm corrected): EL-Gal4 only vs EL-Gal4>VGluT-RNAi: t=1.66 p=0.457, EL-Gal4 only vs EL-Gal4>TβH-RNAi: t=6.66 p=1.83×10^-5^, EL-Gal4>VGluT-RNAi vs EL-Gal4>TβH-RNAi: t=9.66 p=3.39×10^-8^. h) PVA magnitude as a function of rotational speed. Kruskal-Wallis test for difference across genotypes: closed loop – H=9.05 p=0.011, dark – H=9.06 p=0.011. Posthoc Mann-Whitney U (p-values holm corrected): EL-Gal4 only vs EL-Gal4>VGluT-RNAi: closed loop – U=58 p=0.860 dark – U=73 p=0.218, EL-Gal4 only vs EL-Gal4>TβH-RNAi: closed loop – U=94 p=0.020 dark – U=96 p=0.013, EL-Gal4>VGluT-RNAi vs EL-Gal4>TβH-RNAi: closed loop – U=97 p=0.036 dark – U=89 p=0.132. i) Change in PVA angle vs rotational speed. Linear model including genotype, dark vs closed loop, and genotype x rotational speed as predictors. Wald test for significant genotype coefficients: F(4, 1328)=0.79 p=0.529. Wald test for significant genotype x rotational speed interaction: F(4,1328)=1.2 p=0.311.

**Extended Data Figure 12.**
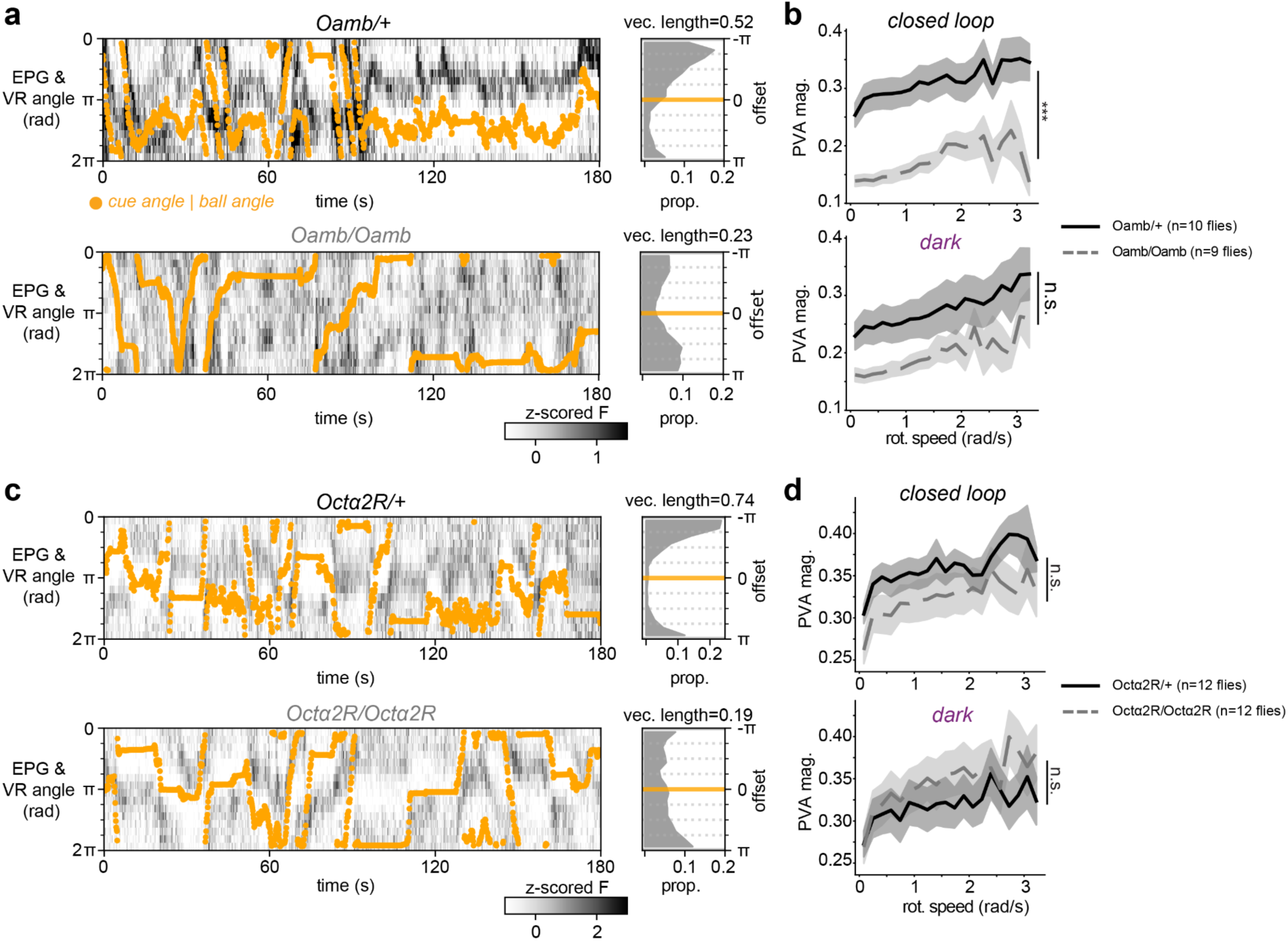
Example sessions and PVA quantifications for octopamine receptor knockout experiments. a) Example EPG imaging data in closed loop from heterozygous control fly (top: *Oamb/+)* and experimental fly (*bottom:* Oamb/Oamb). Data plotted as in Fig. 1g. b) PVA magnitude as a function of rotational speed for *Oamb/+* and Oamb/Oamb genotypes. Data is plotted as mean ± sem across flies. Mixed effects ANOVA on log-transformed values: genotype main effect: closed loop – F(1,17)=27.6 p=6.52×10^-5^, dark – F(1,17)=3.83 p=0.067 c) Example EPG imaging data in closed loop from heterozygous control fly (top: *Octα2R/+)* and experimental fly (*bottom: Octα2R*/*Octα2R*). Data plotted as in Fig. 1g. d) PVA magnitude as a function of rotational speed. Data is plotted as mean ± sem across flies. Mixed effects ANOVA on log-transformed values: genotype main effect: closed loop – F(1,23)=1.99 p=0.17, dark – F(1,22)=0.261 p=0.614.

**Extended Data Figure 13.**
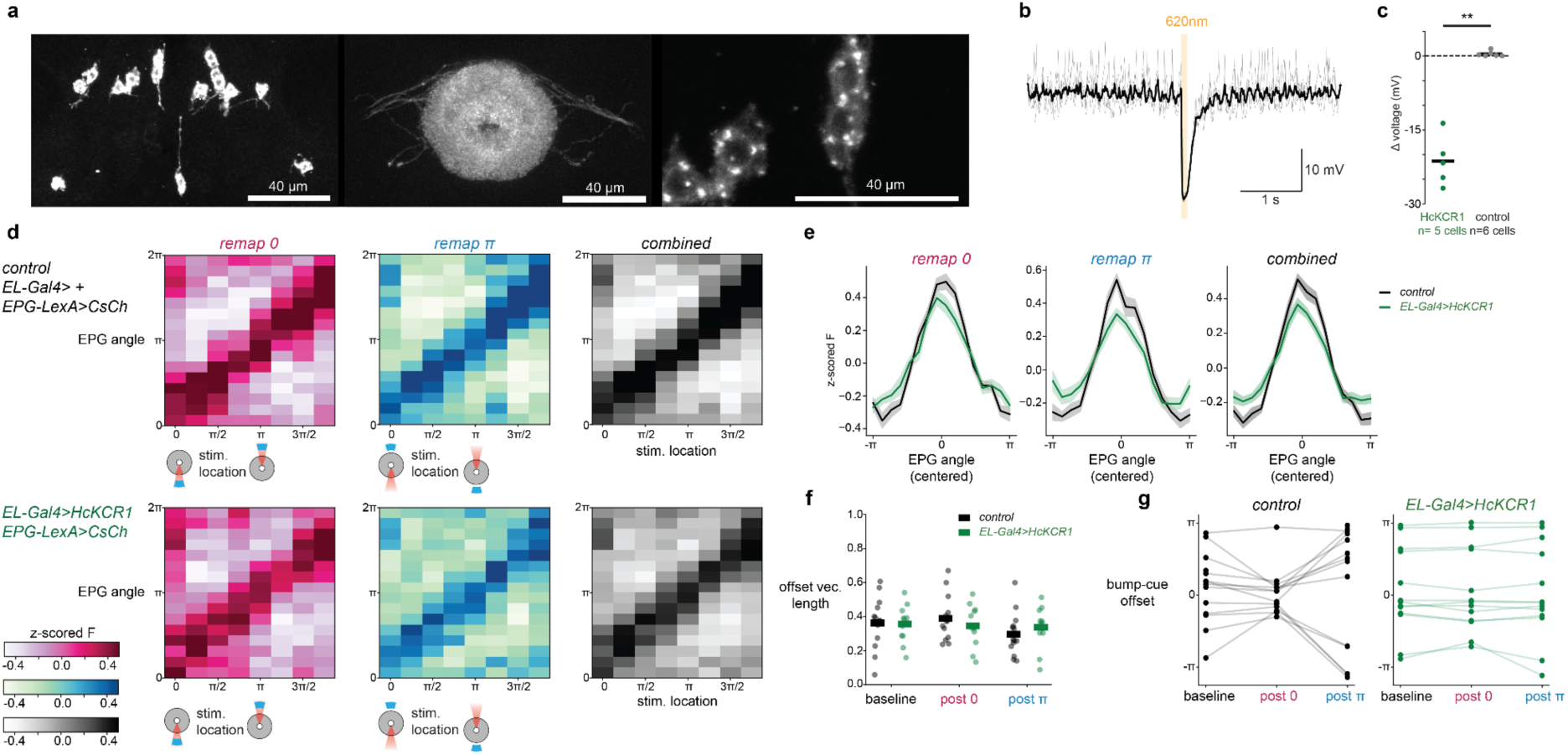
Validation of optogenetic approach to stimulating EPG neurons while silencing EL neurons. Panels in this figure pertain to data included in Fig. 5. a) Adult brain with HcKCR1::V5 expression (anti-V5, gray) driven by the EL split Gal4. Confocal images are partial Z-stack maximum intensity projections of the posterior brain (left), ellipsoid body (middle), and cell bodies (right). Scale bars are 40μm. Note that there may be trafficking issues or toxicity of HcKCR1 as we see what is likely lysosomal buildup in the cell bodies. b) Representative current-clamp trace from an HcKCR1-expressing EL neuron in response to 10 pulses of 100 ms LED stimulation (620 nm; orange bar) delivered at 5-second intervals. Individual responses from three LED pulses are shown in gray; the black trace is the trial-averaged voltage response from all 10 pulses. c) Summary of the peak light-evoked hyperpolarization (*Δ*voltage) in EL neurons expressing HcKCR1 (green) versus GFP-only controls (gray). Each dot represents the mean response from a single cell, and lines indicate group averages. HcKCR1 positive neurons show significantly greater hyperpolarization (Mann–Whitney U test HcKCR1 (n=5) vs. control (n=6): U = 0.0, p=0.0015). d) EPG response to optogenetic stimulation during pairing protocol. The across-fly average z-scored fluorescence in the 200 milliseconds following the optogenetic stimulation is shown for every ellipsoid body wedge (rows) at each stimulation location (columns). The blue cue in the schematic below the x-axis shows the location of the visual cue during optogenetic stimulation. The 0 radian (left-red colormap) and *π* radian (middle-blue colormap) pairings are shown separately as well as the combined responses regardless of the location of the visual cue (right-greyscale colormap). *Top:* EL-Gal4 only control. *Bottom:* EL-Gal4>HcKCR1 e) Average EPG response to optogenetic stimulation combined across stimulation locations. Each column in [d] is shifted to align the locations of stimulation. Data is plotted as across fly mean ± sem. f) Average offset vector lengths for all baseline and post pairing sessions. Each dot represents the vector length from one session from a single fly. Lines indicate across fly means. g) EPG bump-cue offsets for baseline and post pairing sessions. Each dot represents data from a single session. Lines connect data from the same fly.

**Extended Data Figure 14.**
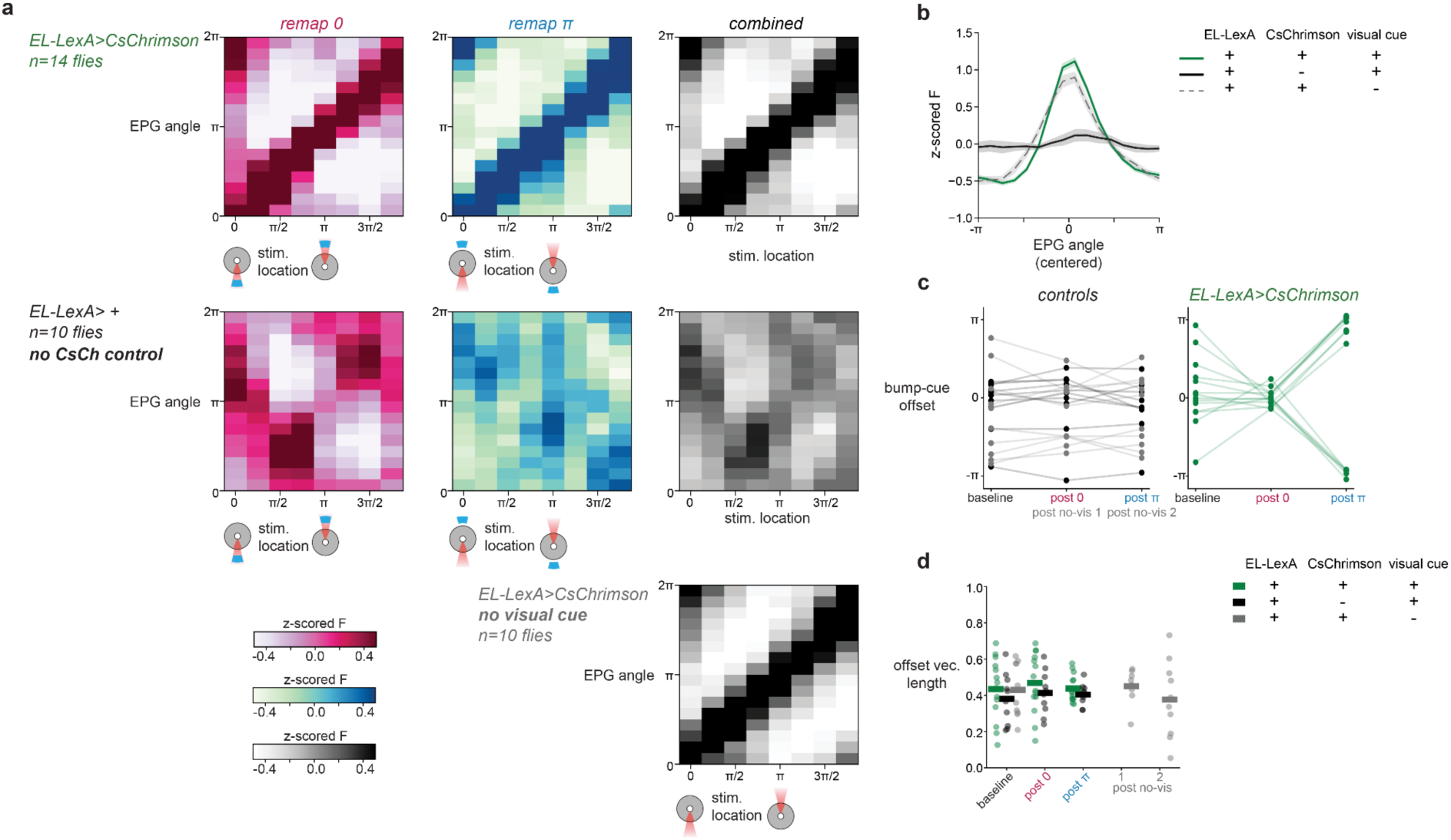
Quantification of the impact of EL activation. Panels in this figure pertain to data in Fig. 6. A,b) Localized optogenetic activation of EL moves the EPG bump (Same as Extended Data Fig. 13d,e for conditions in Fig. 6a-d). Without CsChrimson (EL-LexA> +) there is no bump movement (middle). For the “No visual” control condition with EL-LexA>CsChrimson, only the combined optogenetic response is shown (bottom). C,d) Pairing localized optogenetic activation of EL with visual input changes the bump-cue offset. Same as Extended Data Fig. 13f,g for conditions in Fig. 6a-d. “No CsChrimson” control data is shown in black. “No visual” control data is shown in grey (post no-vis 1 and 2 – closed loop VR imaging sessions that follow optogenetic plasticity induction in which no visual cue was present). EL-CsChrimson is shown in green.

**Extended Data Figure 15.**
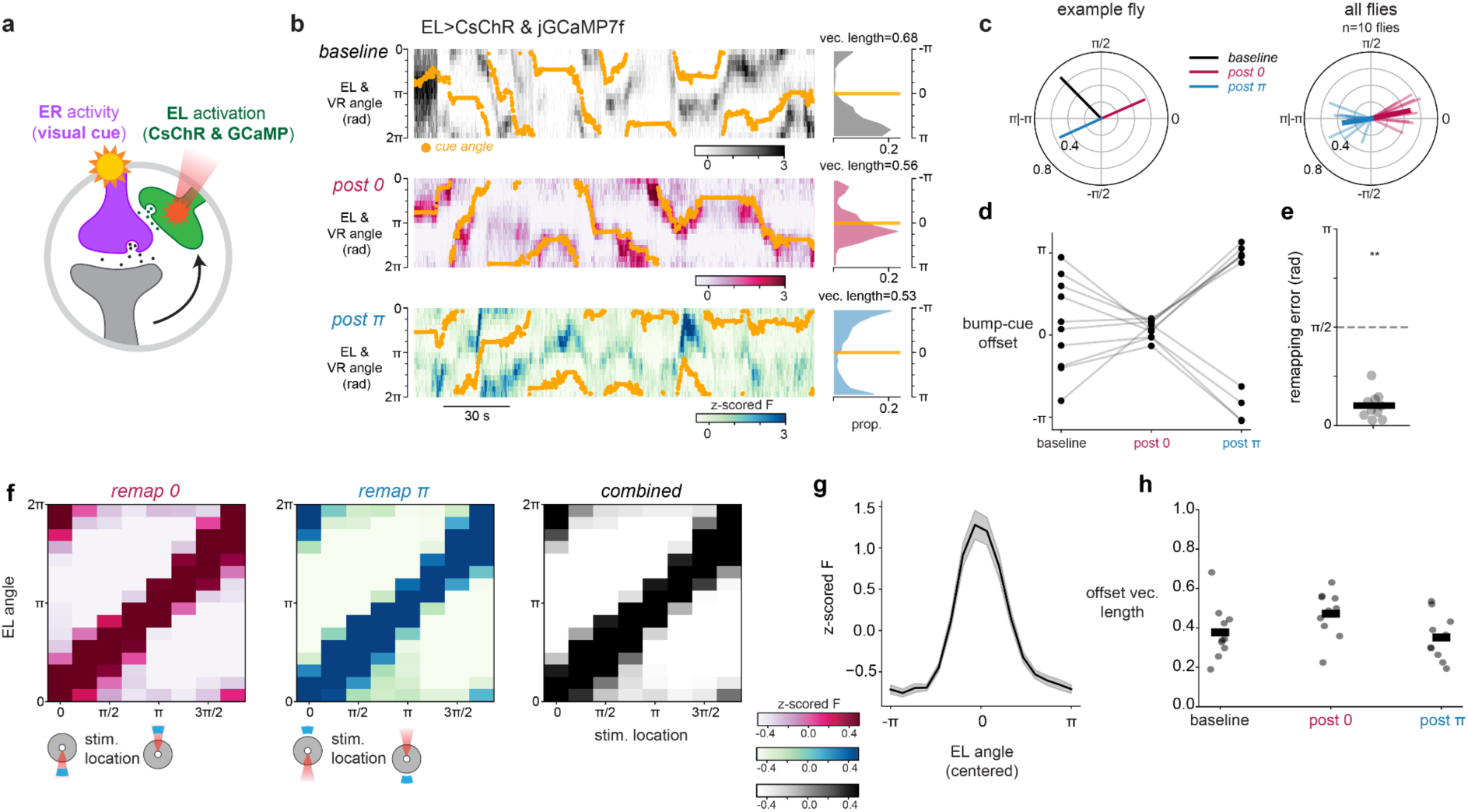
Pairing localized EL activation with visual experience drives plasticity of EL bump-cue offsets. a) Illustration of optogenetic plasticity strategy as in Fig. 6a, but here EL neurons are simultaneously imaged with GCaMP and stimulated using CsChrimson (EL-split Gal4>jGCaMP7f & CsChrimson). b) Example imaging data showing successful EL-induced EL bump-cue plasticity. Data shown as in Fig. 5 & 6. c) Bump-cue offset average vectors for example data in [b] (*left*) and all flies from this genotype (*right*). d) EL bump-cue offsets for baseline and post pairing sessions. Each dot represents data from a single session. Lines connect data from the same fly. e) Remapping errors as shown previously. Wilcoxon signed-rank test for difference from *π*/2: W=0 p=0.002. f) EL response to optogenetic stimulation during pairing protocol as in Extended Data Fig. 13d. g) Average EL response to optogenetic stimulation combined across stimulation locations as in Extended Data Fig. 13e. h) Average offset vector lengths for all baseline and post pairing sessions. Each dot represents the vector length from one session from a single fly. Lines indicate across fly means.

**Extended Data Figure 16.**
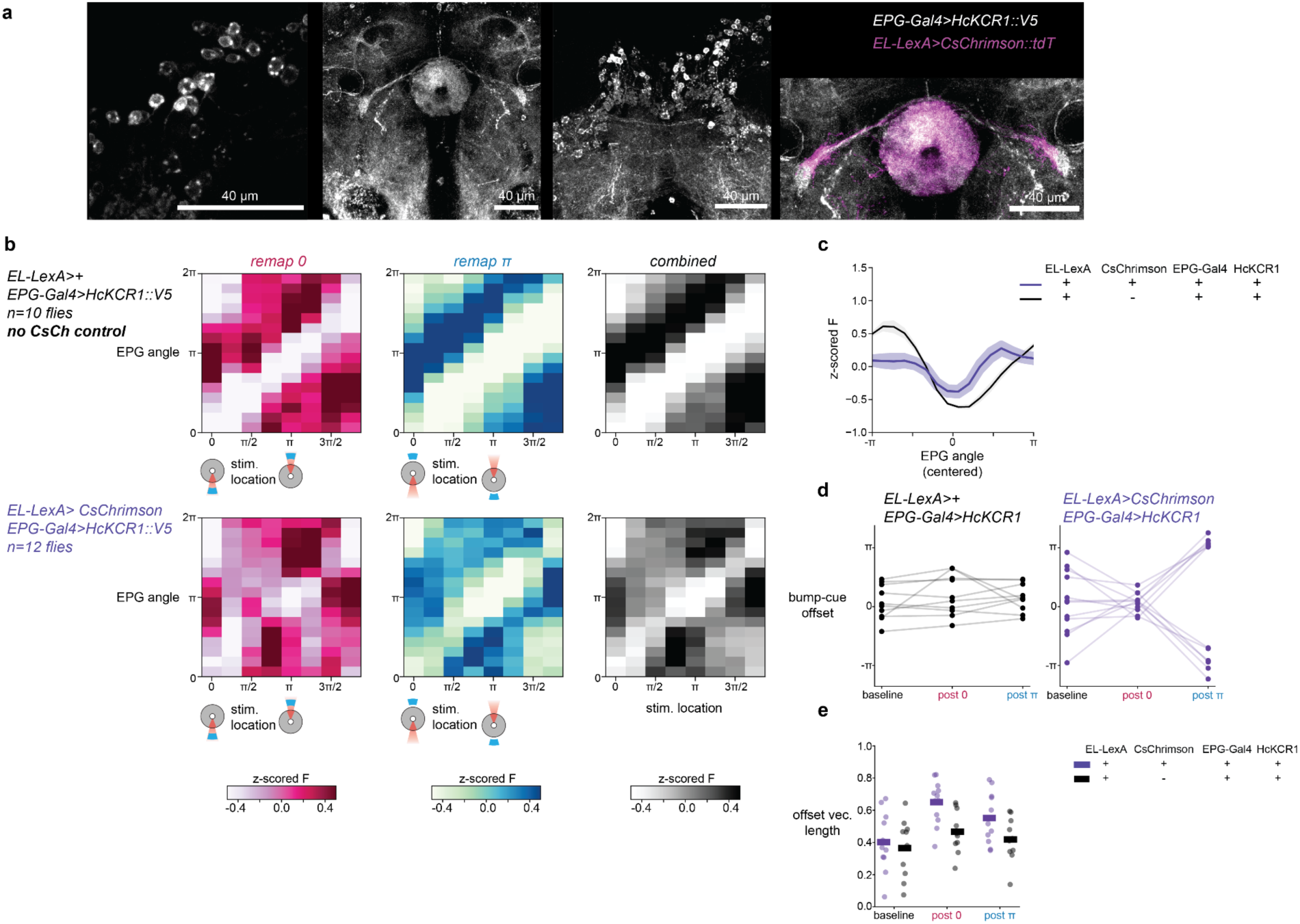
Validation of optogenetic silencing of EPG neurons with HcKCR1. Panels in this figure pertain to data in Fig. 6f-j. a) Adult brain with HcKCR1::V5 expression (anti-V5, gray) driven by the R60D05-Gal4 driver line. Confocal images are maximum intensity projections of cell bodies (left), ellipsoid body (middle left), and protocerebral bridge (middle right). *Right:* brain in which EL neurons are also labeled by CsChrimson::tdTomato (anti-dsRed, magenta).Scale bars are 40μm. b-e) EPG silencing alone (top-black) suppresses the EPG bump but does not induce changes in bump-cue offset. EPG silencing during EL optogenetic pairing (bottom-purple) suppresses the EPG bump and displays successful EPG bump-cue plasticity. b) The heatmap shows the across-fly average z-scored fluorescence in the 200 milliseconds following the optogenetic stimulation is shown for every ellipsoid body wedge (rows) at each stimulation location (columns) (Plotted as in Extended Data Fig. 13d). c) Average EPG response to EPG silencing (black) or EPG silencing during EL activation (purple) combined across stimulation locations. Each column in [b] is shifted to align the locations of stimulation. Data is plotted as across fly mean +/−sem. d) EPG bump-cue offsets for baseline and post pairing sessions. Each dot represents data from a single session. Lines connect data from the same fly. e) Average offset vector lengths for all baseline and post pairing sessions. Each dot represents the vector length from one session from a single fly. Lines indicate across fly means.

**Supplementary Video 1:** Dynamic visualization of 2P imaging data and VR behavior from imaging session shown in Fig 1g.

**Supplementary Video 2:** 2P data and VR behavior from optogenetic pairing experiment shown in Fig 6b

## Methods

### Flies

Unless otherwise specified, *Drosophila melanogaster* were raised on molasses-based food in an incubator (IN034, Darwin Chambers) on a 12 h:12 h light:dark cycle at 25 °C and ∼70% relative humidity. Flies expressing HcKCR1^85^ or CsChrimson^138^ were raised throughout development on Nutri-Fly “German Food” Sick Fly Formulation no. 66-115 (Genesee), containing ∼0.6 mM all-trans-retinal (Sigma) and the anti-fungal agent (Tegosept). Fly vials containing all-trans retinal were wrapped in foil to prevent its photoconversion.

Experimenters were not blinded to fly genotype. Data from experimental and control genotypes were collected concurrently to avoid batch effects. Sample sizes were chosen based on conventions in the field, expected fly-to-fly variability from published results, and pilot data. For all *in vivo* experiments flies had at least one wild-type copy of the white (w) gene. Fly genotypes used in each figure are as follows:

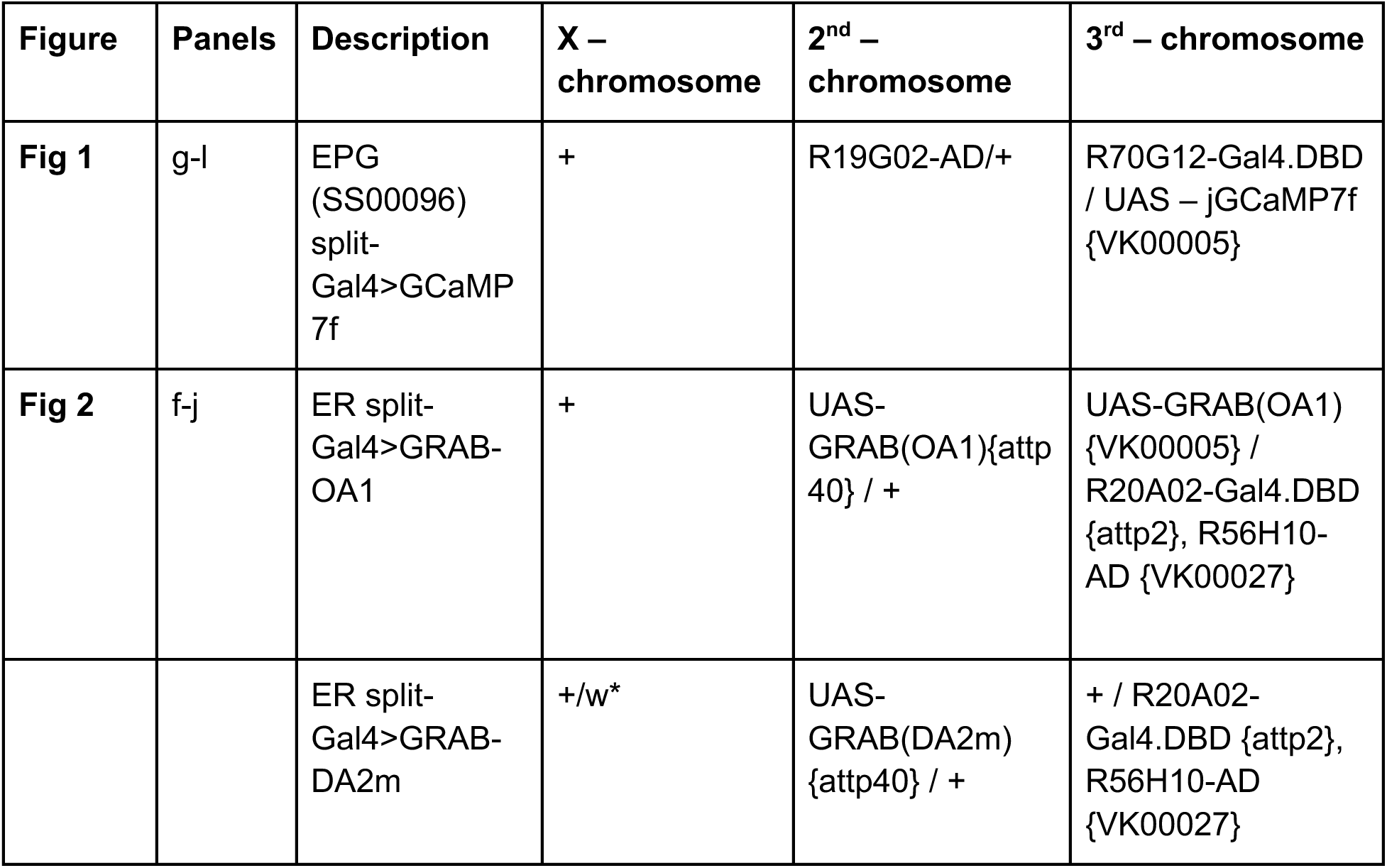
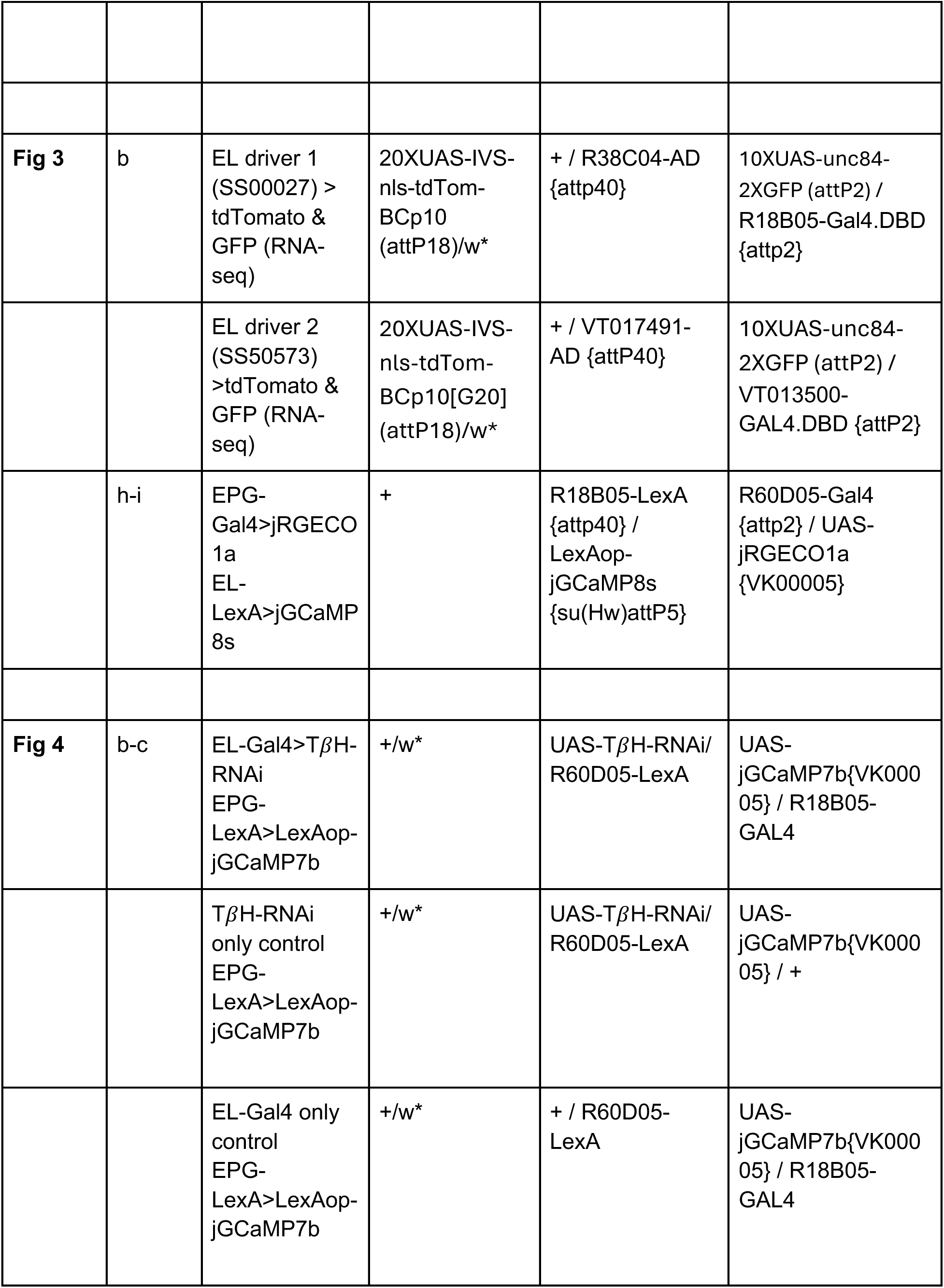
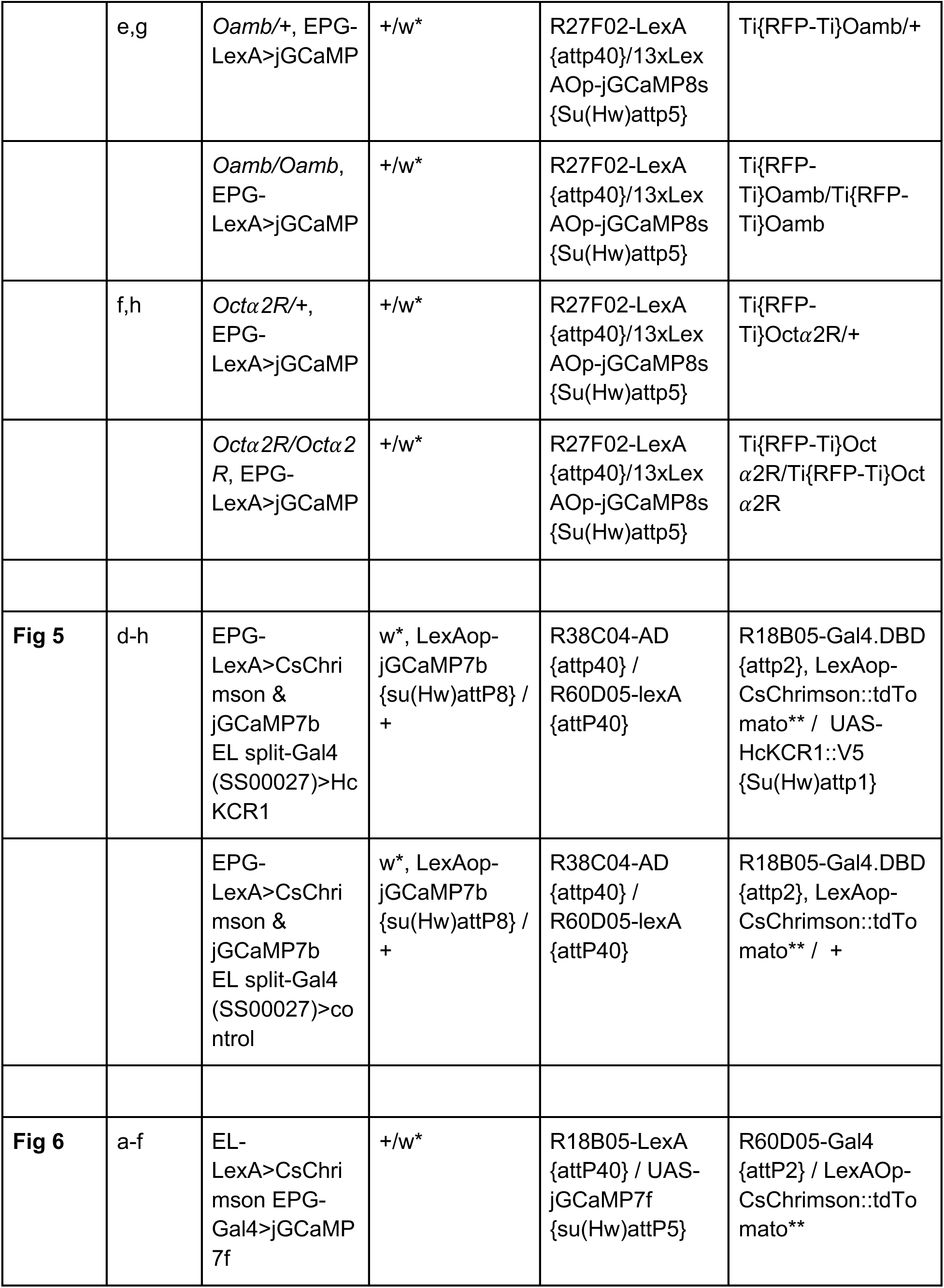
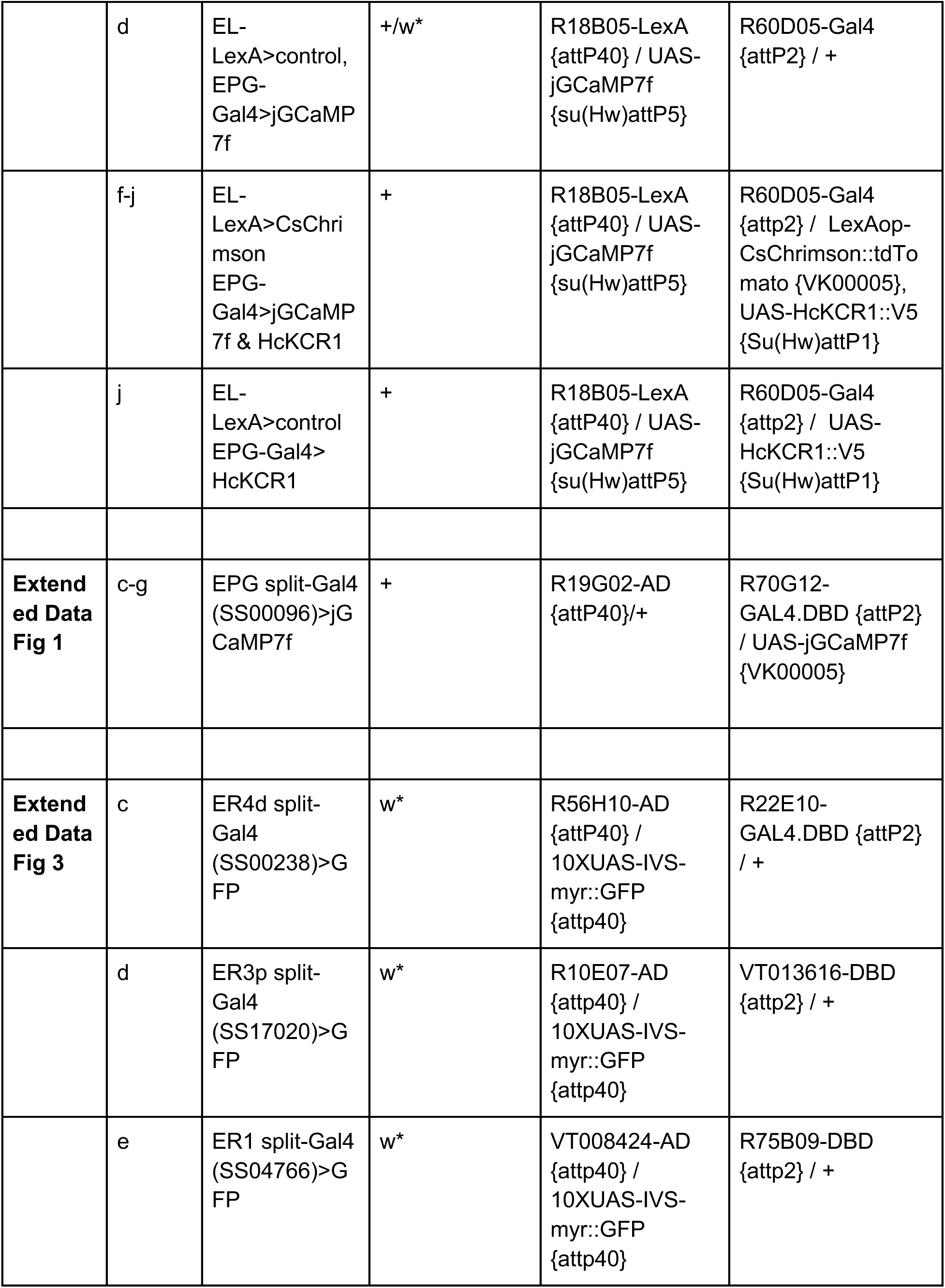
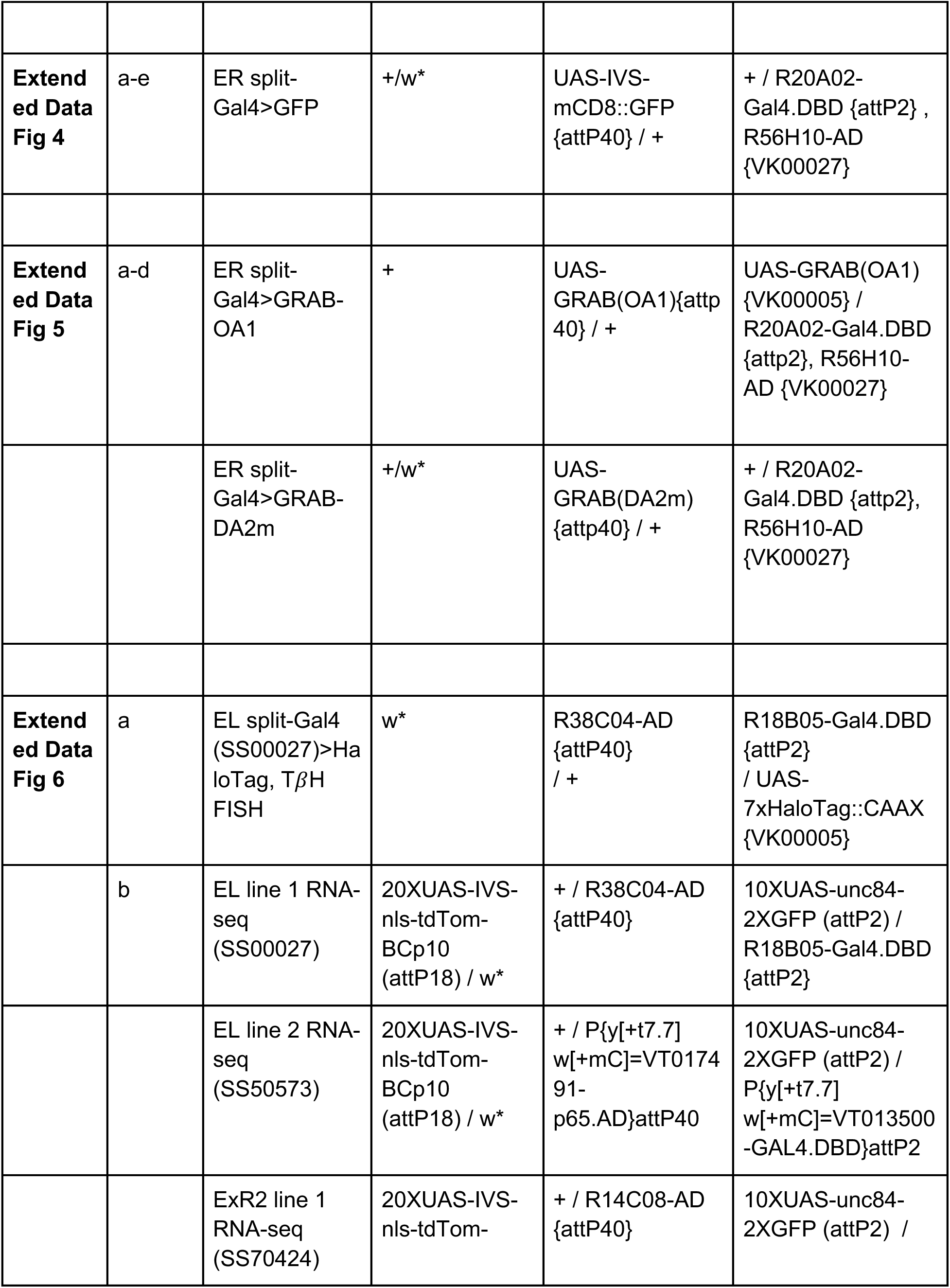
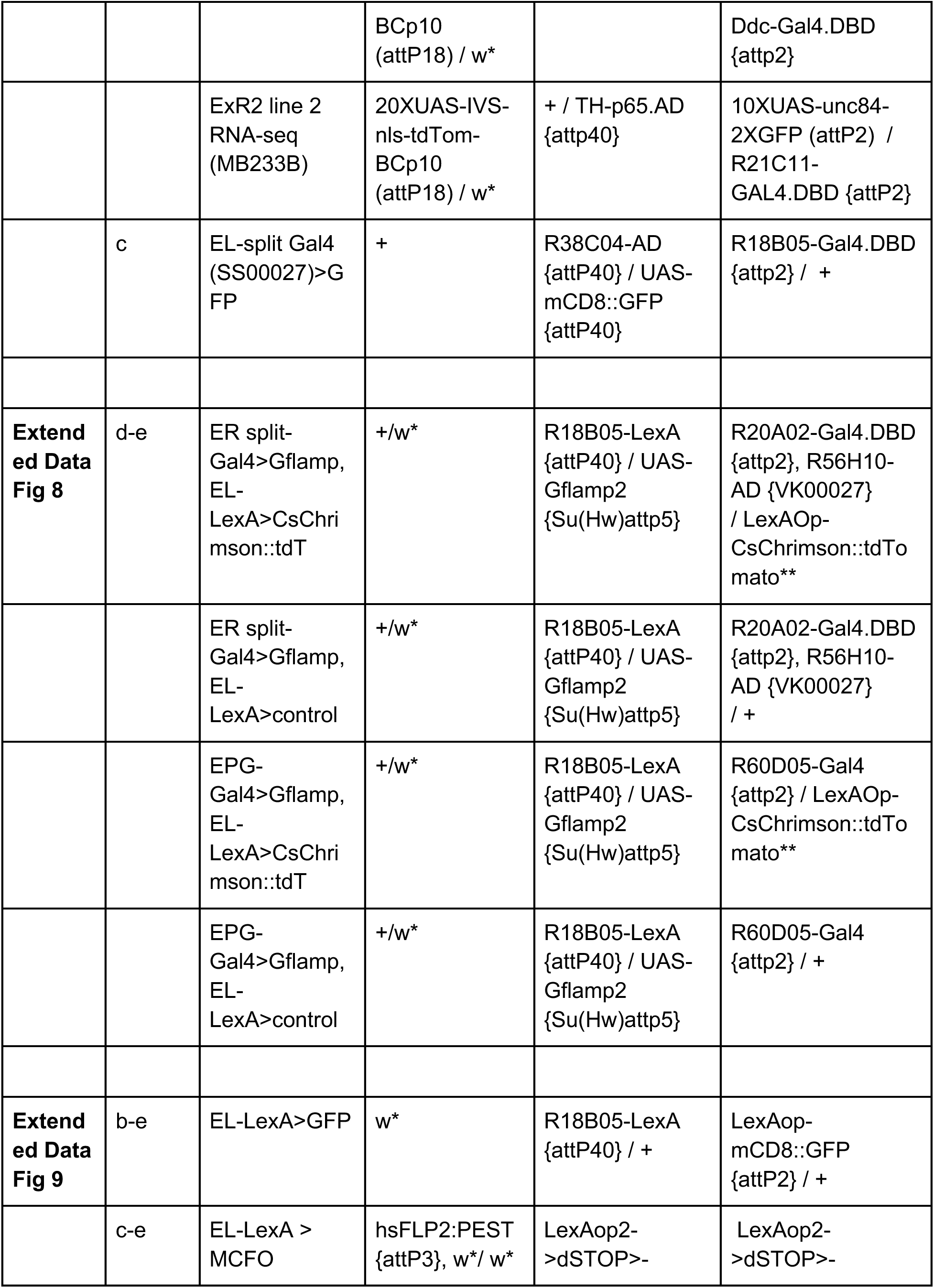
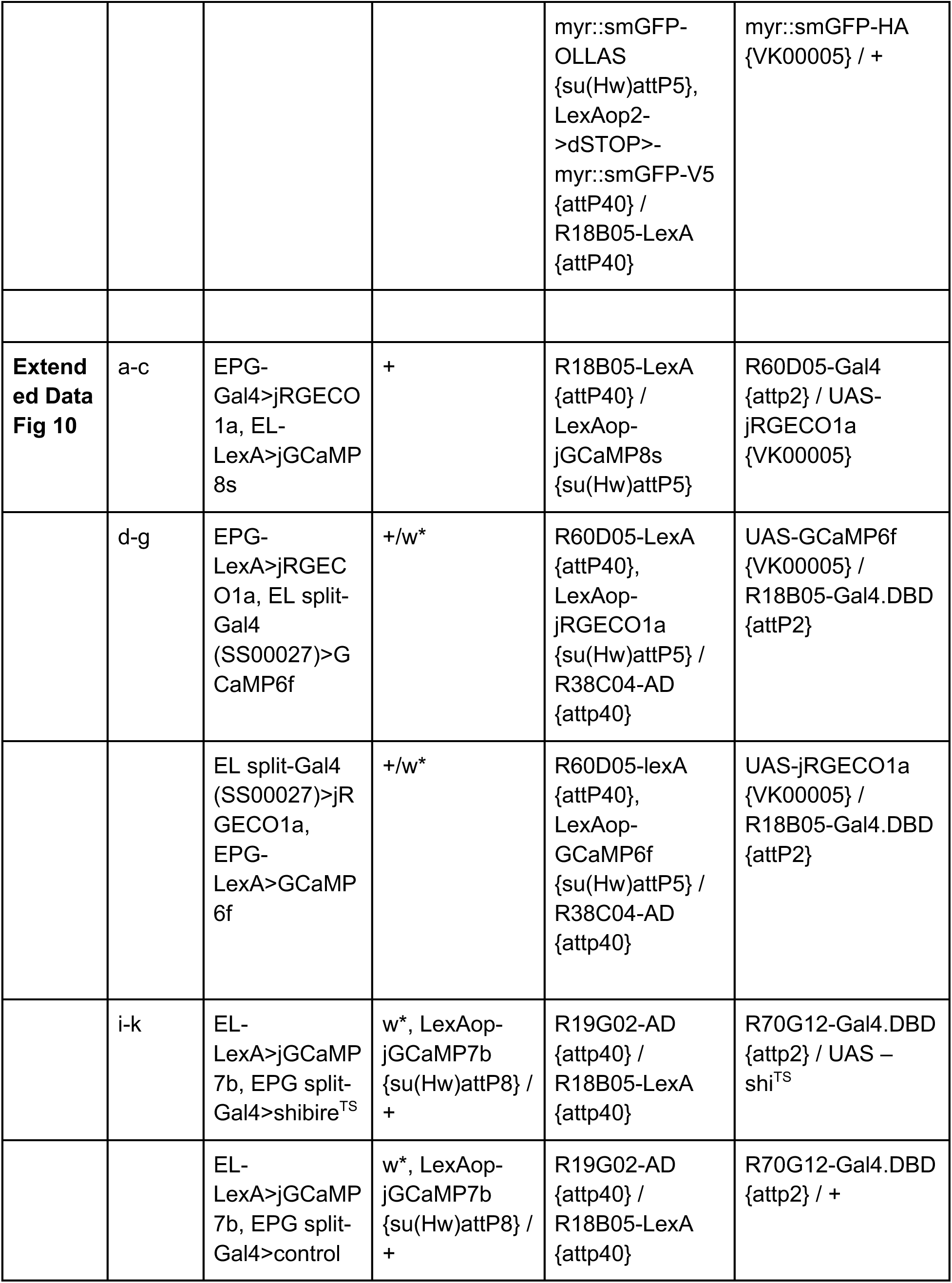
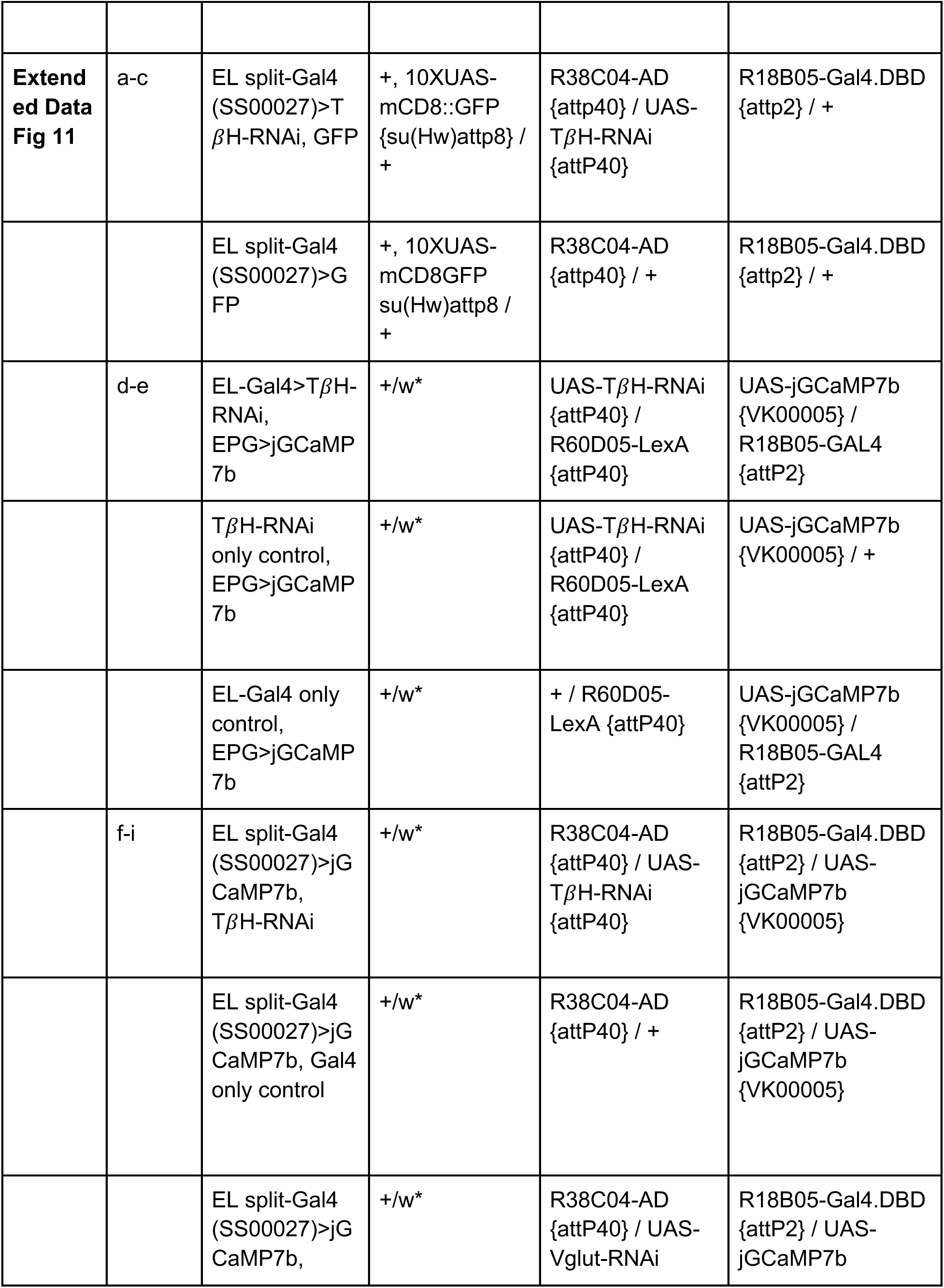
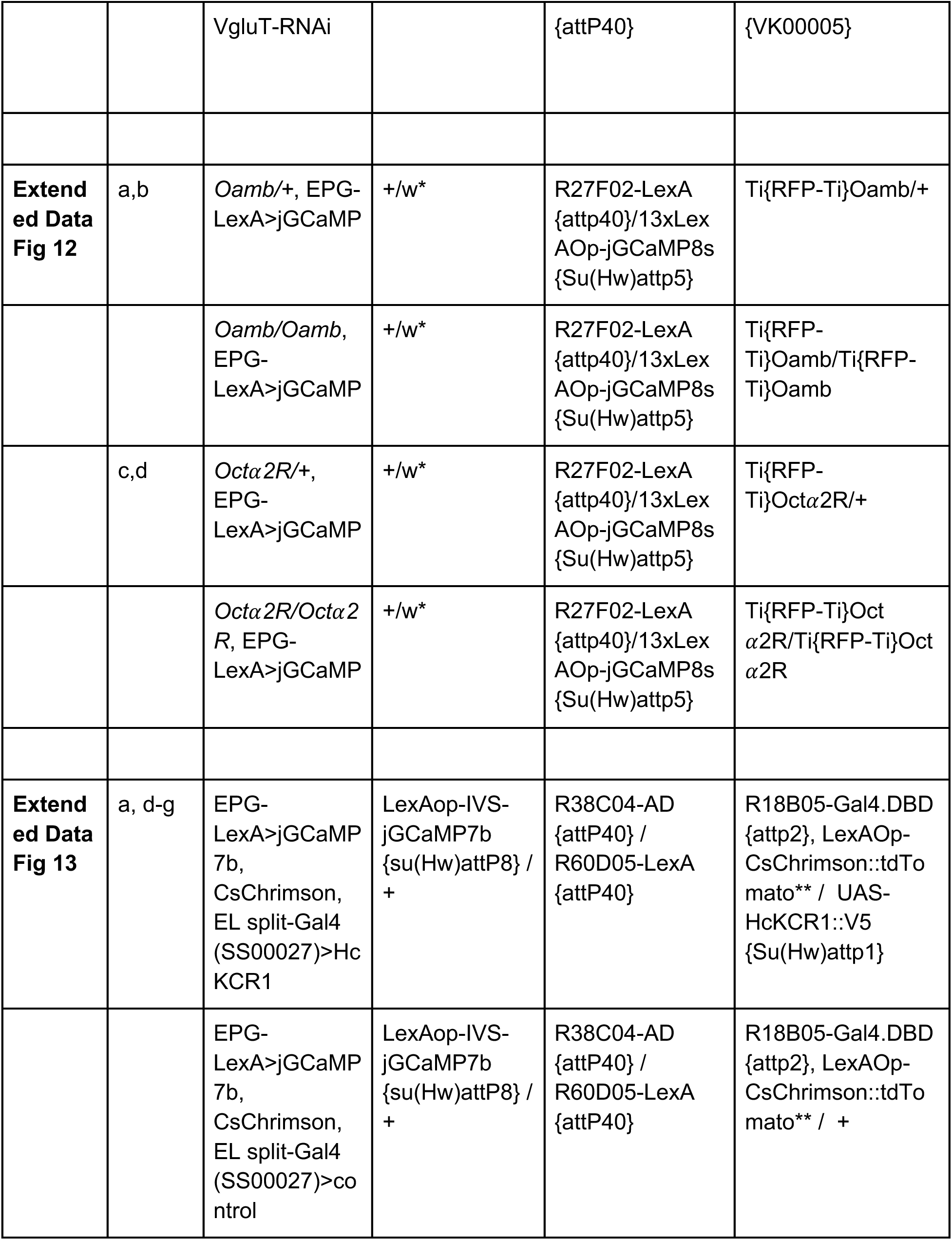
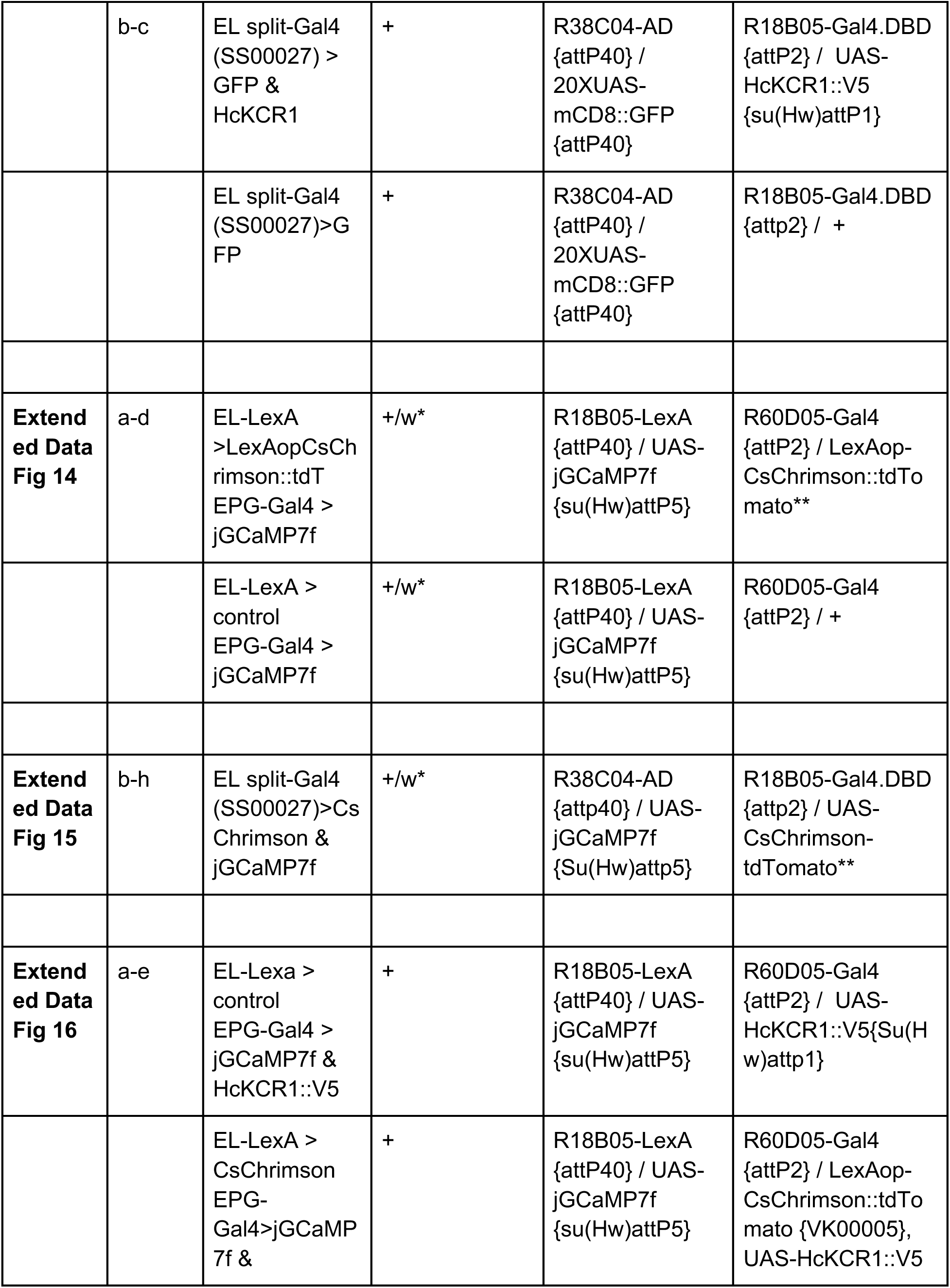
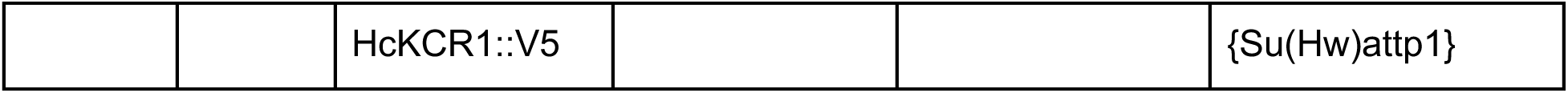

### Origins of Transgenic stocks

| Transgenic line<br>(in figure order of appearance) | Full transgene insertion (if known) | Origin | BDSC number (if applicable) |
|---|---|---|---|
| EPG (fly light number SS00096) split-Gal4 | P{y[+t7.7] w[+mC]=R19G02-p65.AD}attP40; P{y[+t7.7] w[+mC]=R70G12-GAL4.DBD}attP2 | Bloomington Drosophila Stock Center (BDSC) | 86861 |
| UAS-jGCaMP7f {VK00005} | Pbac{y[+mDint2] w[+mC]=20XUAS-IVS-jGCaMP7f}VK00005 | BDSC | 79031 |
| R20A02-DBD {attp2} | P{y[+t7.7] w[+mC]=R20A02-GAL4.DBD}attP2 | BDSC | 600788 |
| R56H10-AD {VK00027} | Pbac{y[+mDint2] w[+mC]=R56H10-p65.AD}VK00027 | BDSC | 601628 |
| UAS-GRAB(DA2m) {attp40} | P{y[+t7.7] w[+mC]=10XUAS-GRAB(DA2m)}attP40 | BDSC | 90878 |
| UAS-GRAB(OA1) {attp40} | P{y[+t7.7] w[+mC]=10XUAS-GRAB(OA1)}attP40 | Gift from Y. Li. Now deposited at BDSC | 604600 |
| UAS-GRAB(OA1.0) {VK00005} | Pbac{y[+mDint2] w[+mC]=10XUAS-GRAB(OA1.0)}VK00005 | Gift from Y. Li. Now deposited at BDSC | 604601 |
| EL driver 1 (fly light number SS00027) split-Gal4 | Pbac{y[+mDint2] w[+mC]=R38C04-p65.AD}attp40 ; Pbac{y[+mDint2] w[+mC]=R18B05-p65.DBD}attP2 | Gift from G. Rubin prior to publication<sup>26</sup> |  |
| EL driver 2 (fly light number SS50573) | P{y[+t7.7] w[+mC]=VT017491-p65.AD}attP40; P{y[+t7.7] w[+mC]=VT013500-GAL4.DBD}attP2 | Gift from G. Rubin prior to publication<sup>26</sup> |  |
| 20XUAS-IVS-nls-tdTom-BCp10 (attP18) |  | Janelia Research Campus. Previously published<sup>27</sup> |  |
| 10XUAS-unc84-2XGFP (attP2) |  | Janelia Research Campus. Previously published<sup>27</sup> |  |
| EPG-Gal4 (R60D05-Gal4) | P{y[+t7.7] w[+mC]=GMR60D05-Gal4}attP2 | BDSC | 39247 |
| EL-LexA (R18B05-lexA) | w[1118]; P{y[+t7.7] w[+mC]=GMR18B05-lexA}attP40 | BDSC | 52525 |
| UAS-jRGECO1a | Pbac{y[+mDint2] w[+mC]=20XUAS-IVS-NES-jRGECO1a-p10}VK00005 | BDSC | 63794 |
| LexAop-jGCaMP8s | P{y[+t7.7] w[+mC]=13XlexAop-IVS-RSET-jGCaMP8s}su(Hw)attP5 | BDSC | 605078 |
| EL-Gal4 (R18B05-Gal4) | P{y[+t7.7] w[+mC]=GMR18B05-GAL4}attP2 | BDSC | 48799 |
| UAS-TβH-RNAi | P{y[+t7.7] v[+t1.8]=TriP.HMS05829}attP40 | BDSC | 67968 |
| LexAop-jGCaMP7b {VK00005} | Pbac{y[+mDint2] w[+mC]=13XlexAop-IVS-jGCaMP7b}VK00005 | BDSC | 80915 |
| EPG-LexA (R60D05-LexA) | P{y[+t7.7] w[+mC]=GMR60D05-lexA}attP40 | BDSC | 52867 |
| EPG-LexA (R27F02-LexA) | P{y[+t7.7] w[+mC]=GMR27F02-lexA}attP40 | BDSC | 52748 |
| LexOp-jGCaMP8s {Su(Hw)attp5} | P{y[+t7.7] w[+mC]=13XlexAop-IVS-RSET-jGCaMP8s}su(Hw)attP5 | BDSC | 605081 |
| Oamb knockout | TI{RFP[3xP3.cUa]=TI}Oamb[attP] | BDSC | 84555 |
| Octα2R knockout | TI{RFP[3xP3.cUa]=TI}Octalpha2R[attP] | BDSC | 84472 |
| LexAop-CsChrimson::tdTomato** | Unknown 3<sup>rd</sup> chromosome insertion. Recombines readily with known insertions at VK00005, but not with insertions at Su(Hw)attp1. | Gift from J. Tuthill |  |
| LexAop-jGCaMP7b {su(Hw)attp8} | w[1118], P{y[+t7.7] w[+mC]=13XlexAop-IVS-jGCaMP7b}su(Hw)attP8 | BDSC | 80911 |
| UAS-HcKCR1::V5 | Pbac{y[+mDint2] w[+mC]=20XUAS-IVS-HcKCR1::V5}Su(Hw)attP1 | This manuscript |  |
| UAS-jGCaMP7f {Su(Hw)attp5} | P{y[+t7.7] w[+mC]=20XUAS-IVS-jGCaMP7f}su(Hw)attP5 | BDSC | 80906 |
| LexAop-CsChrimson::tdTomato {VK00005} | Pbac{y[+mDint2] w[+mC]=13XlexAop2-IVS-CsChrimson.tdTomato}VK00005 | BDSC | 82183 |
| ER4d split-Gal4 (fly light number SS00238) | P{y[+t7.7] w[+mC]=R56H10-p65.AD}attP40; P{y[+t7.7] w[+mC]=R22E10-GAL4.DBD}attP2 | BDSC | 86863 |
| ER3p split-Gal4 (fly light number SS17020) | P{y[+t7.7] w[+mC]=R10E07-p65.AD}attP40; P{y[+t7.7] w[+mC]=VT013616-GAL4.DBD}attP2 | Janelia Research Campus. Previously published<sup>26</sup> |  |
| ER1 split-Gal4 (fly light number SS04766) | P{y[+t7.7] w[+mC]=VT008424-p65.AD}attP40; P{y[+t7.7] w[+mC]=R75B09-GAL4.DBD}attP2 | Janelia Research Campus. Previously published<sup>26</sup> |  |
| UAS-IVS-myr::GFP | P{y[+t7.7] w[+mC]=10XUAS-IVS-myr::GFP}attP40 | BDSC | 32198 |
| UAS-mCD8::GFP {attP40} |  | Gift from B. Pfeiffer and G. Rubin, previously published in<sup>139</sup> |  |
| UAS-7xHaloTag::CAAX {VK00005} |  | Janelia Research Campus. Previously published in<sup>139</sup> |  |
| ExR2 line 1 (fly light number SS70424) | P{y[+t7.7] w[+mC]=R14C08-p65.AD}attP40; Ddc-Gal4.DBD} attp2 | Janelia Research Campus. Previously published<sup>140</sup> |  |
| ExR2 line 2 (fly light number MB233B) | TH-p65.AD {attp40}; P{y[+t7.7] w[+mC]=R21C11-GAL4.DBD}attP2 | Janelia Research Campus. Previously published<sup>141</sup> |  |
| UAS-Gflamp2 {Su(Hw)attp5} | P{w[+mC]=20XUAS-IVS-Gflamp2}Su(Hw)attp5 | Gift from T.R. Clandinin |  |
| LexAop-mCD8::GFP {attP2} | P{y[+t7.7] w[+mC]=13XlexAop2-mCD8::GFP}attP2 | BDSC | 32203 |
| LexAop – MCFO | P{pBPhsFLP2:PEST}attP3; P{13xLexAop2->dSTOP>-myr::smGFP-OLLAS}su(Hw)attP5, P{13xLexAop2->dSTOP>-myr::smGFP-V5}attP40; P{13xLexAop2->dSTOP>-myr::smGFP-HA}VK00005 | Gift from A. Nern and G. Rubin. Previously published<sup>137</sup> |  |
| LexAop-jRGECO1a | P{y[+t7.7] w[+mC]=13XlexAop2-IVS-NES-jRGECO1a-p10}su(Hw)attP5 | BDSC | 64426 |
| UAS-GCaMP6f | Pbac{y[+mDint2] w[+mC]=20XUAS-IVS-GCaMP6f}VK00005 | BDSC | 52869 |
| LexAop-GCaMP6f | P{y[+t7.7] w[+mC]=13XlexAop2-IVS-GCaMP6f-p10}su(Hw)attP5 | BDSC | 44277 |
| UAS-shibire<sup>TS</sup> | P{w[+mC]=UAS-shi[ts1].K}3 | BDSC | 44222 |
| UAS-mCD8::GFP {su(Hw)attP8} | P{y[+t7.7] w[+mC]=10XUAS-IVS-mCD8::GFP}su(Hw)attP8 | BDSC | 32189 |
| UAS-Vglut-RNAi | P{y[+t7.7] v[+t1.8]=TriP.HMS02011}attP2 | BDSC | 40845 |
| UAS-CsChrimson::tdTomato | Unknown 3<sup>rd</sup> chromosome insertion | Gift from J. Tuthill |  |

### Creation of UAS-HcKCR1 transgenic flies

Generation of the UAS-HcKCR1 transgenic fly line was performed by WellGenetics Inc. (New Taipei City, Taiwan). The HcKCR1-V5-6xHis sequence was designed using the HcKCR1 sequence from HcKCR1_mCherry_pcDNA3.1 (Addgene plasmid # 177336)^85^ with the C terminal mCherry sequence replaced by a three alanine linker and V5 and 6xHis sequences followed by a stop codon. This HcKCR1-V5-6xHis 917bp fragment was synthesized with a 4bp Drosophila Kozak sequence added in front of the start codon and cloned into Not1/Xbal sites of the pJFRC7-20XUAS-IVS-mCD8::GFP plasmid replacing the mCD8::GFP coding region using sequence and ligation-independent cloning methods. pJFRC7-20XUAS-IVS-mCD8::GFP was a gift from G. Rubin (Addgene plasmid # 26220)^139^. PhiC31 mediated transgenesis was performed using standard methods to incorporate the plasmid into flies containing the Su(Hw)attP1 landing site to create a final genotype of: P{y[+t7.7]=CaryPw[+mC]=pJFRC7 20XUAS IVS HcKCR1 V5 6xHis}Su(Hw)attP1

### Data analysis

Two-photon imaging, behavioral, electrophysiology, sequencing, and 66onnectomics data were analyzed using custom Python and R scripts (repositories will be made available upon acceptance). Portions of the analysis code were drafted with the assistance of a large language model (Claude Code, Anthropic). Statistical tests were performed using the pingouin (https://pingouin-stats.org/build/html/index.html), statsmodels (www.statsmodels.org/stable/index.html), and SciPy (https://scipy.org/) Python libraries. For parametric tests, Q-Q plots were used to assess normality of residuals. Exact p-values are reported in the figure legends. In figure panels, asterisks denote * p<0.05, ** p<0.01 and *** p<0.001, and n.s. denotes p≥0.05. Where multiple comparisons were performed, p-values were corrected using the Holm method, and corrected values are reported throughout.

### Measuring locomotor behavior

In all two-photon experiments excluding Extended Data Fig. 10d-g, the fly stood on an air-cushioned 9-mm ball made of white foam (FR-4615, General Plastics), painted with black shapes. The ball floated in a custom plastic holder (ABS-like Gray, machined by Proto Labs Inc.) with a semi-spherical depression that cradled the ball. Air flowed into the holder at the base and out of the center of the semi-spherical depression to support the foam ball. The ball was illuminated using infrared LEDs. The movement of the ball was tracked at approximately 450 Hz using a high-speed video camera (Teledyne Flir Chameleon 3, CM3-U3-13Y3M-CS).

FicTrac software^142^ converted the image of the ball into an estimate of its orientation along three axes of rotation. Data from each processed frame was streamed via serial port to a microcontroller (Teensy 4.1). An I2C connection to a 12-bit digital-to-analog converter and a custom op-amp circuit converted the yaw orientation of the ball to a 0-10 Volt analog signal used to control the visual stimulus. We achieved low latency processing such that analog voltage updates were faster than camera frame acquisitions.

### Visual arena and visual stimuli

Visual stimuli were presented using a circular arena composed of modular square panels with a refresh rate of approximately 166 Hz^143^. Each square panel consisted of an 8 x 8 array of blue LEDs (460-475 nm peak wavelength). We used an arena composed of 11 x 2 panels that spanned 330° in azimuth. The remaining 30° degree gap was positioned behind the fly (–150° clockwise from directly in front of the fly) to allow for imaging of the ball. The top edge of the visual arena was approximately aligned with the fly’s vertical position, and the visual arena spanned ∼43° vertically of the fly’s visual field. All visual stimuli consisted of a vertical bar that spanned the full height of the arena and was 2-pixels wide (∼7.5° in azimuth). During closed loop trials, the angle of the bar was set to the yaw angle estimate of the ball via the 0-10 V analog input. During dark experiments, all pixels were turned off. See below (“Optogenetic plasticity protocol”) for description of visual stimuli during optogenetic pairing. To minimize detection of the visual stimulus by the PMTs, the front surface of the visual arena was covered with 3 layers of a blue bandpass gel filter (Tokyo Blue-071 LEE Filters) and a single layer of a neutral density dimming filter (0.9ND-211 LEE Filters). To reduce reflections, a layer of Parafilm was stretched over the front of the filters. The remaining surfaces of the arena were covered with blackout fabric (BK5, Thorlabs Inc.).

### Two-photon imaging and optogenetics

#### Fly preparation for in vivo imaging

To promote walking flies were starved for 12-24 hours prior to imaging. Female flies 5-10 days post eclosion were cold-anaestethized and placed in a custom fly holder with an inverted pyramid shape (Acetal Homopolymer Delrin 150, machined by Proto Labs Inc). The fly’s thorax and head were secured in the holder with the head pitched forward such that most of the eye was below the edge of the holder and the posterior surface of the head was roughly flush with the bottom of the holder. The head was further secured using UV curable glue (Loctite 3972) and a brief pulse (<1s) of UV light (LED-200, Lightning Enterprises). The wings were repositioned to be within the pyramid of the holder and out of the way of the legs. The proboscis was then paralyzed by clipping the rostral protractor muscle using sharpened forceps. The dorsal surface of the fly was covered with saline and a hole was cut in the posterior surface of the head capsule to expose the area above the central complex. Some trachea were removed to allow optical access and reduce brain movement. The external solution contained (in mM): 103 NaCl, 3 KCl, 5 N-tris(hydroxymethyl) methyl-2-aminoethane-sulfonic acid, 8 trehalose, 10 glucose, 26 NaHCO_3_, 1 NaH_2_PO_4_, 1.5 CaCl_2_ and 4 MgCl_2_. For dual calcium indicator imaging (Fig 3 and Extended Data Fig 10a-d), EPG silencing during EL imaging (Extended Data Fig. 10i-k), Oamb knockout (and heterozygotes) with EPG imaging (Fig. 4e,g Extended Data Fig. 12 a,b), and two-photon optogenetic experiments (Fig 5 & 6, Extended Data Fig 13-16) calcium concentration was increased to 2 mM to increase signal to noise ratio. For all external saline, osmolarity was adjusted to 270–275 mOsm. The external solution was bubbled with 95% O2 and 5% CO2 and reached a final pH of 7.1-7.3. After dissection, the fly was immediately placed on the floating ball under the two-photon microscope. The microscope objective was positioned over the region of interest and warmed bubbled saline (25-27°C) was continuously perfused over the brain by a peristaltic pump running at 20-40 revolutions per minute (120 Series Pump (120U/DM2), Watson-Marlow). The fly walked in closed loop with a vertical bar for at least 10 minutes prior to imaging for most experiments. For T*β*H RNAi and Oct*α*2R knockout experiments flies walked in darkness for 10 minutes followed by 10 min of closed loop experience prior to imaging. For Oamb knockout experiments, flies walked in closed loop for 15 minutes prior to imaging. For EPG silencing using shi^TS^ flies walked in darkness for the full duration of the experiment.

#### Ex vivo imaging brain preparation

For *ex vivo* imaging, female flies aged 5-15 days were used for all experiments. Flies were cold anaesthetized and their cuticle was briefly dewaxed with 70% ethanol. The central brain was then rapidly dissected in chilled oxygenated external saline, and the brain was placed on a poly-l-lysine coated coverslip with the posterior surface facing down. The brain was gradually returned to room temperature by perfusing warmed bubbled saline (25-27°C) for 10 minutes prior to two photon imaging.

#### Two-photon imaging

All two-photon imaging and optogenetic experiments with the exception of data included in Extended Data Figure 10d-h were performed on a Bruker Ultima 2P microscope using two fixed-wavelength femtosecond pulsed lasers (Coherent Axon 920 nm and Coherent Axon 1064 nm, a subset of data was collected using a Spark Alcor 920 nm) and piezoelectric objective scanner. Images were collected using a 25x objective (HC FLUOTAR L 25x/0.95 W VISIR, Leica inc.). Emitted fluorescence was first split using a 565 longpass filter (Chroma). Rejected light was filtered with a 525/80 bandpass filter (Chroma) and passed light was filtered with a 610/75 bandpass filter before being collected using GaAsP photomultiplier tubes (Model H10770PB-40, Hamamatsu).

The imaging region was centered on the ellipsoid body and set for each fly to achieve a high imaging rate (resonant-galvo scanning) while ensuring enough boundary for motion correction. A typical imaging field of view was 300 x 250 pixels collected at 4.83x zoom. All data was collected using volumetric z-scanning with 6 z-slices spaced 5 *μ*m apart, resulting in a volumetric scan rate of 7.6-11.7 Hz. Imaging data was synchronized with behavioral data from FicTrac and the visual arena using the Bruker voltage recording module. For all figures a 920 nm laser was used to image green fluorescent proteins. For dual color imaging (Fig. 3 and Extended Data Fig. 10a-c) simultaneous stimulation with the 920 nm and 1064 nm lasers was used. For 2P optogenetic experiments, the 1064 nm laser was controlled with an independent galvo-galvo scanning module. Laser power for each laser line was typically between 5 mW and 15 mW as measured at the sample plane. For *ex vivo* single photon optogenetic experiments, a 617 nm LED (ThorLabs M617L3) was flashed for 50 ms at an intensity of 1.8 *μ*W as measured at the sample plane and a frequency of 0.1 Hz. Each experiment consisted of 20 LED flashes.

For *in vivo* imaging experiments that did not involve optogenetics, imaging was performed in repeated 6-minute sessions. Closed loop sessions in which a vertical blue stripe was visible and rotating with the ball were interleaved with dark sessions in which the visual arena was not illuminated. Between imaging sessions, the imaging field of view was recentered and the tissue was briefly assessed for photobleaching and damage. The fly walked in closed loop between imaging sessions.

Data in Extended Data Figure 10d-h were collected and preprocessed using previously described methods^27^.

#### EPG silencing with temperature sensitive shibire

Saline perfusion temperature was controlled using an inline heater (Warner Instruments). Flies walked for 10 minutes on the air-supported ball under the microscope after dissection with a perfusion temperature of 25°C perfusion at 20 rpm. After 2P imaging at this permissive temperature (6 min). Saline was heated to 33°C and perfusion was increased 40 rpm. Flies continued walking in darkness for 15 min once the bath temperature stabilized before a second 2P imaging trial was performed.

#### Optogenetic plasticity protocol

For optogenetic experiments, 920 nm imaging power was <14 mW as measured during laser scanning at the sample plane. Flies were given at least 10 minutes of closed loop experience prior to baseline imaging sessions. Each fly first underwent a 3 minute baseline imaging session in which they walked in closed loop VR. Next, each fly underwent two rounds of opto-pairing and post opto-pairing imaging sessions: 0 radian bump-cue offset opto-pairing and *π* radian bump-cue offset opto-pairing in a pseudorandom order.

Opto-pairing session: Each opto-pairing session began with a brief closed loop period. The visual arena then went blank for 5 seconds. The blue bar was then flashed at 8 different orientations around the fly for 2 seconds each in clockwise order. This was repeated 5 times. At the onset of each bar flash, a corresponding wedge of the ellipsoid body was stimulated with the 1064 nm laser at the desired bump-cue offset for 1800 msec (10 *μ*m circle, spiral scanned 100 times). For experiments included in Fig 5 and Extended Data Fig 13, 1064 nm laser power was approximately 20 mW, as measured at the sample plane during mock optogenetic stimulation. For all other optogenetic experiments, 1064 nm laser power was approximately 10 mW. A shutter protected PMTs during optogenetic stimulation but reopened for the last 200 msec of each bar flash, allowing us to measure the calcium response to optogenetic stimulation (analysis details below). For experiments except data included in Fig. 6e, immediately after the last blue bar flash, the visual arena went blank for 5 seconds and then the blue bar reappeared in closed loop. We continued imaging in closed loop for a brief period before ending the opto-pairing session. Opto-pairing sessions lasted less than 3 minutes. After recentering the field of view, we began the post opto-pairing session. For experiments in Fig. 6e, the first optogenetic pairing session was performed as described above. For the second optogenetic pairing, immediately following the last blue bar flash, the visual arena went blank for at least 6 minutes. The bar then reappeared in closed loop as above. For all experiments, the bar reappeared in closed loop at the orientation determined by the fly’s integrated motion on the ball over the dark period.

Post opto-pairing imaging sessions were a 3-minute-long closed loop session.

### Data processing for two-photon imaging

#### Preprocessing for in vivo imaging

Images from two-photon microscopy were motion corrected in each z plane independently. For each plane, a reference image was computed using one tenth of the frames. To compute the reference image: first, a temporal average of the reference frames was computed; second, each frame was aligned to the average using the peak of the spatial cross correlation; third, the temporal average was recomputed. The remaining frames from the imaging session were then aligned to the reference image using the maximum of the spatial cross correlation. Using Napari (https://napari.org/stable/)^144^, for each z plane, one region of interest (ROI) was drawn around the perimeter of the ellipsoid body and a second ROI was drawn around the inner circle of the ellipsoid body. Pixels were considered in the ellipsoid body if they fell within the outer ROI but not inner ROI. The center of mass of the inner ROI was considered the center of the ellipsoid body for each z plane, and the phase of each pixel in the ellipsoid body was calculated relative to this origin (*arctan*(*y*/*x*)). The phases were divided into 16 bins to create wedge-shaped ROIs around the ellipsoid body. Bins with the same phase were combined for all z-planes. Average fluorescence values for each volume were extracted from each ROI. To simultaneously remove artifacts from the visual arena and to correct for photobleaching and slow z drift, time series were fit with a linear model that included a single exponential and the timeseries from a background ROI which did not include fluorescent pixels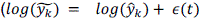 with 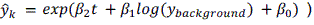 where 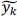 is the raw fluorescence time series of the *kt*ℎ ROI, *y_back_*_g*round*_ is the background signal intensity, *t* is time, *β_i_* are coefficients fit by least squares regression, and *ε*(*t*) is the error of the fit at each time point. The residual time series with nuisance terms removed, 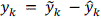, was then z-scored, temporally smoothed by a gaussian kernel of width 100 msec (∼1 imaging volume), and spatially smoothed with a gaussian kernel of width ½ of an ROI (*π*/16 radians of the ellipsoid body).

Fictrac phases were shifted so that a 0 radian orientation aligned with the 0 radian angle of the ellipsoid body and smoothed by a gaussian kernel of width 100 msec. Based on analyses in Extended Data Fig 1, we temporally shifted Fictrac phases by –200 msec (approximately 2 imaging volumes) to maximize correlation between behavioral and neural data.

Data in Extended Data Fig 10d-h was preprocessed using previously described methods^27^. ROI timeseries were smoothed and z-scored using identical parameters as above.

#### Population vector analyses

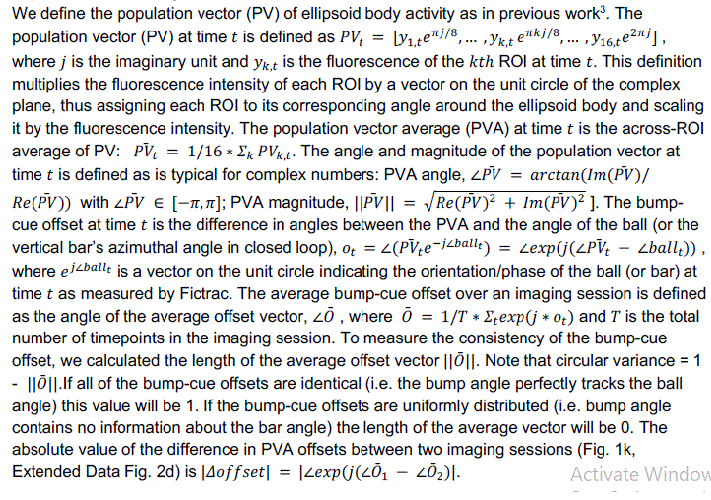

#### Bump width estimation (Fig 1l, 4g,h)

Consider the matrix of ROI fluorescence values over time 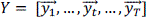 where 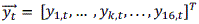 is a column vector. For each *t* we calculated the discrete Fourier transform in the spatial domain yielding the matrix, 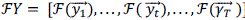. To phase null the data, we multiplied each harmonic, *m*, by *exp*(*jmφ*_1,*t*_), yielding 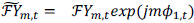, where *φ*_1,*t*_ is the phase of the first harmonic. The first harmonic is equivalent to the population vector with a sign flip in the angles, so −*φ*_1,*t*_ is the PVA angle. This operation circularly shifts the population vector at every timepoint by the PVA angle to sub-ROI precision. The inverse Fourier transform of 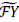 was then rolled to center the bump profile. The phase-nulled profiles were averaged over time, giving one mean bump profile per imaging session.

Bump width was measured from this mean phase-nulled profile as a full width at half maximum (FWHM), with no parametric fit. The 16-point profile was interpolated to 256 points by zero-padding its Fourier spectrum. The interpolated profile was rolled so that its peak lay at the first sample, and the half-maximum level was taken as the average of the profile’s minimum and maximum. Moving outwards from the peak in each direction, the first sample below the half-maximum level was found and the crossing determined by linear interpolation between it and the preceding sample. The FWHM is the distance between the two crossings in radians.

#### ER octopamine/dopamine bump visualization (Fig 2h, Extended Data Fig 5)

To align GRAB imaging data to heading with sub-ROI precision, we performed a similar phase-nulling procedure, but each harmonic was multiplied by *e*^j*m*∠*ball*^*^t^* to shift the data to align to the flies’ heading. We then took the temporal average of the shifted data for the first and second half of the imaging session separately. To perform a cross-validated alignment of the peaks across flies, we calculated the ∠*P*^M^*V* for the first first half’s average profile and used that value to shift the second half’s average in the Fourier domain.

#### Difference in EL and EPG PVA angles (Fig 3, Extended Data Fig 10)

To quantify the average angular distance between the EL and EPG bumps we calculated the temporal average of the absolute angular difference between the EL PVA angle and EPG PVA angle (Fig. 3i, Extended Data Fig. 10f). For Extended Data Fig. 10a, we binned the absolute angular difference by the average PVA magnitude of the EL and EPG populations at each timepoint. Averages were calculated first within flies and then across flies.

To visualize the extent to which the EL PVA can lead the EPG PVA during rapid turns (Extended Data Fig. 10b,g), the columns of *Y_EL_* (the matrix of ROI fluorescence for the EL channel) and *Y_EPG_* (the matrix of ROI fluorescence for the EPG channel) were shifted in the Fourier domain, as described above, by the EPG PVA angle at each timepoint, ∠*P*^M^*V_t_*_,*EPG*_. We separately took the average of timepoints where the fly’s rotational velocity was greater than *π* rad/s or less then –*π* rad/s.

To quantify the extent to which EL PVA angle leads the EPG PVA angle for all turning speeds (Extended Data Fig 10c,h), we calculated the mean PVA angle difference (EL minus EPG) for time points within each rotational velocity bin. Averages were calculated within each fly and then combined data across flies.

#### Response to two-photon optogenetic stimulation (Fig. 6f, Extended Data Fig. 13-16)

Fluorescence was extracted from ellipsoid body ROIs from the period when the PMT-shutter was open, this was the final 200 msec of each bar flash. This fluorescence data was processed the same as all other data with the difference being that no detrending (linear model fit) or smoothing was performed because of the discontinuous nature of the recordings. Instead, background fluorescence was regressed from each ROI and fluorescence values were z-scored for each ROI independently. Each ROI’s time series was averaged across repeated bar flashes/optogenetic stimulations.

#### Remapping error

For each post-pairing session, we calculated the absolute value of the difference between the average PVA angle and the opto-paired PVA angle. We then averaged this value across the two pairing sessions to get remapping error: 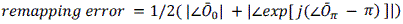, where 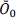 is the average offset vector for the post-0 imaging session and 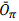 is the average offset vector before the post-*π* imaging session.

#### Ex vivo imaging preprocessing and analysis (Extended Data Fig. 8)

Unlike the in vivo recordings, these time series were not detrended, background-regressed, smoothed or z-scored, because each stimulation is quantified relative to its own preceding baseline. For each stimulation, averaged fluorescence from 2 s before to 8 s after the optogenetic stimulus was linearly interpolated onto a common time base, sampled at 10 Hz, so that trials acquired at slightly different frame times could be averaged together. Imaging frames acquired while the stimulation laser was on were discarded (−0.2 s to +0.22 s relative to the stimulus). The baseline, F_0_, for each stimulation was calculated as the mean fluorescence in the 2 s preceding the stimulus, and the response was expressed as ΔF/F = F/F_0_ − 1. Each stimulation was summarized by the peak of ΔF/F after the excluded window. The 20 stimulations of a session were averaged to give one value per fly.

### Plasticity network model

We translated a previously published model^16^ from MATLAB to Python, implementing both presynaptically-gated and postsynaptically-gated plasticity rules. Following ref. ^16^, “gating” refers to whether weight changes require presynaptic (ER) or postsynaptic (EPG) activity. We modified the learning rate (*ε* in the previously published model) from 0.1 to 10 (0.1, 0.5, 1, 2, 5, 7.5 and 10) to assess robustness of presynaptically and postsynaptically gated learning rules. We simulated a thin vertical bar as a narrow von Mises function with a concentration parameter *κ*=15 as done previously. Uniformly distributed noise was added to rotational velocity at each timepoint *v* = *v_actual_* + 0.4*max*(*v_actual_*) *σ*, where *σ*∼ *U*(−.5,.5) and *v_actual_* is the scaled true rotational velocity of the animal.The visual input drive was set to zero to simulate dark trials. All other parameters were identical to the previous implementation^16^.

Walking trajectories from the flies included in Fig 1g-k were used to simulate network dynamics. The full walking trajectory including data prior to imaging sessions was included in each simulation.

Simulated EPG time series were analyzed similarly to the *in vivo* data except no preprocessing was applied. Each EPG (n=32 model cells) was assigned an orientation around the ellipsoid body at evenly spaced intervals, and we computed PVAs and associated metrics identically as above.

### Cell type naming conventions

All cell types are named as in the hemibrain connectome (v1.2.1)^30^

### Connectome analysis

All connectomics analyses utilized the hemibrain connectome (v1.2.1) and its Python API^30,132^. Summary graphs and cable distance quantifications were created using the NetworkX Python package^145^. For Fig. 3c & Extended Data Fig. 7a-c, only connections between neurons with >3 synapses were considered. For Fig. 2a-e, 3f-g, & Extended Data Fig. 8a-c all connections between neurons were considered. For remaining analyses, synapse number cutoff is stated explicitly in figure.

#### Visualizations in ellipsoid body coordinates

We created a Python implementation of a previously published algorithm for centering visualizations of neuron skeletons on the major axes of a neuropil of interest^31^. Briefly, we performed principal components analysis on the vertices of the ellipsoid body ROI mesh from the hemibrain dataset. We then applied the change of basis from PCA to neuron skeletons and synapse locations. This approach defines the origin as the center of the annulus. The first two principal axes (approximately, medial-lateral and dorsal-ventral) give the plane of the ring, and the third principal axis gives the depth (approximately anterior-posterior). We assigned each voxel an ellipsoid body radius and phase using the coordinates in the first two principal axes.

To obtain peeled convex hulls of each EL neuron’s presynapses (Fig. 3d), we first transformed synapse locations for each neuron into ellipsoid body coordinates and considered only the first two principal axes. To visualize only the most dense region of synapses, we iteratively computed the convex hull of synapse locations and then deleted synapses that formed the vertices of the convex hull. We repeated this procedure 20 times for each cell.

#### Cable distance

To compute cable distances, neuron skeletons of ER and EPG neurons in the hemibrain connectome (v1.2.1) were obtained from Neuprint^132^ and upsampled to maximal resolution, ER-EPG pre– and postsynapses as well as EL-ER and ExR2-ER postsynapses were attached to ER or EPG skeletons, and the skeletons were converted to NetworkX weighted graphs.

Dikstra’s algorithm was used to find the cable distance (through the skeleton) between candidate pairs of synapses. For comparison, comparable analyses were run on *γ*-kenyon cell skeletons analyzing the cable distance between *γ*KC-mushroom body output neuron (MBON) presynapses and dopamine neuron [DAN]-*γ*KC, DANs include *γ*-lobe innervating PPL and PPM neurons) postsynapses.

### Standard IHC protocol & Confocal Imaging

To image adult brains, female flies that were 6–10 days post eclosion were anesthetized with CO_2_ and dissected at room temperature in phosphate buffer solution (PBS).

All immunohistochemistry steps were performed at room temperature in Nunc trays (Cat#:136528, ThermoFisher) with 3-5 brains per well. Brains were transferred between wells using blunt forceps. After dissection, brains were fixed in 24μL 32% aqueous formaldehyde solution in 200μL PBS for exactly 14 minutes. Following fixation, brains were washed in PBS for 10 minutes while moving them between 5 wells. Brains were then incubated for approximately 20 minutes in 5% normal goat serum (NGS) (G9023-10ML, Sigma Aldrich) in PBST (0.2% Triton-X in PBS). After this blocking step, brains were incubated in primary antibody solution for approximately 48 hours while mixing on a Nutator. Then brains were washed in 5 wells filled with PBS with 0.2% Triton X-100 (PBST) for 10 minutes. Next, brains were incubated in a secondary antibody solution for approximately 24 hours while mixing on a Nutator. Finally, brains were then washed in 5 wells filled with PBST for 10 minutes and then transferred to 70% Glycerol in PBS to be stored at 4°C.

Within one week in 70% Glycerol, fly brains were mounted on a slide with Vectashield. Brains were imaged using a Zeiss LSM780 NLO laser scanning confocal microscope (Oberkochen, Germany) housed at the UC Berkeley CRL Molecular Imaging Center, RRID:SCR_017852. The following excitation lasers were used: 488nm, 561nm, 635nm. The samples were imaged using a 40x/1.4 oil-immersion lens. Z-stacks of 120 to 160 optical sections were taken with 1.0-μm spacing. To allow quantitative comparison between genotypes (Extended Data Fig. 11a-c, Tyramine *β*-Hydroxylase RNAi vs control), a fixed gain and laser power was used for all brains. For all other imaging, gain and laser power were optimized for each brain.

### MultiColor Flp Out IHC protocol (R18B05-LexA)

For MultiColor FlpOut (MCFO)^137^ analysis of the R18B05 LexA line, flies were heatshocked for 1-2 hours 8 days after egg laying and dissected at 8 days post eclosion. To administer heatshock, the vial of flies was submerged in a 37°C water bath for 1-2 hours. The same standard IHC dissection, fix, and blocking steps as outlined above was used except with an additional round of staining for the V5 epitope that occurred before brains were transferred to glycerol where brains were incubated in 5% normal mouse serum (NMS) in PBST for 20 minutes then incubated in mouse anti-V5::Alexa 488 (1:500, Invitrogen 37-7500-A488) in 5% NMS in PBST for 24 hours. Finally, brains were washed in 4 wells filled with 10 μL PBST for 10 minutes and placed in 70% glycerol in PBS to be stored at 4°C.

### Primary and Secondary Antibody Solutions

#### R18B05 LexA full line and ER split-Gal4

Primary antibody solution: mouse anti-nc82 (1:20, Developmental Studies Hybridoma Bank, nc82), chicken anti-GFP (1:1000, Abcam, ab13970), and 5% normal goat serum (NGS) (G9023-10ML, Sigma Aldrich) in PBST (0.2% Triton-X in PBS).

Secondary antibody solution: goat anti-mouse Alexa 633 (1:250, Invitrogen, A-21050), goat anti-chicken Alexa 488 (1:250, Invitrogen, A11039), and 5% NGS in PBST.

#### R18B05 LexA MCFO

Primary antibody solution: mouse anti-nc82 (1:20, Developmental Studies Hybridoma Bank, nc82), rabbit anti-HA tag (1:300, Cell Signaling, #3724), rat anti-OLLAS tag (1:200, Invitrogen, MA5-16125), and 5% NGS in PBST.

Secondary antibody solution: goat anti-mouse Alexa 633 (1:250, Invitrogen, A-21050), goat anti-rabbit Alexa Plus 555 (1:500, Invitrogen, A32732), goat anti-rat Alexa Plus 405 (1:250, Invitrogen, A48261) and 5% NGS in PBST.

#### Tyramine Beta Hydroxylase RNAi

Primary antibody solution: mouse anti-nc82 (1:20, Developmental Studies Hybridoma Bank, nc82), chicken anti-GFP (1:1000, Abcam, ab13970), rabbit anti-octopamine (1:100, MyBioSource, MBS2090493), and 5% normal goat serum (NGS) (G9023-10ML, Sigma Aldrich) in PBST (0.2% Triton-X in PBS).

Secondary antibody solution: goat anti-mouse Alexa 633 (1:250, Invitrogen, A-21050), goat anti-chicken Alexa 488 (1:250, Invitrogen, A11039), goat anti-rabbit alexa 568 (1:250, Invitrogen, A11011) and 5% NGS in PBST.

#### HcKCR1::V5 in EL and EPG

Primary antibody solution: mouse anti-V5 (1:500, Abcam, ab27671), chicken anti-GFP (1:1000, Abcam, ab13970), rabbit anti-dsRed (1:250, Takara Bio USA, 632496), and 5% normal goat serum (NGS) (G9023-10ML, Sigma Aldrich) in PBST (0.2% Triton-X in PBS)

Secondary antibody solution: goat anti-mouse Alexa 633 (1:250, Invitrogen, A-21050), goat anti-chicken Alexa 488 (1:250, Invitrogen, A11039), goat anti-rabbit alexa 568 (1:250, Invitrogen, A11011) and 5% NGS in PBST.

### Confocal Image Analysis

For all data maximum intensity Z-projections were created using threshold tools in Fiji/ImageJ. Further analysis was performed on specific genotypes as outlined below:

#### R18B05 LexA full line

For analyzing R18B05 LexA full-line histology, the number of EL neurons per brain was determined by counting cell bodies by eye in Napari^144^.

#### R18B05 LexA MCFO

For R18B05-LexA Multi-Color FlpOut, labeled neurons were identified as EL or non-EL by their morphology. Both the number of distinct labeled cell bodies and distinct wedges in the EB were used to quantify the number of unique EL neurons targeted across the HA, V5, and OLLAS epitope tags. Due to low signal-to-noise, contrast was increased to make the neuronal structure more visible in maximum intensity projections. This introduced significant background noise. FIJI’s freehand selection tool and clearing tool were used to manually remove background noise and improve morphological visualization.

#### Tyramine Beta Hydroxylase RNAi

To compare octopamine levels in TβH RNAi experiments, the fluorescence of Alexa568 (Octopamine) was used. Pixels with a fluorescence value above 100 in the Alexa488 channel (R38C04-AD; R18B05-DBD driving mCD8::GFP) were used to determine which pixels to include in the ROI. Alexa568 fluorescence values within the chosen ROI were averaged to generate the strip plot. An unpaired Mann Whitney U test used to compare genotypes.

### Electrophysiology from EL neurons

#### Electrophysiology fly preparation and dissection

Newly eclosed female flies were collected using cold anaesthesia 3–10 hours before the experiment. Then for the dissection, each fly was cold anaesthetized and placed in a fly holder consisting of flat titanium foil secured within an acrylic platform. The fly’s head was pitched slightly forwards (<30°). The fly was secured in the holder using ultraviolet-curable adhesive (Loctite AA 3972) cured by a brief (<1 s) pulse of ultraviolet light (LED-200, Electro-Lite Co.). To reduce brain movement, muscle 16 and proboscis muscles were clipped with forceps. After the dorsal portion of the head (above the holder surface) was covered in external saline solution, a hole was cut in the head capsule and trachea were removed to expose the posterior surface of the brain. An aperture was made in the perineural sheath by pulling gently with fine forceps or by using suction from a patch pipette containing external saline solution.

The external saline solution was identical to the solution used for two-photon imaging with a 1.5 mM CaCl_2_ concentration.

#### Whole-cell Patch-clamp recordings

Patch pipettes were made from borosilicate glass (Sutter, 1.5 mm o.d., 86 mm i.d.) using a Sutter P-97 puller. Pipettes were fire polished down after pulling using a microforge (ALA Scientific Instruments) to a final resistance of 8–15 MΩ. The internal pipette solution contained (in mM): 140 potassium aspartate, 10 4-(2-hydroxyethyl)1-piperazineethanesulfonic acid, 4 MgATP, 0.4 Na3GTP, 1 ethylene glycol tetraacetic acid, 1 KCl and 13 biocytin hydrazide. The pH was 7.15, and the osmolarity was adjusted to about 267 mOsm. Recordings were performed at room temperature.

To visualize the brain, we used an epifluorescent microscope customized for electrophysiology (Vivoscope, Scientifica) with a Olympus BX51WI fluorescence turret and a 40x water-immersion objective. For illumination far-red light was delivered by a fibre-coupled light-emitting diode (LED; 740 nm, M740F2, Thorlabs) through a ferrule patch cable (200 μm Core, Thorlabs) plugged into a fibre optic cannula (1.25-mm SS ferrule 200-μm core, 0.22 NA, Thorlabs) glued to the recording platform, with the tip of the cannula about 1 cm behind the fly. GFP fluorescence was visualized using a CoolLED light source (pE-300-ultra) with an eGFP-long-pass filter (49018, Chroma). Somatic recordings were obtained in current-clamp mode using a MultiClamp 700B amplifier and a CV-7B headstage (Molecular Devices). Voltage signals were low-pass filtered at 5 kHz before digitization and then acquired with a NiDAQ PCIe-6351 (National Instruments) at 20 kHz. For all electrophysiological recordings, liquid junction potential correction was performed post hoc by subtracting 13 mV from recorded voltages^146^.

#### HcKCR1 stimulation

For optogenetic stimulation during electrophysiology, we used the CoolLED light source (pE-300ultra) to deliver pulses of orange light (590–650 nm, 2 mW, Cy5 long-pass filter cube (49019, Chroma) through the objective. Raw voltage traces from whole-cell current-clamp recordings were collected during a train of 10 light pulses (100 ms duration, delivered every 5 seconds; 0.2 Hz). TTL pulse trains were generated using custom MATLAB scripts and output via a National Instruments PCIe-6351 DAQ board to control illumination timing. A constant depolarizing current was injected before each trial to increase the somatic membrane potential to about –45mV to –55mV.

#### Electrophysiology data analysis

Raw voltage traces from whole-cell current-clamp recordings data were imported into Python and analyzed using custom scripts. Voltage recordings were aligned to stimulation time using a recorded TTL signal from the LED controller. For each trial, the light-evoked voltage response was quantified as the difference between the baseline membrane potential (averaged over a 500 ms window prior to LED onset) and the average voltage during the 100 ms LED stimulation. Responses from individual trials were then averaged to compute the mean response for each cell. Group comparisons were performed using non-parametric statistics (Mann–Whitney U test) to assess whether peak hyperpolarization differed significantly between control and HcKCR1-expressing EL neurons.

### ER octopamine receptor EASI-FISH

Expansion-Assisted Iterative Fluorescence *In Situ* Hybridization (EASI-FISH), DAPI staining, and light sheet imaging were performed as described previously^66,147^. Brains were hybridized with probes designed by Molecular Instruments Inc. to target *Drosophila melanogaster* octopamine receptors (LOT numbers – Oamb: PRH792, Oct-TyR: PRH797, Oct*β*1R: PRH794, Oct*β*2R: PRH795, Oct*α*2R: PRH793 Oct*β*3R: PRH796). Two octopamine receptor types were labeled in each brain. Pair 1:Oamb was detected with Alex Fluor (AF) 546 and Oct-TyR with AF647 hairpins. Pair 2: Oct*β*1R was detected with to AF546 & Oct*β*2R with AF647 hairpins. Pair 3:Oct*α*2R was detected with AF546 & Oct*β*3R with AF647 hairpins. A split-Gal4 line labeling ER neurons was used to express myr-GFP (see “Flies” section above).

### FISH labeling of TβH

FISH for TβH RNA and associated Confocal imaging were performed as described previously^148^. Probe sequences can be found in the original publication (Probe set 2, Figure 3).

### RNA-Seq

#### RNA-Seq: Expression checks

Neurons of interest were isolated by expressing the fluorescent reporters 20XUAS-IVS-nls-tdTom-BCp10 (attP18);; 10XUAS-unc84-2XGFP (attP2) using split-Gal4 drivers specific for particular cell types and then manually picking the fluorescent neurons from dissociated brain tissue. Each driver/reporter combination was ‘expression checked’ to confirm that the marked cells were sufficiently bright to be sorted effectively and that there was minimal off-target expression in neurons other than those of interest.

#### RNA-Seq: Sorting of labeled neurons

Adult flies were collected when they eclosed, and aged 3-5 days prior to dissection. For each sample, 60-100 brains were dissected in freshly prepared, ice cold Adult Hemolymph Solution (AHS, 108 mM NaCl, 5 mM KCl, 2 mM CaCl2, 8.2 mM MgCl2, 4 mM NaHCO3, 1 mM NaH2PO4, 5 mM HEPES, 6 mM Trehalose, 10 mM Sucrose), and the major tracheal branches removed. The brains were transferred to a 1.5 mL Eppendorf tube containing 500 μL 1 mg/ml Liberase DH (Roche, prepared according to the manufacturer’s recommendation) in AHS, and digested for 1 h at room temperature. The Liberase solution was removed, and the brains washed twice with 800 μl ice cold AHS. The final wash was removed completely and 400 μL of AHS+2% Fetal Bovine Serum (FBS, Sigma) were added. The brain samples were gently triturated with a series of fire-polished, FBS-coated Pasteur pipettes of descending pore sizes until the tissue was homogenized, after which the tube was allowed to stand for 2-3 m so that the larger debris could settle.

For FACS, the samples were triturated in AHS+2%FBS that was run through a 0.2 μm filter, and the cell suspension was passed through a Falcon 5 mL round-bottom tube fitted with a 35 μm cell strainer cap (Fisher). Samples were sorted on a SONY SH800 cell sorter gated for single cells with a fluorescence intensity exceeding that of a non-fluorescent control. Replicates of 15 cells (SS70424 and MB233B, ExR2 lines) and 80 cells (SS00027 and SS50573, EL lines) were purity sorted into 3 μL Lysis Buffer (0.2% Triton X-100, 0.1 U/μL RNAsin) and flash frozen on dry ice.

#### RNA-Seq: Library preparation and sequencing

One microliter of harsh lysis buffer (50 mM Tris pH 8.0, 5 mM EDTA pH 8.0, 10 mM DTT, 1% Tween-20, 1% Triton X-100, 0.1 g/L Proteinase K (Roche), 2.5 mM dNTPs (Takara), and ERCC Mix 1 (Thermo-Fisher) diluted to 1E-7) and 1 μl 10 μM barcoded RT primer were added, and the samples were incubated for 5 m at 50°C to lyse the cells, followed by 20 m at 80°C to inactivate the Proteinase K. Reverse transcription master mix (2 μL 5X RT Buffer (Thermo-Fisher), 2 μL 5M Betaine (Sigma-Aldrich), 0.2 μL 50 μM E5V6NEXT template-switch oligo (Integrated DNA Technologies), 0.1 μL 200 U/μL Maxima H-RT (Thermo-Fisher), 0.1 μL 40U/μL RNAsin (Lucigen), and 0.6 μL nuclease-free water (Thermo-Fisher) was added to the lysis reaction and incubated at 42°C for 1.5 h, followed by 10 m at 75°C to inactivate the reverse transcriptase.

PCR was performed by adding 10 μL 2X HiFi PCR Mix (Kapa Biosystems) and 0.5 μL 60 μM SINGV6 primer and incubating at 98°C for 3 m, followed by 20 cycles of 98°C for 20 s, 64°C for 15 s, 72°C for 4 m, and a final extension of 5 m at 72°C. Groups of 8 reactions were pooled to yield ∼250 μL and purified with Ampure XP Beads (0.6x ratio; Beckman Coulter), washed twice with 75% Ethanol, and eluted in 40 μL nuclease-free water. The DNA concentration of each sample was determined using Qubit High-Sensitivity DNA kit (Thermo-Fisher).

To prepare the Illumina sequencing library, 600 pg cDNA from each pooled sample was used in a modified Nextera XT library preparation (Illumina)^149^ using the P5NEXTPT5 primer and unique i7 primers (Illumina) for SS70424 and MB233B, or Illumina Unique Dual Indexes for SS00027 and SS50573, and extending the tagmentation time to 15 m. The resulting libraries were purified according to the Nextera XT protocol (0.6x ratio) and quantified by qPCR using Kapa Library Quantification (Kapa Biosystems). Six to eight sequencing libraries were loaded on a NextSeq High Output flow cell reading 26 bases in Read 1, including the spacer, sample barcode and UMI, 8 bases in the i5 and i7 index reads, and 50 bases in Read 2 representing the cDNA fragment from the 3′ end of the transcript.

Sequencing adapters were trimmed from the reads with Cutadapt v2.10^150^ prior to alignment with STAR v2.7.5a^151^ to the Drosophila r6.34 genome assembly (Flybase). Sequencing adapters were trimmed and aligned as was done for the bulk RNA-seq data. Gene counts were generated with the STARsolo algorithm using the following additional parameters: “–soloType CB_UMI_Simple–soloCBwhitelist smartscrb_whitelist.txt–soloCBstart 2–soloCBlen 8– soloUMIstart 10–soloUMIlen 10 –soloBarcodeReadLength 0–soloCBmatchWLtype 1MM_multi_pseudocounts–soloStrand Forward–soloFeatures Gene–soloUMIdedup 1MM_All– soloUMIfiltering MultiGeneUMI–soloCellFilter None.” The full set of 384 barcodes designed for this assay was used as the allowlist. Gene counts for the subset of barcodes used in this experiment were extracted using custom R scripts.

## Acknowledgements

Fly stocks obtained from the Bloomington Drosophila Stock Center (NIH P40OD018537) were used in this study. We thank G. Rubin for the pJFRC7-20XUAS-IVS-mCD8::GFP plasmid and G. Rubin and T. Wolff for use of the EL split Gal4 fly lines prior to publication. We acknowledge the FlyLight Project Team at Janelia Research Campus for the use of split Gal4 images. We thank Y. Li for sharing UAS-GRAB-OA1 fly lines and T. R. Clandinin for sharing UAS-GFlamp2 fly lines. We thank A. Shenasa for the preliminary analysis of the EASI-FISH data. Most confocal experiments were conducted at the University of California Berkeley CRL Molecular Imaging Center, RRID:SCR_017852, supported by NSF DBI-1041078. We thank H. Aaron and F. Ives for microscopy support. We thank P. Ramdya, D. Feldman, M. Feller, H. Yang, T. R. Clandinin, M. Silies, Y. Aso, A. Simon, S. Chitnis, C. Sauvola, and members of the Fisher lab for comments on the manuscript. The work by M.E., R.P.R., D.B.T.E., and V.J. was supported by Howard Hughes Medical Institute (HHMI). This work was supported by the NIH (DP2NS132373), NSF (2318081 Y.E.F. Co-PI), a Shurl and Kay Curci Foundation grant, a Seed Grant from the Brain Research Foundation, a Alfred P. Sloan Fellowship,, and a Biohub, San Francisco, Investigator Award to Y.E.F.. M.H.P. is a fellow with the Jane Coffin Childs Memorial Fund for Biomedical Research. Y.E.F. is an HHMI Freeman Hrabowski Scholar.

## Contributions

M.H.P., D.B.T.E, V.J. & Y.E.F. designed the study and interpreted data.

M.H.P. performed and analyzed all 2-photon experiments with assistance from D.B.T.E on the dual imaging data set.

J.C.C. performed immunostaining, confocal imaging, and analysis.

R.P.R. & D.B.T.E. performed RNA-seq experiments and analysis.

M.E. & D.B.T.E. performed EASI-FISH and analysis.

M.H.P. assisted with EASI-FISH and RNA-seq analyses.

A.L. performed the electrophysiology experiments and analysis.

M.H.P. & Y.E.F. generated the HcKCR1 transgenic line.

M.H.P. performed connectomics analysis

M.H.P. & D.B.T.E performed computational modeling

M.H.P. & Y.E.F wrote the manuscript with input from all co-authors

## Code and data availability statement

Custom analysis code is available at: https://github.com/yfisher-lab/plitt_turnerevans_oa_instructs_hd_plasticity/ Data will be made available upon manuscript acceptance.

## Notes

### Competing Interest Statement

The authors have declared no competing interest.

### Summary of Updates

Summary of major changes (New data and analyses): -Knocking out single octopamine receptors that are expressed in ER but not EPG neurons (Oamb or Octα2R) prevents the EPG activity bump from tethering to visual stimuli, replicating the EL>Tβ H-RNAi result (Fig. 4e,f). -EPG bump width narrows in VR with a visual cue relative to darkness in control flies (Fig 1l, replicating (Basnak et al., 2025). Knocking out Oamb or Octα2R abolishes this effect on bump width (Fig 4g,h). -Connectomic analyses highlight the advantage of a presynaptic coincidence detection mechanism. A presynaptic mechanism reduces the potential for synaptic cross-talk compared to a postsynaptic mechanism (Fig 2a-e). -Acutely silencing EPG (using shibireTS) abolishes the EL bump (Extended data Fig. 10i-k). -Additional ER subtypes (ER1 and ER3p) likely express multiple octopamine receptors as measured by EASI-FISH (Extended data Fig 3c-e). -EL-activation drives dramatic cAMP production (GFlamp) in ER but not EPG neurons (Extended data Fig. 8d-e) -Plasticity from EL optogenetic pairing persists for a similar duration as endogenous plasticity (Fig 6e)

## References

1. Petrucco, L. et al. Neural dynamics and architecture of the heading direction circuit in zebrafish. Nat. Neurosci. 26, 765–773 (2023).

2. Ben-Yishay, E. et al. Directional tuning in the hippocampal formation of birds. Curr. Biol. 31, 2592–2602.e4 (2021).

3. Seelig, J. D. & Jayaraman, V. Neural dynamics for landmark orientation and angular path integration. Nature 521, 186–191 (2015).

4. Taube, J. S., Muller, R. U. & Ranck, J. B., Jr. Head-direction cells recorded from the postsubiculum in freely moving rats. I. Description and quantitative analysis. J. Neurosci. 10, 420–435 (1990).

5. Griffiths, B. J. et al. Electrophysiological signatures of veridical head direction in humans. *Nat*. Hum. Behav. 8, 1334–1350 (2024).

6. Hulse, B. K. & Jayaraman, V. Mechanisms underlying the neural computation of head direction. Annu. Rev. Neurosci. 43, 31–54 (2020).

7. Taube, J. S. The head direction signal: origins and sensory-motor integration. Annu. Rev. Neurosci. 30, 181–207 (2007).

8. Knierim, J. J., Kudrimoti, H. S. & McNaughton, B. L. Interactions between idiothetic cues and external landmarks in the control of place cells and head direction cells. J. Neurophysiol. 80, 425–446 (1998).

9. Yoder, R. M., Peck, J. R. & Taube, J. S. Visual landmark information gains control of the head direction signal at the lateral mammillary nuclei. J. Neurosci. 35, 1354–1367 (2015).

10. Ajabi, Z., Keinath, A. T., Wei, X.-X. & Brandon, M. P. Population dynamics of head-direction neurons during drift and reorientation. Nature 615, 892–899 (2023).

11. Peyrache, A., Lacroix, M. M., Petersen, P. C. & Buzsáki, G. Internally organized mechanisms of the head direction sense. Nat. Neurosci. 18, 569–575 (2015).

12. Green, J. et al. A neural circuit architecture for angular integration in Drosophila. Nature 546, 101–106 (2017).

13. Turner-Evans, D. et al. Angular velocity integration in a fly heading circuit. Elife 6, (2017).

14. Skaggs, W. E., Knierim, J. J., Kudrimoti, H. S. & McNaughton, B. L. A model of the neural basis of the rat’s sense of direction. Adv. Neural Inf. Process. Syst. 7, 173–180 (1995).

15. Zhang, K. Representation of spatial orientation by the intrinsic dynamics of the head-direction cell ensemble: a theory. J. Neurosci. 16, 2112–2126 (1996).

16. Kim, S. S., Hermundstad, A. M., Romani, S., Abbott, L. F. & Jayaraman, V. Generation of stable heading representations in diverse visual scenes. Nature 576, 126–131 (2019).

17. Fisher, Y. E., Lu, J., D’Alessandro, I. & Wilson, R. I. Sensorimotor experience remaps visual input to a heading-direction network. Nature 576, 121–125 (2019).

18. Haberkern, H. et al. Maintaining a stable head direction representation in naturalistic visual environments. bioRxiv 2022.05.17.492284 (2022) doi:10.1101/2022.05.17.492284.

19. Hardcastle, B. J. et al. A visual pathway for skylight polarization processing in Drosophila. Elife 10, (2021).

20. Omoto, J. J. et al. Visual input to the Drosophila central complex by developmentally and functionally distinct neuronal populations. Curr. Biol. 27, 1098–1110 (2017).

21. Seelig, J. D. & Jayaraman, V. Feature detection and orientation tuning in the Drosophila central complex. Nature 503, 262–266 (2013).

22. Okubo, T. S., Patella, P., D’Alessandro, I. & Wilson, R. I. A neural network for wind-guided compass navigation. Neuron 107, 924–940.e18 (2020).

23. Garner, D. et al. Connectomic reconstruction predicts visual features used for navigation. Nature 634, 181–190 (2024).

24. Hanesch, U., Fischbach, K.-F. & Heisenberg, M. Neuronal architecture of the central complex in Drosophila melanogaster. Cell Tissue Res. 257, 343–366 (1989).

25. Zhang, Z., Li, X., Guo, J., Li, Y. & Guo, A. Two clusters of GABAergic ellipsoid body neurons modulate olfactory labile memory in Drosophila. J. Neurosci. 33, 5175–5181 (2013).

26. Wolff, T. et al. Cell type-specific driver lines targeting the Drosophila central complex and their use to investigate neuropeptide expression and sleep regulation. Elife 14, (2025).

27. Turner-Evans, D. B. et al. The neuroanatomical ultrastructure and function of a biological ring attractor. Neuron 108, 145–163.e10 (2020).

28. Homberg, U., Kingan, T. G. & Hildebrand, J. G. Immunocytochemistry of GABA in the brain and suboesophageal ganglion of Manduca sexta. Cell Tissue Res. 248, 1–24 (1987).

29. Schäfer, S. & Bicker, G. Distribution of GABA-like immunoreactivity in the brain of the honeybee: GABA-LIKE IMMUNOREACTIVITY IN BEE BRAIN. J. Comp. Neurol. 246, 287– 300 (1986).

30. Scheffer, L. K. et al. A connectome and analysis of the adult Drosophila central brain. Elife 9, (2020).

31. Hulse, B. et al. A connectome of the Drosophila central complex reveals network motifs suitable for flexible navigation and context-dependent action selection. eLife 10, (2020).

32. Cope, A. J., Sabo, C., Vasilaki, E., Barron, A. B. & Marshall, J. A. R. A computational model of the integration of landmarks and motion in the insect central complex. PLoS One 12, e0172325 (2017).

33. Green, J. & Maimon, G. Building a heading signal from anatomically defined neuron types in the Drosophila central complex. Curr. Opin. Neurobiol. 52, 156–164 (2018).

34. Basnak, M. A. et al. Multimodal cue integration and learning in a neural representation of head direction. Nat. Neurosci. 28, 1729–1740 (2025).

35. Dan, C., Hulse, B. K., Kappagantula, R., Jayaraman, V. & Hermundstad, A. M. A neural circuit architecture for rapid learning in goal-directed navigation. Neuron 112, 2581–2599.e23 (2024).

36. Fisher, Y. E. Flexible navigational computations in the Drosophila central complex. Current Opinion in Neurobiology 73, 102514 (2022).

37. Giraldo, Y. M. et al. Sun navigation requires compass neurons in Drosophila. Curr. Biol. 28, 2845–2852.e4 (2018).

38. Fisher, Y. E., Marquis, M., D’Alessandro, I. & Wilson, R. I. Dopamine promotes head direction plasticity during orienting movements. Nature 612, 316–322 (2022).

39. Castillo, P. E., Chiu, C. Q. & Carroll, R. C. Long-term plasticity at inhibitory synapses. Curr. Opin. Neurobiol. 21, 328–338 (2011).

40. Sjöström, P. J., Turrigiano, G. G. & Nelson, S. B. Neocortical LTD via coincident activation of presynaptic NMDA and cannabinoid receptors. Neuron 39, 641–654 (2003).

41. Singla, S., Kreitzer, A. C. & Malenka, R. C. Mechanisms for synapse specificity during striatal long-term depression. J. Neurosci. 27, 5260–5264 (2007).

42. Heifets, B. D., Chevaleyre, V. & Castillo, P. E. Interneuron activity controls endocannabinoid-mediated presynaptic plasticity through calcineurin. Proc. Natl. Acad. Sci. U. S. A. 105, 10250–10255 (2008).

43. Woodin, M. A., Ganguly, K. & Poo, M.-M. Coincident pre– and postsynaptic activity modifies GABAergic synapses by postsynaptic changes in Cl-transporter activity. Neuron 39, 807– 820 (2003).

44. Tong, R., Chater, T. E., Emptage, N. J. & Goda, Y. Heterosynaptic cross-talk of pre– and postsynaptic strengths along segments of dendrites. Cell Rep. 34, 108693 (2021).

45. Harvey, C. D. & Svoboda, K. Locally dynamic synaptic learning rules in pyramidal neuron dendrites. Nature 450, 1195–1200 (2007).

46. Harvey, C. D., Yasuda, R., Zhong, H. & Svoboda, K. The spread of Ras activity triggered by activation of a single dendritic spine. Science 321, 136–140 (2008).

47. Murakoshi, H., Wang, H. & Yasuda, R. Local, persistent activation of Rho GTPases during plasticity of single dendritic spines. Nature 472, 100–104 (2011).

48. Schuman, E. M. & Madison, D. V. Locally distributed synaptic potentiation in the hippocampus. Science 263, 532–536 (1994).

49. McPartland, J., Di Marzo, V., De Petrocellis, L., Mercer, A. & Glass, M. Cannabinoid receptors are absent in insects. J. Comp. Neurol. 436, 423–429 (2001).

50. Vrablik, T. L. & Watts, J. L. Polyunsaturated fatty acid derived signaling in reproduction and development: insights from Caenorhabditis elegans and Drosophila melanogaster: PUFA SIGNALING INWORMS ANDFLIES. Mol. Reprod. Dev. 80, 244–259 (2013).

51. Zhu, B. et al. Drosophila neurotrophins reveal a common mechanism for nervous system formation. PLoS Biol. 6, e284 (2008).

52. Kuntz, S., Poeck, B. & Strauss, R. Visual working memory requires permissive and instructive NO/cGMP signaling at presynapses in the Drosophila central brain. Curr. Biol. 27, 613–623 (2017).

53. Ueno, K. et al. Carbon monoxide, a retrograde messenger generated in postsynaptic mushroom body neurons, evokes noncanonical dopamine release. J. Neurosci. 40, 3533– 3548 (2020).

54. Zatsepina, O. G. et al. Genes responsible for H2S production and metabolism are involved in learning and memory in Drosophila melanogaster. Biomolecules 12, 751 (2022).

55. Schuman, E. M. & Madison, D. V. Nitric oxide and synaptic function. Annu. Rev. Neurosci. 17, 153–183 (1994).

56. Duan, J. et al. Nitric oxide signaling modulates cholinergic synaptic input to projection neurons in Drosophila antennal lobes. Neuroscience 219, 1–9 (2012).

57. Wildemann, B. & Bicker, G. Nitric oxide and cyclic GMP induce vesicle release at Drosophila neuromuscular junction. J. Neurobiol. 39, 337–346 (1999).

58. Hawkins, R. D., Abrams, T. W., Carew, T. J. & Kandel, E. R. A cellular mechanism of classical conditioning in Aplysia: activity-dependent amplification of presynaptic facilitation. Science 219, 400–405 (1983).

59. Waddell, S. Reinforcement signalling in Drosophila; dopamine does it all after all. Current opinion in neurobiology 23, 324–329 (2013).

60. Atwood, B. K., Lovinger, D. M. & Mathur, B. N. Presynaptic long-term depression mediated by Gi/o-coupled receptors. Trends Neurosci. 37, 663–673 (2014).

61. Sinakevitch, I. & Strausfeld, N. J. Comparison of octopamine-like immunoreactivity in the brains of the fruit fly and blow fly. J. Comp. Neurol. 494, 460–475 (2006).

62. Bonanno, S. L. et al. Constitutive and conditional Epitope tagging of endogenous G-protein-coupled receptors in Drosophila. J. Neurosci. 44, e2377232024 (2024).

63. El-Kholy, S. et al. Expression analysis of octopamine and tyramine receptors in Drosophila. Cell Tissue Res. 361, 669–684 (2015).

64. Lee, H.-G., Seong, C.-S., Kim, Y.-C., Davis, R. L. & Han, K.-A. Octopamine receptor OAMB is required for ovulation in Drosophila melanogaster. Dev. Biol. 264, 179–190 (2003).

65. Kim, Y.-C., Lee, H.-G., Lim, J. & Han, K.-A. Appetitive learning requires the alpha1-like octopamine receptor OAMB in the Drosophila mushroom body neurons. J. Neurosci. 33, 1672–1677 (2013).

66. Eddison, M. & Ihrke, G. Expansion-assisted iterative fluorescence in situ hybridization (EASI-FISH) in Drosophila CNS v1. (2022) doi:10.17504/protocols.io.5jyl8jmw7g2w/v1.

67. Sun, F. et al. Next-generation GRAB sensors for monitoring dopaminergic activity in vivo. Nat. Methods 17, 1156–1166 (2020).

68. Lv, M. et al. An octopamine-specific GRAB sensor reveals a monoamine relay circuitry that boosts aversive learning. Natl. Sci. Rev. 11, nwae112 (2024).

69. Aimon, S., Cheng, K. Y., Gjorgjieva, J. & Grunwald Kadow, I. C. Global change in brain state during spontaneous and forced walk in Drosophila is composed of combined activity patterns of different neuron classes. Elife 12, e85202 (2023).

70. Dana, H. et al. High-performance calcium sensors for imaging activity in neuronal populations and microcompartments. Nat. Methods 16, 649–657 (2019).

71. Allen, A. M. et al. A high-resolution atlas of the brain predicts lineage and birth order underly neuronal identity. bioRxiv 2025.06.04.657818 (2025) doi:10.1101/2025.06.04.657818.

72. Ishida, I. G. et al. Neuronal calcium spikes enable vector inversion in the Drosophila brain. Cell 189, 748–764.e25 (2026).

73. Westeinde, E. A. et al. Transforming a head direction signal into a goal-oriented steering command. Nature 626, 819–826 (2024).

74. Handler, A. et al. Distinct dopamine receptor pathways underlie the temporal sensitivity of associative learning. Cell 178, 60–75.e19 (2019).

75. Cohn, R., Morantte, I. & Ruta, V. Coordinated and compartmentalized neuromodulation shapes sensory processing in Drosophila. Cell 163, 1742–1755 (2015).

76. Hige, T., Aso, Y., Modi, M. N., Rubin, G. M. & Turner, G. C. Heterosynaptic plasticity underlies aversive olfactory learning in Drosophila. Neuron 88, 985–998 (2015).

77. Nakagawa, H., Maehara, S., Kume, K., Ohta, H. & Tomita, J. Biological functions of α2-adrenergic-like octopamine receptor in Drosophila melanogaster. Genes Brain Behav. 21, e12807 (2022).

78. Sun, Y. et al. Neural signatures of dynamic stimulus selection in Drosophila. Nat. Neurosci. 20, 1104–1113 (2017).

79. Zhang, Y. et al. Fast and sensitive GCaMP calcium indicators for imaging neural populations. Nature 615, 884–891 (2023).

80. Dana, H. et al. Sensitive red protein calcium indicators for imaging neural activity. Elife 5, e12727 (2016).

81. Kitamoto, T. Conditional modification of behavior in Drosophila by targeted expression of a temperature-sensitive shibire allele in defined neurons. J. Neurobiol. 47, 81–92 (2001).

82. Han, K. A., Millar, N. S. & Davis, R. L. A novel octopamine receptor with preferential expression in Drosophila mushroom bodies. J. Neurosci. 18, 3650–3658 (1998).

83. Balfanz, S., Strünker, T., Frings, S. & Baumann, A. A family of octopamine [corrected] receptors that specifically induce cyclic AMP production or Ca2+ release in Drosophila melanogaster: Drosophilaoctopamine receptors. J. Neurochem. 93, 440–451 (2005).

84. Qi, Y.-X. et al. A new Drosophila octopamine receptor responds to serotonin. Insect Biochem. Mol. Biol. 90, 61–70 (2017).

85. Govorunova, E. G. et al. Kalium channelrhodopsins are natural light-gated potassium channels that mediate optogenetic inhibition. Nat. Neurosci. 25, 967–974 (2022).

86. Yamada, D., Davidson, A. M. & Hige, T. Cyclic nucleotide-induced bidirectional long-term synaptic plasticity in Drosophila mushroom body. J. Physiol. 602, 2019–2045 (2024).

87. Maimon, G. & Abbott, L. F. Functional logic of a cognitive brain system for navigation. Annu. Rev. Neurosci. 49, 411–434 (2026).

88. Maqueira, B., Chatwin, H. & Evans, P. D. Identification and characterization of a novel family of Drosophila beta-adrenergic-like octopamine G-protein coupled receptors: Insect β-adrenergic-like octopamine receptors. J. Neurochem. 94, 547–560 (2005).

89. Lipscombe, D., Kongsamut, S. & Tsien, R. W. Alpha-adrenergic inhibition of sympathetic neurotransmitter release mediated by modulation of N-type calcium-channel gating. Nature 340, 639–642 (1989).

90. Ikeda, S. R. Voltage-dependent modulation of N-type calcium channels by G-protein beta gamma subunits. Nature 380, 255–258 (1996).

91. Herlitze, S. et al. Modulation of Ca2+ channels by G-protein beta gamma subunits. Nature 380, 258–262 (1996).

92. De Waard, M. et al. Direct binding of G-protein betagamma complex to voltage-dependent calcium channels. Nature 385, 446–450 (1997).

93. Livingstone, M. S., Sziber, P. P. & Quinn, W. G. Loss of calcium/calmodulin responsiveness in adenylate cyclase of rutabaga, a Drosophila learning mutant. Cell 37, 205–215 (1984).

94. Levin, L. R. et al. The Drosophila learning and memory gene rutabaga encodes a Ca2+/Calmodulin-responsive adenylyl cyclase. Cell 68, 479–489 (1992).

95. Tomchik, S. M. & Davis, R. L. Dynamics of learning-related cAMP signaling and stimulus integration in the Drosophila olfactory pathway. Neuron 64, 510–521 (2009).

96. Gervasi, N., Tchénio, P. & Preat, T. PKA dynamics in a Drosophila learning center: coincidence detection by rutabaga adenylyl cyclase and spatial regulation by dunce phosphodiesterase. Neuron 65, 516–529 (2010).

97. Chevaleyre, V., Heifets, B. D., Kaeser, P. S., Südhof, T. C. & Castillo, P. E. Endocannabinoid-mediated long-term plasticity requires cAMP/PKA signaling and RIM1alpha. Neuron 54, 801–812 (2007).

98. Burke, C. J. et al. Layered reward signalling through octopamine and dopamine in Drosophila. Nature 492, 433–437 (2012).

99. Cassenaer, S. & Laurent, G. Conditional modulation of spike-timing-dependent plasticity for olfactory learning. Nature 482, 47–52 (2012).

100. Suver, M. P., Mamiya, A. & Dickinson, M. H. Octopamine neurons mediate flight-induced modulation of visual processing in Drosophila. Curr. Biol. 22, 2294–2302 (2012).

101. Crocker, A. & Sehgal, A. Octopamine regulates sleep in drosophila through protein kinase A-dependent mechanisms. J. Neurosci. 28, 9377–9385 (2008).

102. Zhou, C. et al. Molecular genetic analysis of sexual rejection: roles of octopamine and its receptor OAMB in Drosophila courtship conditioning. J. Neurosci. 32, 14281–14287 (2012).

103. Youn, H., Kirkhart, C., Chia, J. & Scott, K. A subset of octopaminergic neurons that promotes feeding initiation in Drosophila melanogaster. PLoS One 13, e0198362 (2018).

104. Watanabe, K. et al. A circuit node that integrates convergent input from neuromodulatory and social behavior-promoting neurons to control aggression in Drosophila. Neuron 95, 1112–1128.e7 (2017).

105. Koon, A. C. et al. Autoregulatory and paracrine control of synaptic and behavioral plasticity by octopaminergic signaling. Nat. Neurosci. 14, 190–199 (2011).

106. Bakshinska, D. et al. Synapse-specific catecholaminergic modulation of neuronal glutamate release. Proc. Natl. Acad. Sci. U. S. A. 122, e2420496121 (2025).

107. Shiozaki, H. M. & Kazama, H. Parallel encoding of recent visual experience and self-motion during navigation in Drosophila. Nat. Neurosci. 20, 1395–1403 (2017).

108. Donlea, J. M. et al. Recurrent circuitry for balancing sleep need and sleep. Neuron 97, 378–389.e4 (2018).

109. Liu, S., Liu, Q., Tabuchi, M. & Wu, M. N. Sleep drive is encoded by neural plastic changes in a dedicated circuit. Cell 165, 1347–1360 (2016).

110. Pimentel, D. et al. Operation of a homeostatic sleep switch. Nature 536, 333–337 (2016).

111. Mitchell, R., Dacke, M. & Webb, B. A robot model of compass cue calibration in the insect brain. bioRxiv 2026.06.25.734539 (2026) doi:10.64898/2026.06.25.734539.

112. Aso, Y. et al. Nitric oxide acts as a cotransmitter in a subset of dopaminergic neurons to diversify memory dynamics. Elife 8, (2019).

113. Gonzalez, K. C. et al. Synaptic basis of feature selectivity in hippocampal neurons. Nature 637, 1152–1160 (2025).

114. Milstein, A. D. et al. Bidirectional synaptic plasticity rapidly modifies hippocampal representations. Elife 10, (2021).

115. Bittner, K. C., Milstein, A. D., Grienberger, C., Romani, S. & Magee, J. C. Behavioral time scale synaptic plasticity underlies CA1 place fields. Science 357, 1033–1036 (2017).

116. Li, Y., Briguglio, J. J., Romani, S. & Magee, J. C. Mechanisms of memory-supporting neuronal dynamics in hippocampal area CA3. Cell 187, 6804–6819.e21 (2024).

117. Feldman, D. E. The spike-timing dependence of plasticity. Neuron 75, 556–571 (2012).

118. Brunelli, M., Castellucci, V. & Kandel, E. R. Synaptic facilitation and behavioral sensitization in Aplysia: possible role of serotonin and cyclic AMP. Science 194, 1178–1181 (1976).

119. Eschbach, C. et al. Recurrent architecture for adaptive regulation of learning in the insect brain. Nat. Neurosci. 23, 544–555 (2020).

120. Aso, Y. & Rubin, G. M. Dopaminergic neurons write and update memories with cell-type-specific rules. Elife 5, e16135 (2016).

121. Tritsch, N. X. & Sabatini, B. L. Dopaminergic modulation of synaptic transmission in cortex and striatum. Neuron 76, 33–50 (2012).

122. Lesch, K.-P. & Waider, J. Serotonin in the modulation of neural plasticity and networks: implications for neurodevelopmental disorders. Neuron 76, 175–191 (2012).

123. Bergles, D. E., Doze, V. A., Madison, D. V. & Smith, S. J. Excitatory actions of norepinephrine on multiple classes of hippocampal CA1 interneurons. J. Neurosci. 16, 572– 585 (1996).

124. Bean, B. P. Neurotransmitter inhibition of neuronal calcium currents by changes in channel voltage dependence. Nature 340, 153–156 (1989).

125. Starke, K. Regulation of noradrenaline release by presynaptic receptor systems. Rev. Physiol. Biochem. Pharmacol. 77, 1–124 (1977).

126. Schultz, W. Dopamine reward prediction error coding. Dialogues Clin. Neurosci. 18, 23– 32 (2016).

127. Lerner, T. N., Holloway, A. L. & Seiler, J. L. Dopamine, updated: reward prediction error and beyond. Current opinion in neurobiology 67, 123–130 (2021).

128. Sara, S. J. & Bouret, S. Orienting and reorienting: the locus coeruleus mediates cognition through arousal. Neuron 76, 130–141 (2012).

129. Jensen, K. T., Hennequin, G. & Mattar, M. G. A recurrent network model of planning explains hippocampal replay and human behavior. Nat. Neurosci. 27, 1340–1348 (2024).

130. Whittington, J. C. R. et al. The Tolman-Eichenbaum machine: Unifying space and relational memory through generalization in the hippocampal formation. Cell 183, 1249–1263.e23 (2020).

131. Chandra, S., Sharma, S., Chaudhuri, R. & Fiete, I. Episodic and associative memory from spatial scaffolds in the hippocampus. Nature 638, 739–751 (2025).

132. Plaza, S. M. et al. neuPrint: An open access tool for EM connectomics. Front. Neuroinform. 16, 896292 (2022).

133. Liu, W., Liu, C., Ren, P.-G., Chu, J. & Wang, L. An improved genetically encoded fluorescent cAMP indicator for sensitive cAMP imaging and fast drug screening. Front. Pharmacol. 13, 902290 (2022).

134. Bates, A. S. et al. Distributed control circuits across a brain-and-cord connectome. Nature 1–14 (2026).

135. Dorkenwald, S. et al. Neuronal wiring diagram of an adult brain. Nature 634, 124–138 (2024).

136. Berg, S. et al. Sexual dimorphism in the complete Drosophila male central nervous system connectome. Cell 189, 5504–5526.e15 (2026).

137. Nern, A., Pfeiffer, B. D. & Rubin, G. M. Optimized tools for multicolor stochastic labeling reveal diverse stereotyped cell arrangements in the fly visual system. Proc. Natl. Acad. Sci. U. S. A. 112, E2967–76 (2015).

138. Klapoetke, N. C. et al. Independent optical excitation of distinct neural populations. Nat. Methods 11, 338–346 (2014).

139. Pfeiffer, B. D. et al. Refinement of tools for targeted gene expression in Drosophila.Genetics 186, 735–755 (2010).

140. Vogt, K. et al. Shared mushroom body circuits underlie visual and olfactory memories in Drosophila. Elife 3, e02395 (2014).

141. Shuai, Y. et al. Driver lines for studying associative learning in Drosophila. Elife 13, (2025).

142. Moore, R. J. D. et al. FicTrac: a visual method for tracking spherical motion and generating fictive animal paths. J. Neurosci. Methods 225, 106–119 (2014).

143. Reiser, M. B. & Dickinson, M. H. A modular display system for insect behavioral neuroscience. J. Neurosci. Methods 167, 127–139 (2008).

144. Sofroniew, N., et al. Napari: A Multi-Dimensional Image Viewer for Python. (Zenodo, 2026). doi:10.5281/ZENODO.22100191.

145. Hagberg, A. A., Schult, D. A. & Swart, P. J. Exploring network structure, dynamics, and function using NetworkX. in Proceedings of the Python in Science Conference 11–15 (SciPy, 2008).

146. Gouwens, N. W. & Wilson, R. I. Signal propagation in Drosophila central neurons. J. Neurosci. 29, 6239–6249 (2009).

147. Close, K., He, Y., Jeter, J., Ihrke, G. & Eddison, M. Multiplex detection of gene expression in the intact Drosophila brain using expansion-assisted iterative fluorescence in situ hybridization. J. Vis. Exp. e67656 (2025).

148. Meissner, G. W. et al. Mapping neurotransmitter identity in the whole-mount Drosophila brain using multiplex high-throughput fluorescence in situ hybridization. Genetics 211, 473– 482 (2019).

149. Soumillon, M., Cacchiarelli, D., Semrau, S., van Oudenaarden, A. & Mikkelsen, T. Characterization of directed differentiation by high-throughput single-cell RNA-Seq. bioRxiv 003236 (2014) doi:10.1101/003236.

150. Martin, M. Cutadapt removes adapter sequences from high-throughput sequencing reads. EMBnet J. 17, 10 (2011).

151. Dobin, A. et al. STAR: ultrafast universal RNA-seq aligner. Bioinformatics 29, 15–21 (2013).

